# Heterogeneity of radial spokes structural components and associated enzymes in *Tetrahymena* cilia

**DOI:** 10.1101/2023.08.11.551218

**Authors:** Marta Bicka, Avrin Ghanaeian, Corbin Black, Ewa Joachimiak, Anna Osinka, Sumita Majhi, Anna Konopka, Ewa Bulska, Khanh Huy Bui, Dorota Wloga

**Author notes:** equal contribution (Marta Bicka, Avrin Ghanaeian, Corbin Black).

## Abstract

Radial spokes, RS1-RS2-RS3, are T-shaped, multiprotein complexes that transmit regulatory signals from central apparatus to outer doublet dyneins. RSs, especially RS3, differ in morphology, protein composition, and RS base-docked IDAs. Spokes defects alter cilia beating frequency, waveform, and amplitude leading, in humans, to primary ciliary dyskinesia and infertility. In contrast to RS1 and RS2, the protein composition of RS3 is partly resolved. Moreover, the role of particular spokes is unclear. Ciliate *Tetrahymena thermophila* has three Rsp3 paralogs and two or three paralogs of some other RSPs. Using multiple complementary approaches, we showed that *Tetrahymena* forms RS1 and RS2 subtypes having core composed of various Rsp3 paralogs and one type of Rsp3-less RS3. We elucidated proteomes of RS subtypes and identified novel RS-associated proteins, including enzymatic proteins involved in local regulation of the ADP/ATP levels and protein phosphorylation, whose presence further diversifies RSs properties and likely functions.

**In brief:** Radial spokes differ in their protein composition and architecture. Studies in a ciliate *Tetrahymena* revealed Rsp3 paralogs-dependent RS1 and RS2 subtypes, Rsp3-less RS3, and diversity of the RSP3 mutants’ phenotype. Known RS components and newly identified structural and enzymatic proteins were assigned to particular RSs.

## Introduction

Motile cilia and homologous flagella are highly evolutionarily conserved, hair-like cell protrusions, supported by a microtubular skeleton, the axoneme, composed of nine peripheral doublets and two central singlets. The central microtubules, C1 and C2, together with attached complexes, the projections, form the central apparatus (CA), a structure believed to initiate signals required for planar cilia beating. It is proposed that transient interactions between CA projections and outer doublet-docked radial spokes (RSs) enable transmission of regulatory signals to outer doublet complexes, including nexin-dynein regulatory complex (N-DRC) and motor protein-containing outer and inner dynein arms (ODAs and IDAs) ^1–5^.

RSs are T-shaped, approximately 40-nm-long complexes with three morphologically distinct regions: (i) an elongated stalk docking the entire RS structure to the outer doublets and having some conformational flexibility enabling RS, especially RS1, to tilt ^5,6^, (ii) an orthogonal head transiently coming in contact with the CA projections, and (iii) a neck connecting a stalk and a head. Within the 96-nm axonemal repeat unit, RSs are arranged as triplets (RS1, RS2, RS3). The distance between the neighboring spokes is specific to each spoke pair, enabling their identification^7,8^.

Similar as ODA and IDAf, RSs preassemble within the cells body forming an “L”-shaped, 12S complexes composed of RSPs 1-7 and 9-12. The difference between the calculated and estimated molecular weight of the 12S complex suggests a presence of RSP dimers or some additional proteins ^9,10^. Except the very proximal part of the cilium, the transport of 12S precursor within cilia is mediated by IFT and ARMC2 adaptor protein ^11,12^. The 12S complexes are converted in cilium into 20S mature T-shaped complexes ^9,10,13^. but RSPs can also be added onto preexisting spokes to repair them ^14^.

Ultrastructural studies of cilia and flagella from evolutionarily distant species clearly demonstrate that RS1-RS2-RS3 spokes, initially recognized as morphologically similar structures ^7^ differ in their architecture and protein composition. The most striking example of RSs diversity are spokes in *Chlamydomonas*, where RS1 and RS2 are full length structures while RS3 is short, knob-like ^15–17^, and similar to the base part of the full-length RS3 ( ^17^supplementary Fig 2). Even in species having all full-length RSs ^18^, the RS3 differs from RS1 and RS2. Moreover, some small differences also exist between RS1 and RS2 architecture ^5,19–27^.

Besides differences between spokes within the 96-nm axonemal unit, the architecture of particular spokes varies in different species. In choanoflagellate *S. rosetta* ^28^ and metazoan, sea urchin ^17,29^, *Danio rerio* ^30^, mice, and humans ^23,31^, the RS1 and RS2 heads are symmetrical, H-shaped structures composed of two parallel halves divided by a deep cleft. In contrast, in *Chlamydomonas*, *Tetrahymena*, and *Trypanosoma* the RS1 and RS2 heads are nearly flat, with less pronounced cleft and large lobs extending their area ^15–17,21,29,32^. Also the morphology of RS3 head differs between *Tetrahymena* and metazoans ^29^. Additionally, in *P. rosetta* and studied metazoans, the RS heads are not connected or a thin connection exist between some RS2 and RS3 ^23,29–31^, while in *Tetrahymena*, *Chlamydomonas,* and *Trypanosoma* all RS heads are firmly connected ^15–17,21,29^. Finally, in mice and humans, RSs in sperm flagella and cilia of multiciliated cells also diverged. In sperm cell flagellum, RSs are accompanied by additional structures not observed in cilia of multiciliated cells: the barrel-shaped density near RS1, the RS2-RS3 cross-linker, and a density near RS3 base (RS3 scaffolds) ^31^. Moreover, while RS head subunits, Rsph1 and Rsph4a, are expressed in multiciliated cells, in sperm flagellum, the RSs contain Rsph6a, a paralog of Rsp4a ^33^ and Rsph10 ^34–37^.

In *Chlamydomonas*, the RSP3 dimer, a core component of RSs ^38^, stretches from the RS base (N-termini) to the RS head (C-termini) ^2^ and directly interacts with most of RSPs ^19^. Pioneering studies using the *Chlamydomonas pf14* mutant carrying a mutation in the *RSP3* gene ^39^ and lacking full-length spokes, RS1 and RS2, led to the identification of RS subunits, RSP1-RSP23. Next, cryo-EM studies combined with proteomics, and molecular modeling ^1,19,32,40,41^ yielded a recent atomic model of RS1-RS2 protein composition in this green alga and mice ^4,5,20,27^. However, the RS3 composition is still not fully resolved. The knob-like RS3 is maintained in *Chlamydomonas pf14* mutant ^15,17^ but missing in *fap61* (CaM-IP3) and *fap91* (CaM-IP2) mutants ^42^. In *Tetrahymena*, knockout of *CFAP91* affects RS3 and RS2 base ^25^ while deletion of *CFAP61* and *CFAP251* affects part of RS3 stalk or an arch-like structure at the RS3 base, respectively ^24^. More recently, Lrrc23 was shown to be a component of RS3 head in mice sperm flagellum ^22^. Strikingly, in mice and humans, mutations in *CFAP61*, *CFAP91*, *CFAP251,* and *LRRC23* cause male infertility but not primary ciliary dyskinesia or hydrocephalus, suggesting more differences in the RS3 structure and/or function between sperm flagellum and cilia in multiciliated cells ^43–47^. To date, Cfap61, Cfap251, Lrrc23, and Cfap91 are the only known RS3 components. This raises a question about the remaining subunits of RS3 spoke.

To address this issue, we took advantage of existing *Tetrahymena* knockout mutants with RS(s) defects (CFAP61-KO ^24^, CFAP206-KO ^48^, and CFAP91-KO ^25^, and newly engineered mutants lacking Rsp3 paralogs to compare wild-type and mutant ciliomes using label-free and tandem mass tag (TMT) quantitative mass spectrometry and decipher RSs proteomes. To verify if candidate RSPs (proteins diminished in mutant ciliomes) are indeed present in RSs or their vicinity, we performed co-IP and BioID assays. These proteomic approaches combined with cryo-electron tomography (cryo-ET) studies of ultrastructural alterations in RS mutants and cross-links data ^49^ allowed us to assign with a high probability the substantial number of known RSPs and newly identified candidate proteins to either RS1, RS2, or RS3. Importantly, among newly identified proteins, we found not only structural proteins but also proteins having predicted enzymatic folds and thus likely having an enzymatic activity (further shortly enzymes), including serine-threonine kinases and adenylate kinases that locally controls ADP/ATP levels. We also observed a destabilization of some single-headed inner dynein arms in the analyzed RS mutants and obtained evidence suggesting interactions of RS with specific central apparatus components.

Here, we propose a model of *Tetrahymena* RS protein composition, showing both ubiquitous and RS-specific RSPs, new candidates of RS structural proteins, and RS-associated enzymes. We postulate that *Tetrahymena,* has sub-types of RS1 and RS2 spokes that together with different associated enzymes further increase the heterogeneity of RSs and likely their function. Our data, besides expanding the general knowledge regarding RSs composition, can contribute to a better understanding of the molecular mechanisms regulating cilia motion.

## Results

### Deletion of *Tetrahymena* Rsp3 paralogs differently affects cilia beating

In contrast to *Chlamydomonas* and animals, ciliates have more than one RSP3 ortholog. *Tetrahymena* genome encodes three RSP3 orthologs, Rsp3A, Rsp3B, and Rsp3C (Figure S1). Rsp3B is most similar to *Chlamydomonas* and metazoan RSP3 within the axoneme targeting domain ^50^ and the radial spoke 3 domain ^51^ (39% identity and 63% similarity to human RSPH3). Based on the amino acids’ conservation, *Tetrahymena* Rsp3 paralogs have an AKAP domain scaffolding the cAMP-dependent protein kinase holoenzyme ^52^ and amphipathic helices, AH-R, binding RIIa domains, and AH-D, binding DPY-30 domains ^53^ (Figure S1). Three LC8-interacting TQT-like motifs, common in N-termini of *Chlamydomonas* and vertebrates RSP3^54^, are predicted only in Rsp3B while Rsp3A and Rsp3C have one and two TQT-like motifs, respectively. Based on the consensus of the LC8 recognition motifs ^55,56^, other LC8 binding sites in *Tetrahymena* Rsp3 paralogs are unlikely. Interestingly, the Rsp3C is unusual in having predicted N-terminal ARF (ADP-ribosylation factor) domain and a C-terminal tail enriched in glutamic acid residues. In silico, the Rsp3C-like RSP3 orthologs were found only in ciliates closely related to *Tetrahymena* such as *Paramecium*, *Ichthyophthirius*, and *Pseudocohnilembus* but not in *Stentor* or *Stylonychia* species.

Ciliary Rsp3A, Rsp3B, and Rsp3C have several isoforms, suggesting their posttranslational modifications, likely including most common, phosphorylation (Figure S2). In human RSPH3, two threonine residues, T243 and T286, are phosphorylated by ERK1/2 ^52^. In corresponding positions, the RSP3 orthologs have either threonine or serine residues (Figure S1) except *Tetrahymena* Rsp3A, missing first, and Rsp3B, missing a second putative phosphorylation site.

*Tetrahymena* Rsp3 paralogs expressed as fusions with a C-terminal 3xHA tag under the control of transcriptional promoter, localize along the entire cilia length except the ciliary tip (Figure 1A-C). When co-expressed, 3xHA and GFP-tagged Rsp3 paralogs were incorporated concurrently into growing cilia (Figure S3) but some minor fluctuation of Rsp3C-GFP signal was noticeable, in growing and full-length cilia (Figure 1 D-E). To explore the significance of Rsp3 paralogs, we constructed knockout strains (Figure S4). Deletion of a single *RSP3* gene affected cells swimming and cilia beating parameters (Figure 2) and in the case of RSP3B-KO, reduced cilia length to ∼84% of the wild-type cilia (Figure S5A). The loss of Rsp3B had the most profound effect on cell motility and reduced the swimming speed to ∼12% of the wild-type, while deletions of either *RSP3A* or *RSP3C* were less damaging (70% and 56% of the wild-type speed, respectively). *Tetrahymena* cells are propelled by the forces generated by cilia beating in metachronal waves. The cilium beating cycle has two phases, the power and recovery strokes that occur in different planes in relation to the cell surface, perpendicular and parallel, respectively. During the power stroke the cilium is basically straight while during the recovery stroke it bends, and the bend position shifts from the cilium base to the tip as the recovery stroke progresses (Figure 2F, Video S1) ^25,57^. Deletion of *RSP3A* or *RSP3C* did not change the ciliary waveform and beat amplitude but slightly affected cilia metachrony and beating frequency (Figure 2F-H, Figure S5B-E and Video S2, Video S4). In contrast, cilia lacking Rsp3B were beating in a less coordinated manner, with significantly lower frequency, inconsistencies in the waveform and amplitude between neighboring cilia and between subsequent beating cycles performed by the same cilium, exhibiting periods of nearly normal and highly modified pattern. Moreover, the distal end of the majority of RSP3B-KO cilia was still bending during the power stroke (Figure 2F, Figure S5B-E Video S3). A delayed straightening of the very distal end was also sometimes observed in cells lacking Rsp3C.

**Figure 1.**
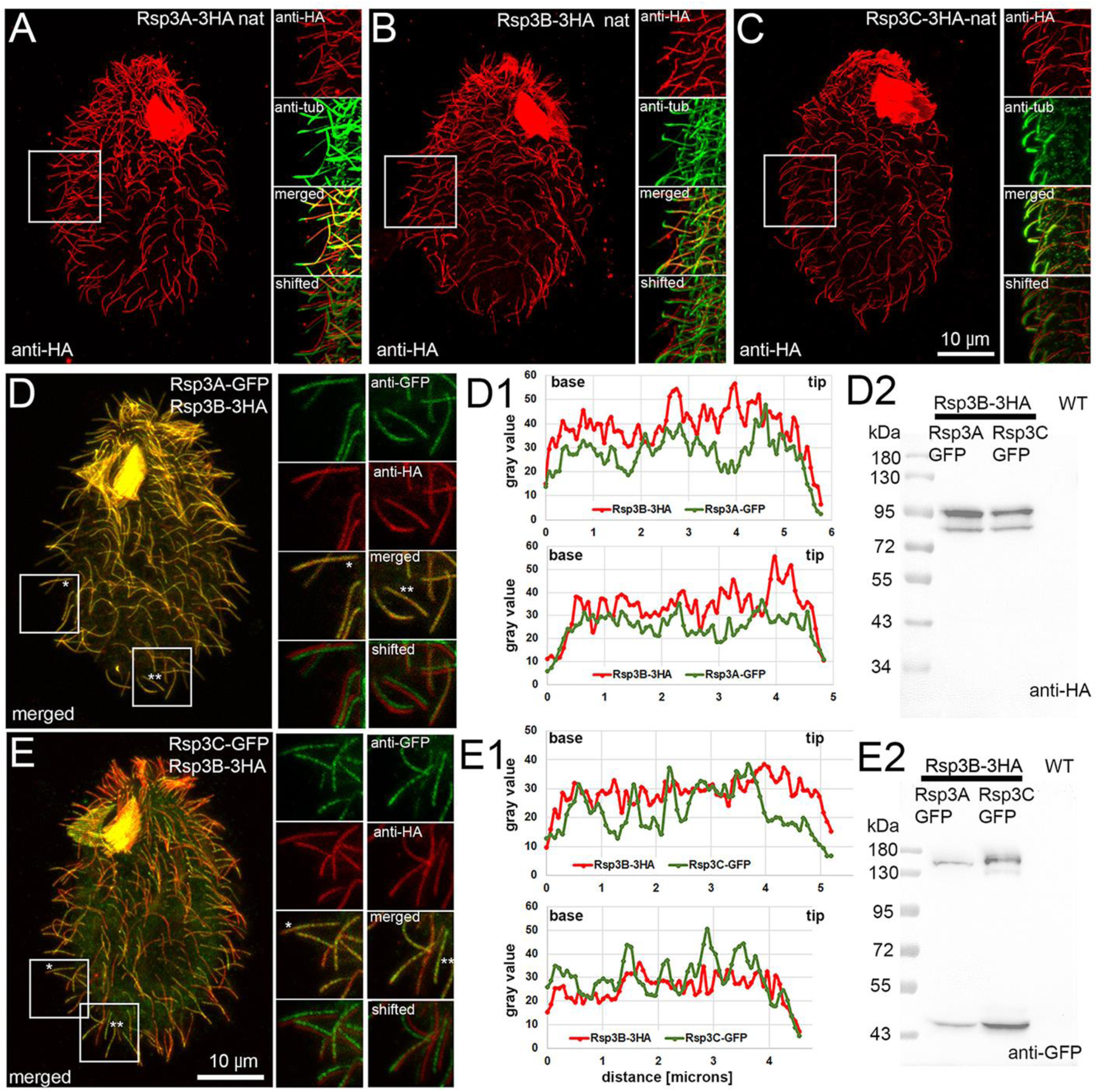
Ciliary localization of Rsp3 paralogs. (A-C) 3HA-tagged Rsp3 paralogs expressed under the control of the transcriptional promoters localize along the entire cilia length except for the ciliary tip. Cells were co-labeled with (A-B) anti-acetylated tubulin antibody, or (C) polyG to visualize ciliary shaft. To the right, the magnified fragments of the cells as indicated by the white insets, stained with anti-HA (red) and anti-tubulin (green) antibodies, and merged images (red and green). Below are images showing both channels but with some shifts to better visualize the presence of the Rsp3 paralogs along the entire cilia. (D-E2) Co-localization of Rsp3B-3HA and (D) GFP-tagged Rsp3A or (E) Rsp3C. To the right, the magnified fragments of the cells as in (A-C). (D1-E1) Plots representing the intensity of the 3HA (red) and GFP (green) fluorescence signal in exemplary cilia marked by stars in (D and E) and enlarged insets. (D2-E2) Western blot analyses of co-expressed Rsp3 paralogs.

**Figure 2.**
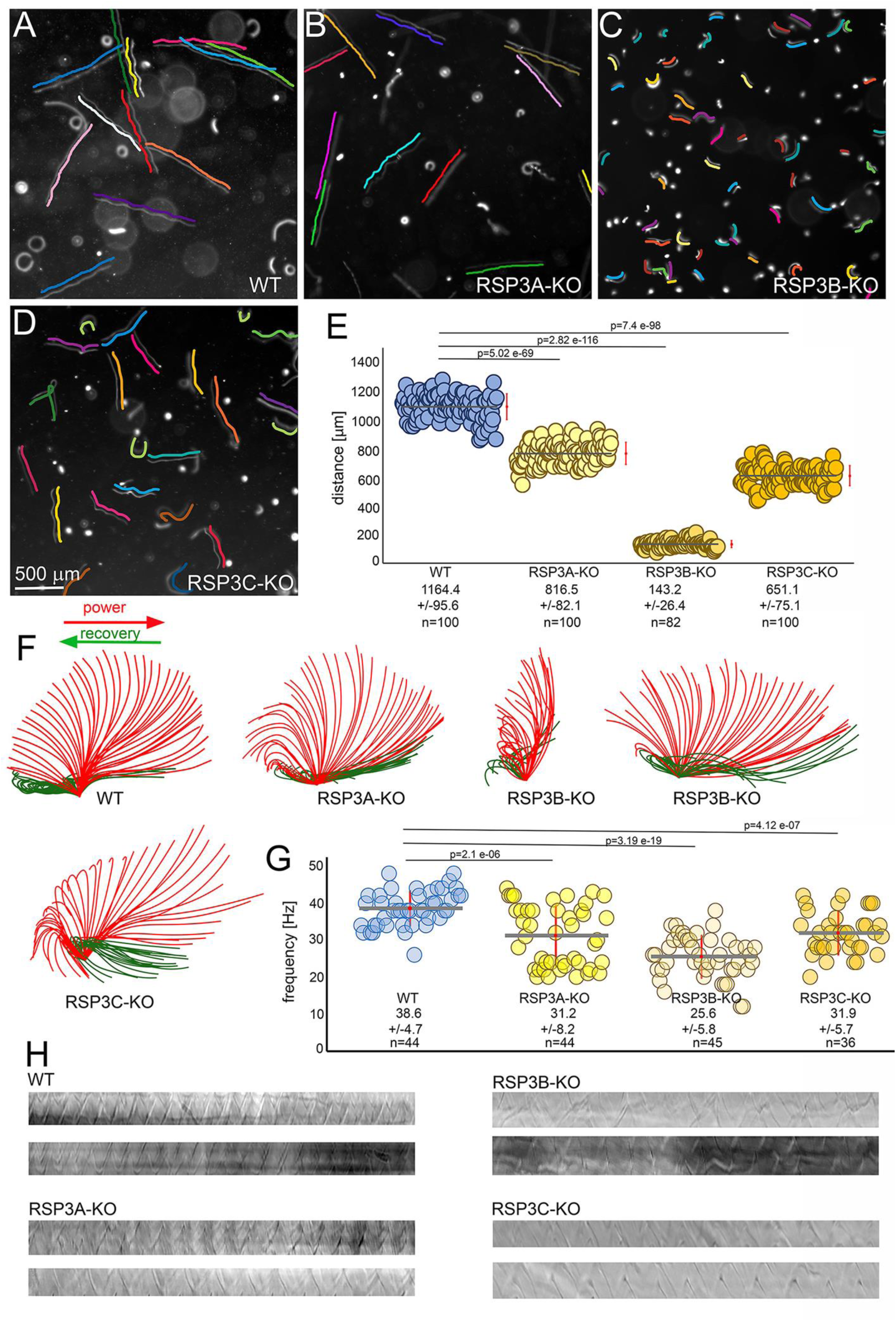
Deletion of each *RSP3* gene differently affects cilia beating. (A-D) WT (A), RSP3A-KO (B), RSP3B-KO (C), and RSP3C-KO (D) trajectories recorded for 3 sec using a high-speed video camera. The cell swimming paths are indicated by the parallel colored lines. Bar = 500 µm. (E) A comparison of the length of trajectories recorded for 3 sec. The red bar position to the right represents the standard deviation. The wild-type cells swam on average 388+/-32 µm/s (n= 100), and the swimming speeds of the RSP3A-KO, RSP3B-KO, and RSP3C-KO mutants were reduced to 272+/-27 µm/s (n=100), 48+/-9 µm/s (n=82), and 217+/-25 µm/s (n=101), respectively. Statistical significance was calculated using student t-test. (F) Drawings showing examples of the most frequently observed subsequent positions of a cilium of WT, *RSP3A*, and *RSP3C* knockout cells and most extreme difference in cilia beating in RSP3B-KO cells. The ciliary waveform and amplitude in the RSP3B-KO mutant can vary both in the case of neighboring cilia and during subsequent cycles of the same cilium. (G) Graph showing a range of cilia beating frequencies in wild-type and *RSP3* knockout mutants. The red bar represents the standard deviation. Statistical significance was calculated using student t-test. (H) Examples of the kymographs used to calculate cilia beating frequency.

### RSP3 paralogs and heterogeneity of RS1 and RS2 spoke structure

The differences in the phenotype of *RSP3* knockouts suggest that lack of the particular paralog affects different RS types and/or numbers. The cryo-ET subtomogram averaging of the three RSP3-KO mutants revealed paralog-specific ultrastructural defects (Figure 3, Table S1). Averages of all 96-nm subunits show that overall, RSP3A-KO and RSP3C have only the base part of RS1 and RS2 respectively and RSP3B-KO lacks the complete RS2 and the head part of RS1 (Figure 3A). Interestingly, in all *RSP3* mutants, the RS3 was intact, strongly suggesting that none of the Rsp3 paralogs is a RS3 component.

**Figure 3.**
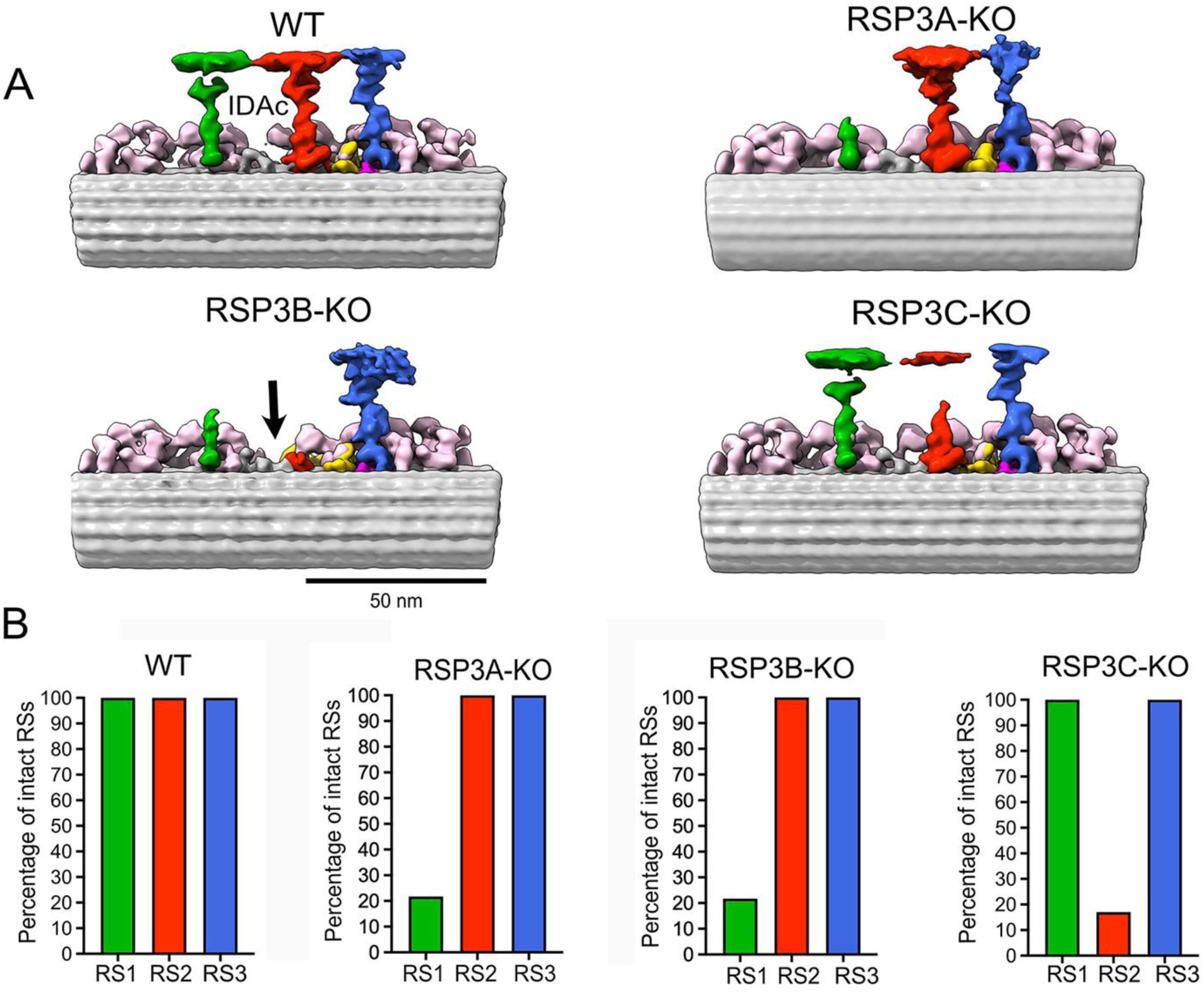
Cryo-ET and subtomogram averaging analyses of the ciliary ultrastructure of wild-type cells and *RSP3* knockout mutants. (A) Subtomogram averages of WT, RSP3A-KO, RSP3B-KO and RSP3C-KO mutants. RS1: green; RS2: red; RS3: blue. Inner dynein arms (IDAs): light pink, N-DRC: yellow, Ccdc96-Cccdc113 complex: dark pink. The black arrow in the RSP3B-KO subtomogram points to the place corresponding to the position of IDAc in the WT cilia. (B) Quantification of intact RSs in WT and RSP3 mutant cells. Each graph shows the percentage of intact RSs. The analysis was performed using subtomogram classification, revealing structural differences in RS integrity.

Existing densities at the bases of the RS1 and RS2 in the averaged maps suggest that there are heterogeneities in RS distribution in those KO mutants. Therefore, we performed image classification of the base and head of the missing RSs to reveal their patterns (Figure 3B, Figure S6 and S7). Our classification reveals that while RSP3A-KO lacks RS1 in most of the 96-nm, 22% of the RS1 looks intact in RSP3A-KO. In RSP3C-KO mutant, 19% of 96-nm repeating units lack RS2 completely while 17% retain the entire RS2, and 58% still contain a long stump of RS2. In RSP3B-KO, 100% of the 96-nm repeat lack RS2 completely while 22% of them have intact RS1.

Thus, Rsp3A is a component of the majority of RS1 spokes and Rsp3C is a component of RS2 spokes. In contrast, Rsp3B is likely a main core protein in the majority of RS1 and all RS2 spokes. The absence of Rsp3C affected only a fraction of RS2 spokes. To evaluate the distribution of the intact RS2 along the doublet microtubule (DMT) in RSP3C-KO, we performed a statistical analysis of the classes in which RS2 remains intact in RSP3C-KO subtomograms. We found that there was no significant difference in the distribution of particles between the proximal and medial-distal regions of each doublet within the axoneme (Figure 4).

**Figure 4.**
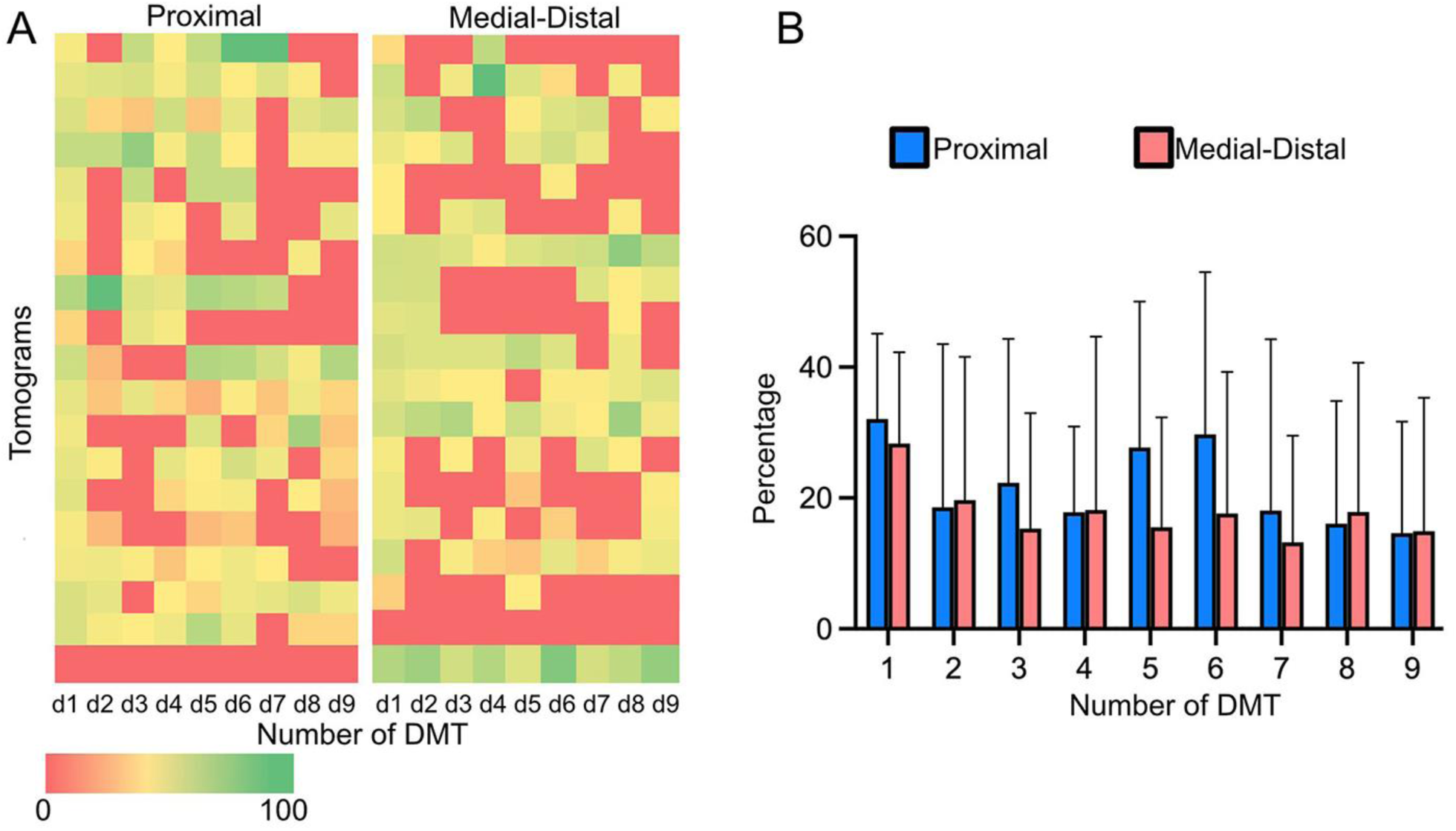
Ultrastructural analyses of Rsp3C-containing RS. (A-B) Distribution of intact RS2 in the RSP3C-KO axoneme. (A) Heat map showing the distribution of particles within each doublet at the proximal and medial-distal ends of the axoneme. The doublet with the highest particle count is considered as the first doublet in the heat map. To assess the distribution of subtomograms with intact RS2 across the doublets, counts were normalized; “0” indicates that none of the subtomograms have intact RS2, “100” that particular doublet, have all RS2 intact. (B) Graph representing mean percentage of intact RS2 subtomograms for a specific doublet. Tomograms were treated as biological replicates, thereby collecting biological variability within each doublet. Error bars (standard deviation) indicate the variability in the data across tomograms.

To study the extend of RS defects, we isolated cilia from wild-type and *RSP3-KO* cells and performed comparative proteomic mass spectrometry analyses using label-free (LFQ) and TMT 10-plex isobaric mass tagging approaches. The experiments were repeated three times to generate independent sets of data. At least 75% of proteins were present in all three replicas. Approximately one-thousand ciliary proteins from wild-type cells and *RSP* mutants were identified in LFQ experiments, while nearly four times more were detected using the TMT approach. Based on principal component analysis of obtained ciliomes using Ingenuity Pathway Analysis, the RSP3A-KO ciliome is most similar to that of a wild-type (Figure S8). Changes in the protein levels were analyzed using Perseus software (v. 2.0.3) ^58^. The t-test (p <0.05) was used to identify proteins whose level significantly changed (a ±1.5-fold) in mutants’ versus that in wild-type cilia. The obtained data are graphically represented as Volcano plots -log (*P* value) *vs.* log_2_ (fold change of mutant/WT, Figure S8) and Venn diagrams (Figure S9A-C). If there were discrepancies between LFQ and TMT data, we interpret changes in the protein level according to the latter analyses.

Global mass spectrometry analyses of the RSP3A-KO, RSP3B-KO, and RSP3C-KO cilia confirmed that the protein encoded by the targeted gene was completely eliminated in respective mutants (Table S2). The knockout of *RSP3A* or *RSP3C* did not significantly affect the levels of other Rsp3 paralogs while in RSP3B-KO cilia the level of Rsp3A was diminished, suggesting that Rsp3A-Rsp3B could form a heterodimer or that in the absence of RS2, some RS1 are unstable. Of note, in none of the *RSP3* mutants, the level of remaining Rsp3 paralogs was significantly elevated, suggesting that substitution of missing paralog by another one if occurs is a minor phenomenon.

The RSP3 is present in axonemal RS as a dimer ^38^. Based on cryo-ET and proteomic data we propose that *Tetrahymena* cilia could have up to three RS1 subtypes with a core composed of: (i) Rsp3A dimer (RS1 missing in RSP3A-KO but intact in RSP3B-KO cilia), (ii) Rsp3B dimer (RS1 missing in RSP3B-KO but present in RSP3A-KO cilia), and (iii) RS1 containing Rsp3A-Rsp3B heterodimer. The existence of this last class is supported by proteomic data showing that the level of Rsp3A was substantially reduced in *RSP3B* mutant cilia. On the other hand, in *Tetrahymena,* the RSs’ heads are connected and therefore the RSs may stabilize one another. Thus, some Rsp3A-containing RS1 spokes could be destabilized (secondary effect) in the RSP3B-KO mutant, leading to a moderate reduction of the Rsp3A level. To address this issue, we search data from cross-link experiments of *Tetrahymena* cilia and found a highly confident cross-link between Rsp3A and Rsp3B (RSP3A K172 to RSP3B K195) suggesting a direct contact ^49^. We also engineered cells expressing Rsp3A-BCCP under the control of the *RSP3A* promoter and analyze BCCP localization using cryo-ET and subtomogram averaging. A comparison of the subtomogram-averaged map from the WT and Rsp3A-BCCP samples revealed an additional density on the IDA-facing side of the RS1 head (Figure 5A, B). Moreover, when the entire RS1 model from *Chlamydomonas* was fitted into our maps (Figure 5C), the extra globular density corresponding to streptavidin was found near the C-terminus of only one copy of the *Chlamydomonas’s* RSP3 (Figure 5D). Taken together, the above data suggest that some of RS1 spokes could indeed have a core composed of Rsp3A-Rsp3B heterodimers and thus it is possible that in *Tetrahymena* there are three types of RS1, depending on the Rsp3 dimer composition.

**Figure 5.**
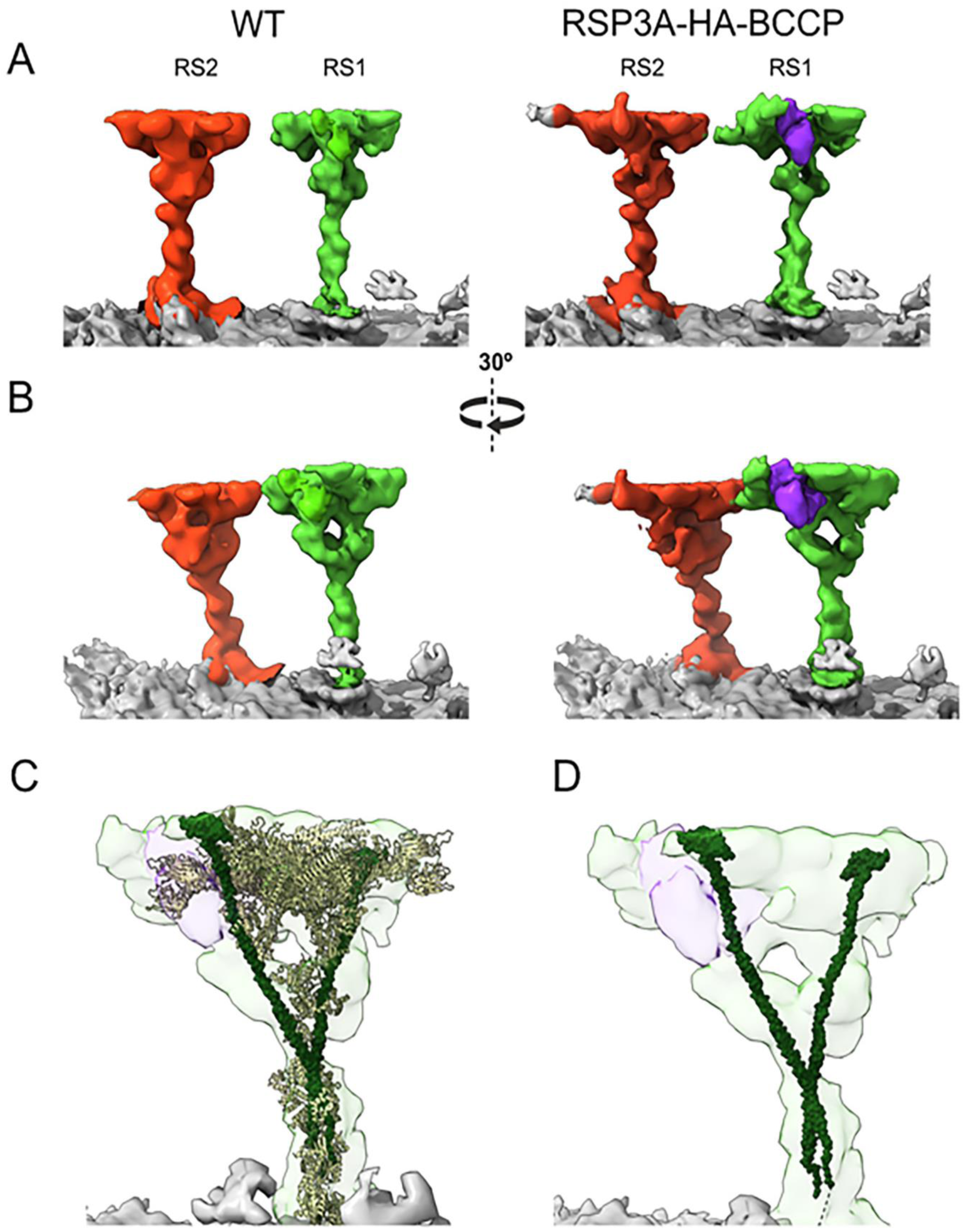
Subtomogram structure of the outer doublet in wild-type and Rsp3A-HA-BCCP expressing cells. (A) Locally refined structures of RS1 and RS2 in the WT and BCCP-tagged axoneme. RS1 is shown in green, RS2 in red, and streptavidin in purple. (B) View of the DMT with a 40-degree rotation. (C) The model of RS1 from *Chlamydomonas reinhardtii*, fitted into the RS1 density from the Rsp3A-HA-BCCP map. (D) The complete RS1 model from *Chlamydomonas reinhardtii*, with all proteins hidden except RSP3 proteins.

The absence of Rsp3C affected the majority of RS2 but only in ∼19% of the analyzed subtomograms, the entire RS2 was missing, which is in striking contrast with RSP3B knockout in which all RS2 were eliminated. Thus, a presence of RS2 spokes containing Rsp3C homodimer is unlikely or very infrequent. We propose that Rsp3B is a component of all *Tetrahymena* RS2 spokes and that RS2 affected in RSP3C-KO mutant, could contain either Rsp3B-Rsp3C heterodimer or Rsp3B homodimer additionally stabilized by Rsp3C. The unaltered level of Rsp3B in *RSP3C* knockout suggests that in the absence of Rsp3C, the Rsp3B-containing partly collapsed RS2-remnants were still attached to the axoneme but no longer maintained their T-shape structure (Figure S6, RS2 head masked). Thus, likely, Rsp3C is present in a subset of RS2 spokes. Surprisingly, the level of Rsp3C in RSP3B-KO cilia was also unaltered, suggesting that perhaps Rsp3C docks to the axoneme independently of Rsp3B. We did not found data supporting the cross-link between Rsp3B and Rsp3C, however, such result does not exclude that Rsp3B and Rsp3C can co-assemble. To summarize, in contrast to RSs from other studies species, in *Tetrahymena* there are subtypes of RS1 and RS2 spokes, increasing spokes heterogeneity. Based on the available data, the significance of this phenomenon remains unknown.

### Identification of *Tetrahymena* RSP orthologs and protein composition of RSs

The RSP3 dimer interacts with other RSPs, together forming the RS head, neck, and stalk ^10,15,19,41^. In bioinformatics search of *Tetrahymena* RSP orthologs (Figure S10, Table S3) we found that besides three Rsp3 paralogs, *Tetrahymena* genome encodes three paralogs of Rsp4/6 and two of Rsp7, Rsp12, Rsp16, and Cfap198 while other RS proteins are encoded by a single gene. The majority of *Tetrahymena* RS proteins are well evolutionarily conserved (Table S3) and their identification was straightforward, but finding true orthologs of Rsp1, 5, 7, 10, and 12 required additional data. Therefore, we engineered *Tetrahymena* cells expressing RS proteins in fusion with either C-terminal 3HA or HA-BirA* tags under the control of the transcriptional promoter and performed co-IP or BioID assays, respectively, followed by mass spectrometry analyses to identify ciliary proteins that either directly or indirectly interact with 3HA-tagged Rsp3A, Rsp4A, RSP4B, or Rsp4C (Table S4), or are in the vicinity of Rsp3 or Rsp4 paralogs, Cfap61, Cfap91, or Cfap206 (Figure S11, Table S5). The combined in silico and biochemical data together with RS mutant ciliome analyses (see below) enabled identification of the remaining less conserved RS proteins (Table S6).

Rsp1 and Rsp10 are MORN-domain containing proteins. *Tetrahymena* genome encodes 129 proteins having MORN domain ^59^, numerous with similarity to *Chlamydomonas* RSP1 and RSP10. However, only two of such MORN proteins co-precipitated with Rsp4-3HA and were identified in BioID assays. A search of the predicted *Tetrahymena* proteome with *Chlamydomonas* RSP5 aldo-keto reductase as a query, failed to identify an obvious ortholog. Thirteen proteins annotated in the Tetrahymena Genome Database (TGD) as aldo-keto reductases were present in our ciliomes, of which four were significantly reduced in the *RSP3* knockouts. However, neither of these proteins was detected in co-IP or BioID assays (Table S4 and S5). Thus, similar as mammals ^41^, *Tetrahymena* seems to lack RSP5 ortholog. Interestingly, search of the NCBI protein database using *Chlamydomonas* RSP5 as a query, identified RSP5 orthologs only in *Chlamydomonas*-related algae. *Chlamydomonas* RSP7 has an RIIa domain (dimerization-anchoring domain of cAMP-dependent PK regulatory subunit) at its N-terminus and several EF hand motifs in the middle and C-terminal regions. A search of the *Tetrahymena* proteome led to the identification of two proteins, Rsp7A and Rsp7B, both with limited similarity to *Chlamydomonas* RSP7 (Figure S10). The Rsp7A is a 39 kDa protein containing N-terminal RIIa and C-terminal IQ calmodulin-binding motif while 66 kDa Rsp7B has predicted AKA28 (A-kinase anchoring protein) domain and two EF hand motifs. RSP12 is a peptidyl-prolyl cis-trans isomerase (PPIase) ^41^. Out of 15 *Tetrahymena* genome-encoded PPIases, seven were present in cilia, but only two, named here Rsp12A and Rsp12B, were significantly reduced in RS mutants.

To more accurately assign the RS proteins to the particular spokes, we analyzed ciliomes of previously described *Tetrahymena* RS mutants (Figure S9D-E): (i) CFAP206-KO, missing either the entire RS2 or only its base, rarely accompanied by minor RS3 defects ^48^, (ii) CFAP61-KO, lacking a part of the RS3 stalk ^24^, and (iii) CFAP91-KO, missing RS3 and a part of the RS2 base ^25^. Proteins whose level was reduced in CFAP91-KO cilia were assigned as RS3 subunits if their level was unchanged in RS2 mutants (RSP3B-KO, RSP3C-KO, CFAP206-KO) or as RS2 components if were diminished in RS2 mutants. Of note, when only a fraction of spokes is affected in a mutant, the reduction of the protein could not be statistically significant. Furthermore, because in *Tetrahymena* RSs heads are connected, proteins present in the neighboring spokes are likely to be detected in co-IP and BioID assays, all together making the analyses more difficult. Therefore, we identified the RS1, RS2, and RS3-specific components and built a proteomic model of *Tetrahymena* RSs by combining our proteomic and cryo-ET data (Figure 6, Table S7 and S8) with previously published cross-linking data ^49^. For a simplicity, here we compared our RS model with that of *Chlamydomonas* ^19,41^.

**Figure 6.**
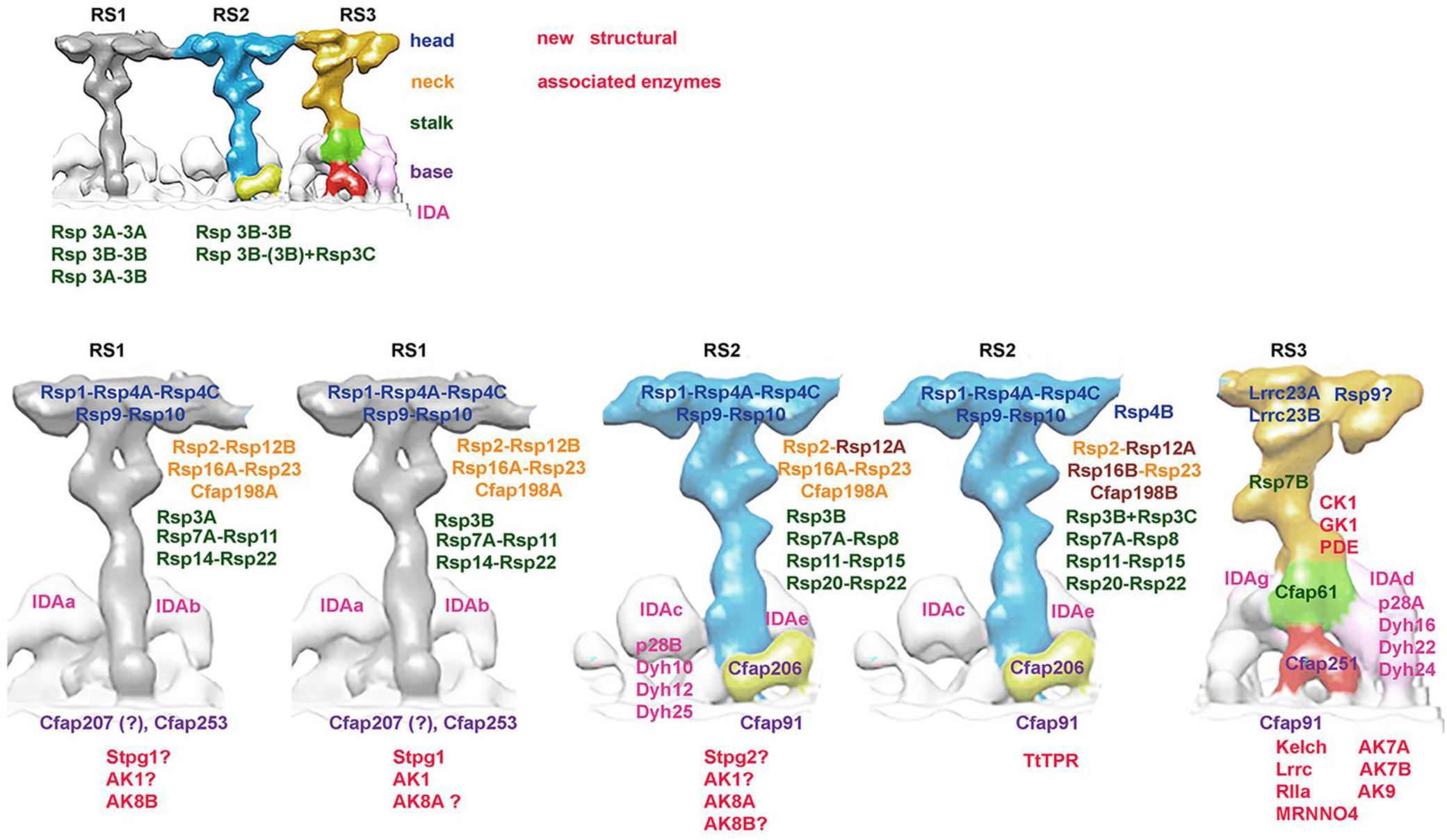
A schematic summary of the RS protein composition in *Tetrahymena* cilia prepared based on collected ultrastructural and proteomic data. A fragment of the Figure 3B published in our earlier paper^24^ was used as RSs template.

#### Components of radial spoke head

In *Chlamydomonas,* the RS head consists of two identical lobes connected by an RSP16 dimer, a neck component. Each lobe, besides paralogous RSP4 and RSP6, is composed of RSP1, 5, 9, 10, a part of the neck protein, RSP2, and a C-terminus of RSP3 ^15,19,41^. *Tetrahymena* has three paralogous of RSP4/6, Rsp4A, Rsp4B, and Rsp4C, and a single ortholog of Rsp1, Rsp2, Rsp9, and Rsp10 but lacks RSP5 ortholog. The levels of all head proteins, except for Rsp4B, were significantly diminished in RSP3B-KO mutant, lacking all RS2 and two-third of RS1 spokes, but not in RSP3A-KO, in which only fraction of RS1 were missing. Based on available data and earlier RS1 models ^4,5,19,20^ we propose that *Tetrahymena* RS1 and RS2 heads are composed of the same subunits except Rsp4B, which was almost eliminated in RSP3C-KO cilia (Table S2) suggesting that Rsp4B is present in Rsp3C-containing RS2 spokes.

The protein composition of the RS3 head is not fully resolved. It was recently reported that leucine-rich repeat-containing 23 (LRRC23) is a subunit of the RS3 head in human and mouse sperm axoneme ^22^. In *Tetrahymena*, the orthologous Lrrc23A and Lrrc23B were unaltered in mutants affecting only RS1 and/or RS2 but reduced in *CFAP91* mutant ^25^. Both Lrrc23 proteins have a C-termini enriched in glutamic acid residues (especially Lrrc23B) which may increase the negative charge of the RS3 head surface. We propose that in *Tetrahymena*, Lrrc23A and Lrrc23B are RS3 head components.

Among proteins biotinylated in cells expressing BirA*-tagged RS head proteins, were also central apparatus components (Table S9), mainly subunits of the C1b, C1d, and C2b projections, suggesting that primary these projections interact with RS head or that contact between RS head and these projections is longer, so their components can be more efficiently biotinylated. Furthermore, C1d components, Cfap46 and Cfap56 are preferentially biotinylated when RS1-RS2 head components were expressed as BirA*fusions, while in cilia where RS3 head Lrrc23 were fused with BirA*, besides C1d, also C2b and C1b subunits were biotinylated, suggesting possibility of some preferences in RS-CA projection interactions.

#### Components of radial spoke neck

In *Chlamydomonas*, the V-shaped neck is composed of RSP2, RSP12, RSP16, RSP23/FAP67, FAP198, and FAP385 ^19^. *Tetrahymena* has a single Rsp2, likely a single ortholog of Rsp23, and paralogous Rsp12A and Rsp12B, Rsp16A and Rsp16B, Cfap198A and Cfap198B. The 70 amino acids long FAP385 is strikingly similar to the very N-terminal fragment of adenylate kinase 8A (Ak8A). *Tetrahymena* and *Chlamydomonas* RSP2 and RSP23 orthologs share similarity only within their N-termini containing predicted DPY-30 ^60^ and NDK (nucleoside diphosphate kinase) ^61^ domains, respectively. In *RSP2* knockdown cilia (Table S2), only Rsp12A and Rsp12B were significantly diminished, suggesting that Rsp12 likely directly binds to Rsp2. The genome of *Tetrahymena* encodes two proteins similar to Rsp23, Cfap67A (TTHERM_000372529) and Cfap67B (TTHERM_00266490). Although neither of them was diminished in analyzed mutants, Cfap67A but not Cfap67B was identified in co-IP and BioID assays, suggesting that Cfap67A is the main if not the only one Rsp23. Interestingly, recent single-particle cryo-EM and molecular modeling studies showed that Cfap67A is a A-tubule MIP protein ^62^. Thus, based on previous and here presented data we propose that out of two *Tetrahymena* Cfap67 orthologs, Cfap67A likely dominates and has double role in cilia - as RSP and as MIP.

Based on the available data (Table S7) we propose that the neck region of all types of RS1 is composed of Rsp2, 12B, 16A, 23/Cfap67A, and Cfap198A. The RS2 lacking Rsp3C likely have a neck composed of the same proteins except that Rsp12B is replaced by Rsp12A, while Rsp2, 12A, 16B, 23/Cfap67A, and Cfap198B are subunits of RS2-containing Rsp3C (Figure 6).

#### Components of radial spoke stalk and base

Rsp3 paralogs form a core of RS1 and RS2 stalk. The moderately diminished levels of Rsp3B and Rsp3C proteins in *CFAP91* mutant (Table S2 and S7) can be due to defect in some RS2 ^25,48^. In *Chlamydomonas*, several proteins were assigned to RS stalks ^19^. Similar to *Chlamydomonas*, *Tetrahymena* has two ARM-like (armadillo-like) motif-containing RS proteins (Rsp8 and Rsp14) and LRR-containing, Rsp15. Rsp14 is a subunit of RS1 while Rsp8 and Rsp15 are RS2 components, independent upon the Rsp3 paralog forming RS stalk. *Chlamydomonas* RSP7 and RSP11 form a heterodimer positioned near RSP8 or RSP14 ^19,63^. Out of two RSP7 orthologs, *Tetrahymena* Rsp7A is a component of RS2 and likely of RS1 while Rsp7B is an RS3 subunit. *Tetrahymena* Rsp11, a small (7.9 kDa), basic protein (pI = 9) with a predicted RIIa domain and a limited similarity to *Chlamydomonas* RSP11 ^63^, similar as Rsp7A contributes to both, RS1 and RS2 structure.

*Chlamydomonas* RSP20/CaM and RSP22/LC8 (cytoplasmic dynein light chain 2) ^40,41^ are RS1 and RS2-base proteins. CaM binds to FAP253 between RS1 and RS1-docked IDAs while RSP22/LC8 forms multimer, likely, a RS docking site, composed of four homodimers in RS1 and two homodimers in RS2 ^19^.

Our proteomic data suggest that in *Tetrahymena* Rsp20/CaM and Rsp22/Lc8 associates with RS2 and perhaps RSP3B-containing RS1. On the other hand, the cryo-ET data showed that the RS1 base is missing only in a minor fraction of 96-nm subunits in RSP3A-KO mutant (Figure 3) which coincides with the unaltered levels of RS1 base proteins, Rsp20/CaM1, Rsp22/Lc8, Cfap207, and Cfap253 (Table S2 and S8) and together suggest their presence at the base of Rsp3A-containing RS1.

*Chlamydomonas* FAP207, a MORN motif-containing protein, was modelled as a part of the base of both RS1 and RS2, while the IQ motif-containing FAP253, orthologous to *Ciona* CMUB116 ^18^ and murine Iqub ^20^ was shown to be an RS1 adaptor ^19,20^. *Tetrahymena* and *Chlamydomonas* FAP253 are smaller than mammalian orthologs and lack ubiquitin domain predicted in IQUB. In all tested *Tetrahymena* mutants, the level of Cfap253 was unaltered but protein was biotinylated in cells expressing BirA*-tagged Rsp3A or Rsp3B, together indicating that Cfap253 is likely an RS1 adaptor. Similar to Cfap253 and Lc8, the level of Cfap207 was unaltered in *RSP3A* knockout but diminished in cells with RS2 defects except for *RSP3C* mutant suggesting that Cfap207 presence at RS2 base is Rsp3C-independent. To summarize, presence of different Rsp3 paralogs as core components of the RS1 and RS2 spokes diversified the protein composition of RS head and neck but not RS stalk.

The RS3 structure was unaltered in all RSP3-KO mutants, indicating that RS3 is a Rsp3-less spoke in *Tetrahymena.* Previous analyses led to identification of Cfap61 and Cfap251 as RS3 stalk and base components ^24^ and Cfap91 as a protein playing a role in RS2 base and RS3/RS3 base stability ^25^. Here we showed that RS3 stalk contains also Rsp7B, which has limited similarity to Rsp7A.

### New radial spoke candidate proteins

We have identified several uncharacterized proteins whose level was diminished in analyzed mutants, suggesting that those are either new RS subunits or proteins positioned in the RS base vicinity (Figure S11, Tables S2, S10, and S11). Based on the identified domains, we divided those proteins into two groups, contributing to the RS or RS-vicinity structure and with putative enzymatic activity (shortly enzymes).

The tetratricopeptide repeat-containing (TTHERM_00623040, TtTpr) and Kelch repeat-containing (TTHERM_00760390, TtKelch) proteins (Tables S4, S5, S7, and S10), missing vertebrate or *Chlamydomonas* orthologs, likely locates near the base of Rsp3C-containing RS2 and RS3 base, respectively. Also Lrrc (TTHERM_00046820), Lrrc (TTHERM_01084360), RIIa domain (TTHERM_00537370), and MRNN04 (TTHERM_00324550) could contribute to RS3 or its vicinity. Furthermore, two sperm-tail PG-rich repeat protein (STPG1 and STPG2, the later similar to Stpg2 in mice and *Chlamydomonas* CHLRE_09g415650v5), were reduced in RS1 or RS2 mutants. We propose that Stpg proteins are not RS components but are rather position near the RS base.

Cilia beating depends upon a constant supply of ATP and its uniform distribution within the cilium. Ciliary ATP is either delivered from the cell body by diffusion or produced within cilia ^64^. Adenylate kinases (Ak) reversibly catalyze the ATP+AMP↔2 ADP reaction and thus locally control the ATP level and maintain the homeostasis of adenine nucleotides ^65^. Six Aks (types 1, 7, 8, and 9) identified in co-IP and BioID assays were also diminished in different RS mutants (Table S11). Based on these data we propose that Ak8B is a component of Rsp3A-containing RS1, Ak1 and Ak8A (whose very N-termini is similar to FAP385) are associated with Rsp3B-containing spokes, while Ak7A, Ak7B, and Ak9 which were reduced in CFAP91-KO cilia but unaltered in *RSP3* mutants, likely associate with RS3 or are located in its vicinity. Interestingly, the Ak8B in *RSP3A* mutant and Ak7B and Ak9 in *CFAP91* knockout cilia were nearly completely eliminated strongly suggesting that Rsp3A-containing RS1 and Cfap91 stabilized RS structures are the main if not the only localization of those enzymes. Interestingly, guanylate kinase GK1 and phosphodiesterase, PDE are likely also RS3-associated enzymes and cGMP could play a role in the signal transduction in RS3 vicinity.

The RSP3 has an AKAP domain ^66^ and thus locally regulates PKA localization. PKA functions as a tetramer composed of two regulatory and two catalytic subunits ^67^. We identified two PKA catalytic subunits and one regulatory subunit whose levels were altered in RS mutants (Table S11). Two of these subunits co-precipitated with Rsp3A and or Rsp4. Interestingly, one of the *Tetrahymena* casein kinase 1 (Ck1)-type enzymes, was completely eliminated in CFAP91-KO cilia (Table S2) suggesting its specific association with RS2/RS3 base or their vicinity.

### RS defect-related instability of inner dynein arms

Single-headed IDAs are docked in pairs at the RSs’ bases, dynein a and b at RS1, c and e at RS2, and d and g at RS3. They are composed of IDA type-specific dynein heavy chain (Dyh), actin, and centrin (dyneins b, e, g) or light chain protein, p28 (dyneins a, c, d) ^68^. The RS defects affected some IDAs (Table 12). *Tetrahymena* has three orthologs of the single-headed IDA light chain protein p28: p28A, p28B, and p28C ^69^. The level of p28B diminished in *RSP3B* and *CFAP206* knockouts indicating together with cryo-ET data that p28B is a dynein c subunit. In CFAP91-KO, not only the level of p28B was reduced but also that of p28A ^25^ suggesting its/their presence in RS3 docked IDAd.

The levels of Dyh10, 12, and 25 diminished in the *RSP3B* and *CFAP206* knockouts, which coincides with lack of dynein c density (Figure 3) and ^48^, as well as in CFAP91-KO mutants in which Cfap206 is nearly completely eliminated ^25^. In contrast, dynein e was unaffected in those mutants suggesting differences in dynein c and dynein e docking. As we previously reported, in *CFAP91* knockout, besides Dyh10, 12, and 25, the levels of Dyh16, 22, and 24 are also reduced ^25^ suggesting that these dyneins are IDAd or IDAg components.

## Discussion

Identification of RSs components and RS-interacting proteins is crucial to understand the mechanism(s) enabling transduction of signal from the central apparatus to dynein arms. Despite the extensive research conducted using diverse ciliated species, including *Chlamydomonas* ^19,40,41^, *Tetrahymena* ^24,25,48^, *Ciona intestinalis* ^70–72^, and mice ^5,20,22,27,47,73^, the RS3 protein composition was only recently shown. Here, using bioinformatics, genetic, proteomic, and cryo-ET approaches, we solved RSs protein composition including RS3 in a ciliate *Tetrahymena*, showing spokes’ heterogeneity and complexity. Strikingly, we found differences not only between RS1, RS2, and RS3 spokes, but also identified likely existence of the subtypes of RS1 and RS2, generated primarily by the presence of different Rsp3 paralogs. The advantage of having Rsp3 paralogs which, based on our data, do not substitute one another, is not clear. The presence of three (Rsp4) or two (Rsp7, Rsp12, Rsp16, and Cfap198) paralogs further diversifies RSs’ structure and perhaps function. The existence of paralogous RS proteins agrees with *Tetrahymena* genome analyses revealing duplication of some genes encoding proteins involved in sensing or structural complexity^74^.

To the best of our knowledge, besides *Chlamydomonas pf14* mutant, cilia or flagella of other species with either deleted or mutated RSP3, were not analyzed using cryo-EM. Thus, this study shows for the first time, that Rsp3 is not a RS3 component. This conclusion agrees with previous studies addressing RS3 protein composition and with AlphaFold2 predicted models ^5,24–26^. Strikingly, while damaging mutations in so far studied genes encoding RS1 and RS2 components, cause PCD ^23,29,34,51,75–77^, mutations in genes encoding RS3 proteins, *CFAP61* ^45,46^, *CFAP91* ^44^, *CFAP251* ^43^, and *LRRC23* ^22^ result in male infertility but not PCD. Exception is the RS1 base protein, CFAP253/IQUB whose mutation also cause male infertility but not PCD ^20,27^. Accordingly, in mice, Cfap61, Cfap91, Cfap251, Lrrc23, and Iqub/Cfap253 are highly expressed in testis but not in lung or brain ^20,46,47^. Thus, either RSs (especially RS3) contribute differently to the regulation of sperm flagella and multiciliated cell cilia motion or the protein composition of RS1 base and RS3 is not identical in those organelles ^26^. Based on here presented data and previous *Tetrahymena* mutants’ analyses ^24,25^, it seems that *Tetrahymena* cilia, with their complex 3D waveform, are more similar to mammalian flagella than to cilia of multiciliated cell.

It seems that *Tetrahymena* Rsp3 paralogs differently contribute to the RS structure and are preferentially recognized by other Rsp and/or RS-interacting/associated proteins. In RSP3A-KO and RSP3B-KO mutants, the RS1 base remains intact and the level of RS1-base protein, Cfap253 ^19,20^ is similar to that in wild-type cilia. Strikingly, deletion of *RSP3B* but not *RSP3A* or *RSR3C* reduces the level of Cfap207 and the RS2-base component, Cfap206. Moreover, deletion of *CFAP206* has a stronger impact on the level of Rsp3C than Rsp3B, together suggesting that Rsp3C could dock to Cfap206 which in turn is perhaps partly stabilized by Rsp3B. In *Tetrahymena*, the RS2 is docked to the A-tubule by front, side, and back prongs and knockout of *CFAP206* in the vast majority of subtomograms (83%) eliminates the entire RS2 and front and back prongs. In remaining 17%, RS2 was nearly unchanged while front prong was missing and back prong reduced^48^. Interestingly, the number of RS2 remaining in *CFAP206* and *RSP3C* knockouts is similar. Based on the above data, it is also possible that the side prong maintained in CFAP206-KO cilia, could be an Rsp3B dimer-dependent structure.

It is proposed that the RS head can transiently interact with CA projections and that such interactions enable transmission of signals from the central apparatus via RSs to dynein arms. These interactions can be mechano-chemical and electrostatic in nature ^1,2,5,19^. Compared to the Rsp3A and Rsp3B, the C-terminal end of Rsp3C is enriched in glutamic acid residues. If the Rsp3C C-terminal end, similar to the *Chlamydomonas* RSP3, is a part of the RS head, its presence increases the negative charge of the head surface and thus might locally modulate the electrostatic interactions with CA projections. Interestingly, the N-terminal part of Rsp3C is also unusual in having an ARF domain, which could play a role in signaling.

Our data suggest that heads of the RS1 and RS2 are likely build of the same proteins (except for Rsp4B, likely co-assembling with Rsp3C), while RS3 head contains Lrrc23 paralogs. This data agree with observations in mice and humans carrying mutations in Rsph1/RSPH1 or Rsph4A/RSPH4A showing that in multiciliated cell, only RS1 and R2 spokes are head-less ^23,29^ and a recent finding that Lrrc23 mutation results in headless RS3 ^22^.

Whether heads of all three RS interact with CA projections in a similar way and if all CA projections are involved in such interactions, is still an open question. Based on fitted multi-scale axonemal structure, Meng and co-authors proposed a rigid contact mode between C1d and Rsph4a of DMT8 and elastic contact between C2b-C2d and Rsph1 of DMT4 ^5^. Our BioID data indicate that during cilia beating the BirA* ligase is close enough to the central apparatus to biotinylate some of the CA components (Table S9). Interestingly, Cfap46 and Cfap54, both the subunits of C1d projection ^78^ were biotinylated when BirA* was fused to Rsp3A, Rsp4A or Rsp4C. In contrast, in cells expressing Lrrc23-HA-BirA* fusions, Cfap54 was more prominently biotinylated than Cfap46. Moreover, in Lrrc23-BirA* expressing cells, hydin and Cfap47, the C2b components ^79^ and Spef2A and androglobin (Adgl), the C1b subunits ^80^, were also biotinylated. According to the model of *Chlamydomonas* CA, FAP46 and FAP54 form the outer surface of C1d while CPC1/SPEF2 and FAP42 build the distal part of C1b ^81,82^. We previously showed that Adgl has limited similarity to FAP42 ^80^ and therefore likely has similar position within C1b projection. Taken together, the above data support existence of the interactions between head of RS1 and RS2 and C1d as well as RS3 head and C1d, and long projections, C1b and C2b. The subunits of other projections if biotinylated, were identified by a low number of peptides. This raises a question about the involvement of remaining CA projections in CA-RS interactions calling for further studies.

### Differences in IDA docking

Destabilization of RSs affects some IDAs docking. In RSP3B-KO mutant the level of Dyh10, 12, and 25 was significantly reduced which coincided with the lack of dynein c (Figure 3). Dynein c (cryo-ET) and Dyh10, 12, and 25 (proteomic studies) were also missing in *Tetrahymena* cells with knocked out *CFAP206* encoding RS2-base protein ^48^ and this work. Strikingly, another RS2 dynein, dynein e, is not affected in RSP3B-KO or CFAP206-KO cilia, suggesting differences in the dynein c and dynein e docking. Indeed, the atomic model of *Chlamydomonas* RS2 shows that IDAc docks to the RS2 through FAP207 and IDAc subunit, p28 ^19^. In *Tetrahymena* RSP3B-KO and CFAP206-KO mutants, the level of Cfap207 is reduced (Table S8) strongly suggesting that also in *Tetrahymna* docking of IDAc depends upon Cfap207.

We did not find IDA defects in RSP3A-KO and RSP3C-KO mutants affecting RS1 and RS2, respectively. These observations agree with the atomic models of *Chlamydomonas* RS1 and RS2, suggesting that although FAP207 is present at the base of both RS1 and RS2, the p28 dimer of IDAa interacts with RS1-specific FAP253 ^19^. Accordingly, in *Tetrahymena* the level of Cfap253 remains unaltered in all studied RS mutants.

In *Tetrahymena*, knockout of *CFAP91* primarily affects the RS3 and the base of RS2 as the levels of Cfap206 and Cfap207 are substantially diminished (^25^ , and this work). In CFAP91-KO cilia, besides Dyh10, 12, and 25 (dynein c), the levels of Dyh16, 22, and 24 are moderately reduced ^25^ and this work), suggesting that these dynein heavy chains are components of IDAd or IDAg. Of note, Dyh22p and Dyh15p co-precipitate with GFP-tagged RS3-base protein, Cfap251^24^. Some of the remaining dyneins could be the components of IDAa or IDAb. Of note, in growing *Tetrahymena* cells, Dyh13, 17, 18, and 23 are expressed at a very low level (TGD Gene expression profiles and our proteomic data) and thus are likely minor components of *Tetrahymena* dynein arms or expressed during a specific life-phase.

Out of three *Tetrahymena* p28 paralogs ^69^, p28B is likely the main if not the only p28 ortholog present in RS2base-docked IDAc. In CFAP91-KO cilia, besides p28B, also the level of p28A is reduced ^25^ and this study. Thus, p28A is likely a subunit of IDAd but it remains to be determined if it forms homo-or heterodimers with p28B.

### RS-associated proteins containing predicted enzymatic domains

The global comparative analyses of wild-type and RS mutant ciliomes led to another interesting discovery – the identification of several enzymes whose level was reduced in RS mutants suggesting that those enzymes likely dock to specific RSs. Among them are (i) adenylate kinases that together with arginine kinase locally regulate ATP homeostasis, and (ii) enzymes playing role in cGMP cycle, and (iii) serine-threonine kinases that could play a role in the control/regulation of RS-mediated signal transduction. Thus, the phenotypic outcome of the deletion of genes encoding RS structural proteins, likely is not a sole consequence of the RS damage and some IDAs-defects, but also changes in protein phosphorylation-mediated signaling, level of guanylate nucleotides, and the accessibility to the ATP-stored energy.

Adenylate kinases locally control the ATP levels and maintain the homeostasis of adenine nucleotides ^65^. We have found that adenylate kinases of specific type likely associate with specific RS. Strikingly, the presence of AK7B, AK8B, and AK9 in cilia nearly completely depend upon the presence of certain class of RS suggesting that RSs are their sole docking sites. Our data are in agreement with recently published molecular modeling also suggesting association of adenylate kinases with radial spokes ^26^. Interestingly, also Gk1 and PDE enzymes were nearly completely eliminated from CFAP91-KO cilia, suggesting their presence near RS3 or its vicinity. GK1, is one of two Tetrahymena guanylate kinases. GK catalyzes phosphate transfer from ATP to GMP, producing GDP and ADP. The phosphorylation of GDP to GTP is catalyzed by nucleoside-diphosphate kinases (NDK), and GTP can be converted to cGMP by guanylate cyclase, while 3^’^5’-cyclic nucleotide phosphodiesterase (PDE) hydrolyzes cGMP ^83–85^. The RSP23 has NDK domains ^61^ however, its level remains unchanged in analyzed mutants. Early analyses of the ciliary beating in Paramecium showed that cyclic nucleotide monophosphates-driven chemical signaling controls ciliary beat ^86–88^. Thus, the immotility of the CFAP91-KO mutants might be caused not only by the lack of RS3 and some IDAs ^25^ but also by the perturbation of the guanidine nucleotide homeostasis. It is tempting to speculate that some RSPs undergo phosphorylation and that the phosphorylation status controls RS-mediated transduction of the regulatory signals. We found that the level of one of the casein kinase 1-type enzymes is completely eliminated in *CFAP91* mutant while the levels some cAMP-dependent protein kinase (PKA) subunits, are altered in RS mutants. The presence of Ck1 in the RS3 vicinity was confirmed by co-IP and BioID studies. In *Chlamydomonas,* CK1 docks to outer doublets and regulates the phosphorylation of IC138, the IDAI1/f intermediate chain ^89^. In *Tetrahymena*, this role can be played by other Ck1-type enzymes.

## Materials and Methods

### RESOURCE AVAILABILITY

#### Lead contact

Further information and requests for the resources and reagents should be directed to the lead contacts.

#### Materials availability

Plasmids, *Tetrahymena* mutants generated during this study are available from the lead contact

#### Data and code availability

Raw data generated from mass spectrometry analyses (co-IP, BioID, LFQ, and TMT data) are available upon request (will be deposited in the pubic repository).

### EXPERIMENTAL MODEL

#### Tetrahymena cell culture

*Tetrahymena thermophila* (Tetrahymena Stock Center) cells were grown to the mid-log phase (2-4x10^5^ cells/mL) with shaking (80-110 rpm) at 30°C. Wild-type cells and motile mutants were cultured in a standard SPP medium (1% proteose peptone, 0.1% yeast extract, 0.2% glucose) ^90^, while mutants with major ciliary defects were grown in a rich MEPP medium (2% proteose peptone, 2 mM Na-citrate, 1 mM FeCl_3_, 30 µM CuSO_4_, 1.7 µM CaCl_2_) ^91^, both supplied with an antibiotic-antimycotic mix (Sigma-Aldrich, St. Louis, MO, USA) at 1:100 (SPP) or 1:50 (MEPP). Before biolistic transformation and BioID assay, cells at mid-log phase were washed in 10 mM Tris pH 7.4 and next grown overnight (14-22 h) in the same buffer.

### METHODS DETAILS

#### Generation of Tetrahymena mutants

Engineering and phenotypic analyses of *Tetrahymena* mutants with deleted *CFAP61* ^24^, *CFAP206* ^48^, or *CFAP91* ^25^ were described before. The *RSP3* genes’ fragments used to obtain knock-out and knock-in transgenes were amplified by PCR with the addition of restriction sites using Phusion^TM^ Hot Start II DNA high-fidelity polymerase (Thermo Fisher Scientific) and the appropriate primers listed in Table S13.

#### Knock-outs

To delete a part of the *RSP3* genes (0.8-1 kb), approximately 1.2-1.5 kb fragments positioned upstream and downstream of the targeted gene fragment were amplified and cloned subsequently on both sites of the neo4 resistance cassette ^92^. Approximately 60 µg of plasmid was digested with ApaI and SacII to separate the transgene from the plasmid backbone and precipitated onto approximately 0.63 mg of 0.5-0.8 nm gold particles (Thermo Fisher Scientific) by mixing with 1 M CaCl_2_ and 20 mM spermidine (final concentrations). DNA-coated gold particles were washed with ethanol and place on macrocarriers (Bio-Rad) and after drying used to transform conjugating *Tetrahymena* cells (strains CU428 and B2086) at the early stages of meiosis. Transformed cells were selected in SPP medium supplemented with 1.5 µg/mL CdCl_2_ and 100 µg/mL paromomycin, followed by selection in 6-methylpurine (15 µg/mL)-supplemented SPP. Double resistant-cells after sexual maturation and validation by crossing to CU427 cells, were crossed to infertile A*III strain to obtain heterokaryons. Next, heterokaryons were crossed to obtain knockout cells ^93,94^. The deletion of the targeted fragments was confirmed by PCR.

To eliminate a fragment of the RSP2 gene, we used a co-deletion approach ^95^ and primers listed in Table S13. The targeting plasmid (∼15 µg) was introduced to conjugating CU428 x CU427 cells using biolistic gun as described above. After 14 hours cells were transferred to SPP medium, grown for 6 hours and exposed to paromomycin selection (100 µg/mL) on 96-well plates. To verify *RSP2* gene fragment deletion, the genomic DNA were purified from the wild-type (control) and paromomycin-resistant cells and the extend and completeness was analyzed by PCR using primers listed in Table S13.

#### Knock-ins

To express RS proteins as fusions with C-terminal -3HA, -HA-BirA*, or -HA-TtBCCP tag under the control of the transcriptional promoter, approximately 1 kb fragments of the open reading frame immediately upstream of the STOP codon and ∼1 kb fragment of the 3’UTR were amplified by PCR as described above (primers used are listed in Table S13) and cloned into pCfap44-3HA-neo4, pCfap44-HA-BirA*-neo4, and pCfap44-TtBCCP-neo4 plasmids, respectively ^96,97^ to replace fragments of *CFAP44* gene. To express Lrrc23 with C-terminal Turbo tag (TurboID) we amplified Turbo coding region from IFT52-Turbo plasmid adding restriction sites and replaced BirA* coding region. To express proteins with C-terminal GFP tag under the control of the transcriptional promoter and pPur cassette enabling selection of transformed *Tetrahymena* cells with puromycin ^98^, the coding sequence of the 2xV5 tag in pCfap44-2V5-pPur plasmid ^96^ was replaced by GFP coding region and fragments of *CFAP44* gene were replaced by the fragments of a coding region and 3’UTR of the gene of interest as described above. Approximately 10-15 µg of plasmid was digested with MluI and XhoI to separate the transgene from the plasmid backbone and precipitated onto approximately 0.20 mg of 0.5-0.8 nm gold particles (Thermo Fisher Scientific) as described above. Approximately 10^7^ CU428 cells were transformed by biolistic transformation and after 2 h recovery in SPP medium supplied with 1.5 µg/mL CdCl_2_, cells were transferred to 96-well plates and transformants were selected using 100 µg/mL paromomycin for 3-4 days.

To co-express Rsp3B-3HA with Rsp3A-GFP or Rsp3C-GFP, Rsp3B-3HA cells were transformed with appropriate constructs as described above except that after transformation cells were grown for 24 h in a SPP medium supplied with 2.5 µg/mL CdCl_2_. Positive transformants were selected on 96-well plates using 200 µg/mL puromycin.

After selection transformants and double transformants were grown in SPP medium supplied with the growing concentration of paromomycin or/and puromycin and reduced concentration of CdCl_2_ to promote transgenes assortment ^99^.

#### Phenotypic analyses

The measurements of the cell swimming rate and the analyses of cilia beating (amplitude, waveform and frequency) were described in detail ^25^. Briefly, for swimming rate analyses, cells at a density of 2-3x10^3^ cells/mL were viewed and recorded at room temperature using a Zeiss Discovery V8 Stereo microscope (Zeiss, Oberkochen, Germany) equipped with a Zeiss Plans 10_ FWD 81 mm objective and an Axiocam 506 camera, and ZEN2 (blue edition) software. The length of the trajectories was measured using ImageJ software and the color lines parallel to trajectories were added in the Adobe Photoshop program. For each strain experiments were done in triplicates and at least 100 trajectories were registered and analyzed in each experiment. Cilia beating was analyzed as described in ^25^. Cells from a mid-log phase were cultured at room temperature for 3 hours, centrifuged, placed between two pieces of adhesive tape fixed on the glass slide, covered with a coverslip, and recorded using a Phantom Miro C110 high-speed camera (Vision Research, Wayne, NJ, USA) mounted on an AXIO Imager M2 microscope (Zeiss, Germany) with either a 40 x oil immersion lens (analyses of cilia beating frequency) or a 63x oil immersion lens (numerical aperture 1.4, analyses of ciliary waveform). Videos were recorded at 900 frames/s. For each strain at least 10-15 cells were recorded, aligned in ImageJ and analyzed using ImageJ (frequency) or Adobe Photoshop (waveform, amplitude).

#### Immunofluorescence

To analyze the localization of HA-tagged Rsp proteins, cells from the overnight culture were fixed 1:1 v/v on coverslips with a mix of 1% Triton and 4% PFA/ or 1% NP-40 substitute and 4% PFA, both in a PHEM buffer (12 mM PIPES, 5 mM HEPES, 2 mM EGTA, 0.8 mM MgSO_4_, pH 6.9). After drying and blocking with 3% BSA in PBS, cells were stained overnight at 4 °C with a mix of primary antibodies, (i) rabbit monoclonal anti-HA antibody (BioLegend, San Diego, CA, USA) 1:300 and mouse monoclonal an anti-acetylated α-tubulin antibody 6-11 B1 (1:2000) or (ii) mouse monoclonal anti-HA (1:200) and rabbit polyclonal polyG (1:2000) ^100^. To co-localize Rsp3 paralogs, cells were stained with a mix of primary antibodies: mouse monoclonal anti-HA 16B12 (1:200) (BioLegend, San Diego, CA, USA) and rabbit polyclonal anti-GFP (1:6000) (Abcam). After washing with PBS, samples were stained for 1.5 h at RT with a mix of the secondary antibodies, anti-mouse and anti-rabbit IgG, conjugated with either Alexa-488 or Alexa-555 (Invitrogen, Eugene, OR, USA) both diluted 1:300. Coverslips were mounted in Fluoromount-G (Southern Biotech., Birmingham, AL, USA). Cells were recorded using either a Zeiss LSM780 (Carl Zeiss Jena, Germany) or a Leica TCS SP8 (Leica Microsystems, Wetzlar, Germany) confocal microscope.

#### Deciliation of Tetrahymena cells

For immunofluorescence analyses of regenerating cilia, cells co-expressing 3HA and GFP-tagged Rsp3 paralogs were grown overnight to mid-log phase, rinse with 10 mM Tris-HCl, pH 7.4 buffer. Approximately 4x 10^5^ cells were collected, suspended in 100 µL of a 10 mM Tris-HCl, pH 7.4 buffer and mixed with 1 mL of 10% Ficoll in 10mM Tris-HCl, pH 7.5 and immediately deciliated by passing 5-6 times through the syringe needle (diameter 0.8 mm). After 60-90 sec cells were transferred to 10 mL of SPP medium for cilia regrowth.

For biochemical analyses cilia were purified from 6-8 x 10^7^ cells using a pH shock method ^101^. In brief, cells were collected, rinsed with Tris-HCl buffer, pH 7.4, resuspended in deciliation buffer (10 mM Tris-HCl, pH 7.4, 10 mM CaCl_2_, 50 mM sucrose), and deciliated by addition of acetic acid (10.5 mM final concentration). After 1-1.5 min the pH was raised to physiological level with potassium hydroxide (10.8 mM final concentration). Deciliation was monitored under a microscope. Deciliated cell bodies were separated from cilia-containing supernatant by centrifugation (twice at 1680 x g for 5 min) and cilia were collected by centrifugation at 26,900 x g for 30 min. Next, cilia were resuspended in deciliation buffer supplemented with protease inhibitors (cOmplete mini EDTA-free protease inhibitor cocktail, Roche Diagnostics GmbH, Mannheim, Germany) and protein concentration was estimated using Pierce™ BCA Protein Assay Kit (Thermo Scientific, Bartlesville, OK, USA). Purified cilia were further analyzed using Western blotting, mass spectrometry (ciliomes), co-immunoprecipitation or BioID assays.

#### Co-immunoprecipitation and proximity labeling (BioID) assays

All buffers used in biochemical studies were supplemented with protease inhibitors (cOmplete mini EDTA-free protease inhibitor cocktail, Roche Diagnostics GmbH, Mannheim, Germany).

For co-immunoprecipitation assay, approximately 6 x 10^7^ cells from a mid-log phase culture either wild-type (control) or expressing -3HA tagged Rsp proteins, were spun down and washed with 10 mM Tris-HCl buffer (pH 7.4). After deciliation, collected cilia were re-suspended in deciliation buffer and combined with an equal volume of 2% NP-40 and 1.2 M NaCl in 80 mM Tris-HCl buffer, pH 7.5. After 15 min incubation on ice, axonemes were pelleted at 21,000 × g, 4°C, for 15 min and treated with 0.5 M KI, 30 mM NaCl, 5 mM MgSO_4_, 0.5 mM EDTA, 1 mM DTT in 10 mM HEPES, pH 7.5. After 30 min on ice, the axonemes were centrifuged (21,000 × g for 15 min at 4 °C). The supernatants were diluted 500x with 50 mM Tris–HCl, pH 7.4 and concentrated on ultracentrifugation columns (Vivaspin® Turbo 4, Sartorius, Niemcy). Collected proteins (0.5-1 mg) were incubated overnight with agarose beads-conjugated anti-HA antibody (Thermo Fisher Scientific, Waltham, MA) at 4 °C. The bead-bound proteins were identified by mass spectrometry.

The proximity-labeling (BioID) assay ^102^ was performed as described in detail ^80^. Wild-type were used as a control because ∼30 kDa BirA* tag can enter cilia by diffusion and randomly biotinylate components of different ciliary complexes. Briefly, approximately 6x10^7^ wild-type or Rsp-HA-BirA fusion expressing cells from mid-log phase, were cultured for 16-18h in 10 mM Tris-HCl buffer (pH 7.4) and next incubated with 50 µm biotin for 4 h at 30°C in the same buffer. After deciliation, cilia were suspended in axoneme stabilization buffer (20 mM potassium acetate, 5 mM MgSO_4_, 0.5 mM EDTA in 20 mM HEPES, pH 7.5) supplied with 0.2% NP-40 to release unbound biotin. After 5 min, the axonemes were spun down (10 min, 21,000x g, 4 °C), washed, suspended in lysis buffer (50 mM Tris–HCl, pH 7.4, 0.4% SDS, 0.5 M NaCl, 1 mM DTT), and incubated at RT for 1 h. After spinning down (8000x g at 4 °C), the collected supernatant was diluted with 50 mM Tris–HCl, pH 7.4 (1:3) and incubated overnight with streptavidin-coupled Dynabeads (Dynabeads M-280 Streptavidin, Thermo Fisher Scientific, Waltham, MA) at 4 °C. The bead-bound biotinylated proteins were analyzed by Western blot with use of HRP-conjugated streptavidin (1:40 000) (Thermo Fisher Scientific, Rockford, IL, USA) and identified by mass spectrometry.

#### Protein gel electrophoresis

Approximately 30 µg of ciliary proteins or 10% of bead-bound proteins were loaded on the SDS-PAGE polyacrylamide gel, separated in standard conditions and transferred to nitrocellulose membrane for 1h at 170 mA. Two-dimensional electrophoresis was performed as described ^25^. Approximately 30 µg of ciliary extract purified from cells expressing Rsp-3HA fusion proteins were cleaned with ReadyPrep 2-D cleanup (Bio-Rad) and dissolved in rehydration buffer (7 M urea, 2 M thiourea, 2% CHAPS, 0.1%, Tergitol NP7, 40 mM DTT and 0.2% BioLytes). Protein solution was used to rehydrate ReadyStrip IPG pH 3-10 or 4-10 (Bio-Rad) for at least 12 h. Proteins were separated in Protean IEF Cell (Bio-Rad) for ∼80kVh, at 4000V. and next subjected to standard SDS-PAGE and transferred to nitrocellulose membrane.

#### Western Blot

After transfer to nitrocellulose and blocking for 1h with either 5% skimmed milk in TBST, blots were incubated with mouse monoclonal anti-HA antibody (1:2000) or rabbit polyclonal anti-GFP antibody (1:60,000) both diluted in 5% skimmed milk in TBST (overnight, 4°C). After washing (4 x 10 min, TBST) and 1h incubation at RT with HRP-conjugated secondary antibodies: goat anti-mouse IgG (1: 10 000 ) (Jackson ImmunoResearch, West Grove, PA, USA) or goat anti-rabbit IgG (1:20,000) (Sigma-Aldrich) and washed as before. To analyze biotinylated proteins nitrocellulose was blocked with 3% BSA in TBST, incubated with HRP-conjugated streptavidine (1:40,000 in 3% BSA in TBST, 3h, RT) (Thermo Fisher Scientific) and washed again (4x10 min TBST). Proteins were visualized using a Westar Supernova kit (Cyanagen, Italy).

#### Cryo-ET preparation

The axonemes for cryo-ET were cross-linked with glutaraldehyde (final concentration 0.15%) for 40 min on ice and quenched with 35 mM Tris pH 7.5. The axoneme solution at 3.6 mg/mL was mixed with 5 (Cytodiagnostics) or 10 (Aurion) nm gold beads in a 1:1 ratio for a final axoneme concentration of 1.8 mg/mL. Inside the Vitrobot Mk IV (Thermo Fisher) chamber, 4 μl of crosslinked axoneme sample was applied to negatively glow discharged (10 mA, 10 s) C-Flat Holey thick carbon grids (Electron Microscopy Services). The sample was incubated at 23 °C and 100% humidity for 45 seconds on the grid, followed by 8 seconds of blotting with force 0 and plunge frozen in liquid ethane.

#### Cryo-ET acquisition and reconstruction

Tilt series were collected using the dose-symmetric scheme from -60 to 60 degrees with an increment of 3 degrees at 2.12 Å per pixel using a Titan Krios equipped with Gatan K3 and BioQuantum energy filter. The acquisition was performed using SerialEM ^103^. The defocus for each tilt series ranges from -2.5 to -6 μm. The total dose for each tilt series is 120 to 160 e-per Å2. For each view, a movie of 10-13 frames was collected. Motion correction of each view was performed with Alignframes ^104^. Tomograms were reconstructed using IMOD ^105^.

#### Subtomogram Averaging

CTF estimation for each tilt series was performed with WARP ^106^. The doublet microtubule for subtomogram averaging was picked using IMOD by tracing the line along the microtubules ^105^. Subtomogram averaging of the 4-times binned 96 nm repeating unit of WT and mutant strains was performed using the “axoneme align” program ^107^. The subtomogram coordinates and alignment parameters were converted to Relion 4.0 for local refinement and classification ^108^. The resolutions for the 96-nm repeating unit of the axoneme of wild type, *RSP3A-KO*, *RSP3B-KO*, and *RSP3C-KO* are 18, 20, 22, and 17 Å, respectively. Axonemal repeats from *RSP3A* and *RSP3B* mutants showed strong heterogeneity in the occurrence of RS1 and RS2. To analyze the heterogeneity of the mutant strains, three-dimensional (3D) classification without alignment was performed in Relion by masking RS2 and RS1. The axoneme of the *RSP3C* mutant strain showed heterogeneity in the RS2 head and neck region. Therefore, unsupervised 3D-classification by masking the head and neck region was performed with three classes.

The 3D classification of n=2099 axonemal 96-nm repeats with RS1 defects revealed either the lack of the entire RS1 (n=639 units, ∼30%) or RS1 except for the RS1 base (n=1417, 67%). The RS1 in the remaining 43 units was not classified due to the low number of particles. The identification of intact RS1 in Relion was unsuccessful, likely due to the structural heterogeneity of intact RS1 in RSP3A-KO compared to WT and/or conformational flexibility enabling RS, especially RS1 to tilt ^5,6^. However, visual inspection of denoised tomograms revealed the presence of intact RS1 in ∼22% of units (n=455). Additionally, the averaged subtomogram map indicates that existing RS1 spokes are thinner in RSP3A-KO than in WT, suggesting potential structural differences associated with the knockout (Figure S6). In contrast, RSP3B-KO mutant lacked RS2 in all analyzed axonemal units (n=2092). Moreover, ∼77% of the axonemal repeats (n=1622) showed also RS1 defects; the RS1 structure was either missing (51%, n=1078) or had only a base part (26%, n=544). Among the collected n=2790 axonemal repeats, 173 axonemal repeats (∼7%) were unclassified. The analyses of the remaining n=2617 axomenal repeats revealed that ∼19% (n=527) lacked the entire RS2. The 3D classification of the remaining repeats using the RS2 mask covering an RS2 head and stalk showed that the remaining units grouped into two categories: (i) with an intact RS2 structure (n=469) and (ii) with a well-visible RS2 base (n=1621) (Figure S6).

For visualization, tomograms were CTF deconvolved and missing wedge corrected using IsoNet^109^. The UCSF ChimeraX package was used for the visualization of subtomogram averages, surface rendering, segmentation, and fitting ^110^.

#### Heat map

Heat maps illustrating the distribution of particles within each doublet were generated using Microsoft Excel. The doublet with the highest particle count was designated as the first doublet. To assess the distribution of subtomograms with intact RS2 across doublets, counts were normalized such that a value of ’0’ indicates no subtomograms with intact RS2, while ’100’ represents a doublet where all subtomograms have intact RS2. Tomograms were categorized into proximal and medial-distal regions based on the presence of the CCDC81B MIP signal. Tomograms exhibiting this signal were classified as proximal, while those lacking the signal were designated as medial-distal ^111^. Bar graph visualization and statistical analyses were performed using GraphPad Prism (GraphPad Software, San Diego, CA, USA). Mean percentages of intact RS2 subtomograms for each doublet were calculated in GraphPad Prism, treating tomograms as biological replicates to account for biological variability. Statistical significance was assessed using an unpaired t-test, and error bars represent the standard deviation across tomograms.

#### Differential Quantitative Cilia Proteome Analyses

##### Sample Preparation

Cilia purified from 5x10^7^ cells from mid-log phase culture were resuspended in 10 mM Tris pH 7.4 and protein concentration was determined with Pierce™ BCA Protein Assay Kit (Thermo Scientific, Bartlesville, OK, USA). Three hundred micrograms of protein were precipitated using ReadyPrep 2-D Cleanup Kit (Bio-Rad Laboratories, USA). Protein pellets were dissolved in 0.1% RapiGest in 500 mM Tetraethylammonium bromide (TEAB) and then incubated at 850 rpm for 45 min at 37 °C (Eppendorf Comfort Thermomixer, Eppendorf, USA). Proteins were digested by trypsin (Trypsin Gold, Mass Spectrometry Grade, Promega; protein:enzyme (w/w) ratio – 100:1, at 37 °C for 16 h) according to a standard protein digestion protocol including reduction (by 1,4-dithiothreitol) and alkylation (by iodoacetamide). The digestion reaction was stopped by the addition of 55% trifluoroacetic acid (final concentration of 5%), samples were centrifuged at 20,000 x g for 30 min at 4 °C to precipitate RapiGest, and pellets were discarded. Supernatants containing the obtained peptides were purified using Pierce™ Peptide Desalting Spin Columns (Thermo Scientific, Bartlesville, OK, USA) according to the manufacturer’s protocol, dried in a vacuum concentrator at RT (SpeedVac Concentrator Plus, Eppendorf, USA) and stored at -80 °C for further analysis.

##### LC‒MS/MS analysis of labeled peptides (TMT analysis)

Desalted peptides were dissolved in 100 µl of 100 mM TEAB solution, and peptide concentrations were determined using the Pierce™ Quantitative Fluorescent Peptide Assay (Thermo Scientific, USA). Next, a TMT labeling reaction was performed according to the procedure provided by the manufacturer (Thermo Fisher Scientific). Briefly, a volume of sample containing 30 µg of peptides was labeled with a corresponding tandem mass tag. The labeling reaction was carried out for 1 h at room temperature and quenched with 5% hydroxylamine. Additionally, a test sample was prepared to check the efficiency of labeling. Labeled peptides were purified and fractionated using liquid chromatography at high pH. Separation was carried out for 26 minutes at a flow rate of 0.8 ml/min using a UPLC system (Acquity UPLC Class H system, Waters). The mobile phases consisted of water (A), acetonitrile (B) and 100 mM ammonia solution (C). The percentage of phase C was kept constant at 10% throughout the separation. Fractions were collected every 1 minute, starting from the second minute of the run. The peptide elution was monitored spectrophotometrically at 214 nm. Twenty-four fractions were collected and combined to obtain 12 measurement samples. Samples were dried in a vacuum concentrator at room temperature. Peptides were resuspended in 100 μl of 5% acetonitrile and 0.1% formic acid.

Peptides were analyzed on the Evosep One system (Evosep Biosystems, Odense, Denmark) coupled to the Orbitrap Exploris 480 mass spectrometer (Thermo Fisher Scientific, USA) according to ^112^. One µg of peptides was loaded onto Evotips C18 trap columns (Evosep Biosystems, Odense, Denmark) according to the manufacturer’s protocol with some modifications. Chromatographic separation of peptides was carried out using a mobile phase flow rate of 500 nl/min in gradient elution mode for 44 min on an EV1106 analytical column (Dr. Maisch C18 AQ, particle size 1.9 µm, 150 µm x 150 mm, Evosep Biosystems, Odense, Denmark). The following gradient elution was used: 0 min – 1% B, 120 min – 35% B, 121 min – 95% B, 124 min – 1% B, 127 min – 95% B, 130 min – 1% B. The eluted peptides were ionized in the positive ion mode in the nano-ESI source with a capillary voltage of 2.1 kV and the temperature of transfer capillary 275 °C. Survey scans from 300 *m/z* to 1700 *m/z* were acquired by an Orbitrap mass analyzer (Thermo Fisher Scientific, Waltham, MA, USA) at a resolving power of 60 000. The resolving power in the MS2 spectrum measurement mode was 30 000 with the TurboTMT function set to TMT Reagents. HCD-MS/MS spectra (normalized collision energy of 30%) were generated for 25 multiply charged precursor ions from each survey scan. A precursor fit filter was applied to reduce peptide co-fragmentation. Dynamic exclusion was set to 20 s, and the precursor ion intensity threshold was set to 5x10^3^.

##### LC-MS/MS analysis of non-labeled peptides (label-free analysis)

Desalted peptides were resuspended in 100 μl of 5% acetonitrile and 0.1% formic acid and loaded on Evotips C18 trap columns and separated for 88 mins as described above. Positive ionization and DDA (data dependent acquisition) mode were applied to collect data. MS1 parameters were: resolving power 60,000, AGC target 300%, and the m/z range of 300 to 1600. MS2 parameters were: resolving power 15,000, normalized AGC target. Forty of the most abundant precursors ions within an isolation window of 1.6 m/z were fragmented. The intensity threshold was set up at 5x10^3^. The higher energy collisional dissociation mode with a normalized collision energy of 30% was applied for precursor ion fragmentation.

##### Data analysis

MS data were analyzed with FragPipe (v. 17.1) (Nesvilab, University of Michigan, Ann Arbor, MI), MSFragger (v. 3.4) (Kong et al., 2017) and Philosopher (v. 4.2.1) (da Veiga Leprevost et al., 2020). ProteoWizard’s MSConvert (v. 3.0.1908) (Palo Alto, CA) (Chambers et al., 2012) was used to convert the raw MS data to mzML format. The *Tetrahymena thermophila* UniProt database (canonical and isoform sequences; 27,027 entries) was searched using the following search parameters: (i) digestion enzyme-trypsin/P, up to two missed cleavage sites were allowed, (ii) precursor and fragment ion mass tolerance ±10.0 ppm and ±20 ppm, respectively, (iii) fixed modifications: carbamidomethyl (C), (iv) variable modifications: oxidation (M), deamidation (N), (Q). For TMT samples, additional fixed modification was used (v) TMT modification of lysine (+229.16293) and variable modification (vi) TMT modification of N-termini of protein and peptide (+229.16293). Proteins and peptides were identified using the target-decoy approach with a reversed database. The peptide mass range was set from 500 Da to 5 000 Da. The results were processed with FDR (false discovery rate) set to 1% at the PSM, peptide and protein levels. Quantitative analysis was performed using IonQuant (label-free quantitation) and TMT Integrator (TMT-based quantitation). Statistical analysis was performed with Perseus (v. 2.0.3) (Max Planck Institute of Biochemistry, Martinsried, Germany). In label-free quantitation, missing values were replaced based on quantile regression imputation of left-censored data (QRILC) ^113^. Student T-test was used for statistical analysis. Proteins were considered to be differentially expressed if the difference in abundance was statistically significant (FDR adjusted p-value < 0.05) and the fold change was equal to or higher than 1.5.

#### Quantification and Statistical analyses

Data are given as mean+/-SD and were compared via two-tailed distribution Student’s t-tests. For statistical analysis t-student test in majority of experiments was used. Perseus (v. 2.0.3) program was used for statistical analysis of TMT proteomic data.

#### Reagent availability

All plasmids and *Tetrahymena* mutants are available on request. LFQ and TMT mass spectrometry data and list of all protein identified in co-IP and BioID experiments will be submitted to the open repository. Nucleotide sequences of the primers used to amplified PCR fragments are provided in Table S13.

## Acknowledgment

We thank Dr. Jacek Gaertig for critical reading of the initial version of the manuscript. The IFT52-TurboID plasmid was obtained from Dr. Jarema Malicki. Confocal imaging was performed at the Laboratory of Imaging Tissue Structure and Function which serves as an imaging core facility at the Nencki Institute of Experimental Biology and is part of the infrastructure of the Polish Euro-BioImaging Node. Polish Node is supported by the project financed by the Minister of Education and Science based on contract No 2022/WK/05 (Polish Euro-BioImaging Node “Advanced Light Microscopy Node Poland”). We also thank the Facility for Electron Microscopy Research (FEMR) at McGill University for their support in cryo-electron tomography data collection. The mass spectrometry analyses were done in cooperation with the Mass Spectrometry Laboratory, Institute of Biochemistry and Biophysics, PAS, Warsaw, Poland and Institute for Biochemistry and Molecular Biology (IBMB), Medical Faculty, University of Bonn, Bonn, Germany. The authors have no competing financial interests to declare.

## Author contributions

Marta Bicka, Investigation, Validation, Formal analysis, Visualization

Avrin Ghanaeian, Formal analysis, Investigation, Writing – review & editing, Visualization Corbin Black, Investigation,

Ewa Joachimiak, Methodology, Investigation, Validation, Formal analysis, Visualization,

Funding acquisition, Writing—Review and Editing

Anna Osinka, Investigation, Visualization

Sumita Majhi, Investigation,

Anna Konopka, Methodology, Validation, Resources, Supervision

Ewa Bulska, Supervision, Resources , funding acquisition, writing - review & editing

Khanh Huy Bui, Investigation, Supervision, Resources, Funding acquisition, Writing – review & editing

Dorota Wloga, Conceptualization, Investigation, Validation, Visualization, Resources, Formal analysis, Supervision, Funding acquisition, Project administration, Writing – original draft, Writing – review and editing

