## Supplementary material for "Heterogeneity of radial spokes structural components and associated enzymes in *Tetrahymena* cilia": Supplementary Figures.pdf

**Table of content**

Supplementary Figures……………………………………………………………….2 - 27

Supplementary Tables………………………………………………………………28 - 47

Supplementary Movies………………………………………………………………47

#### Supplementary Figures

A

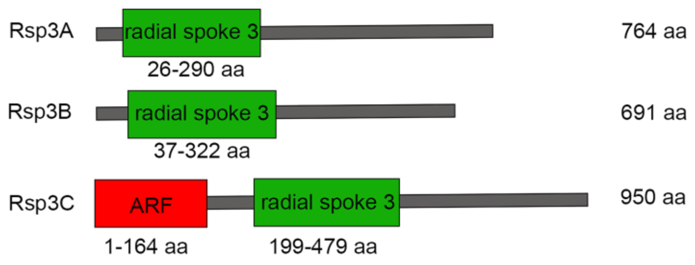

B

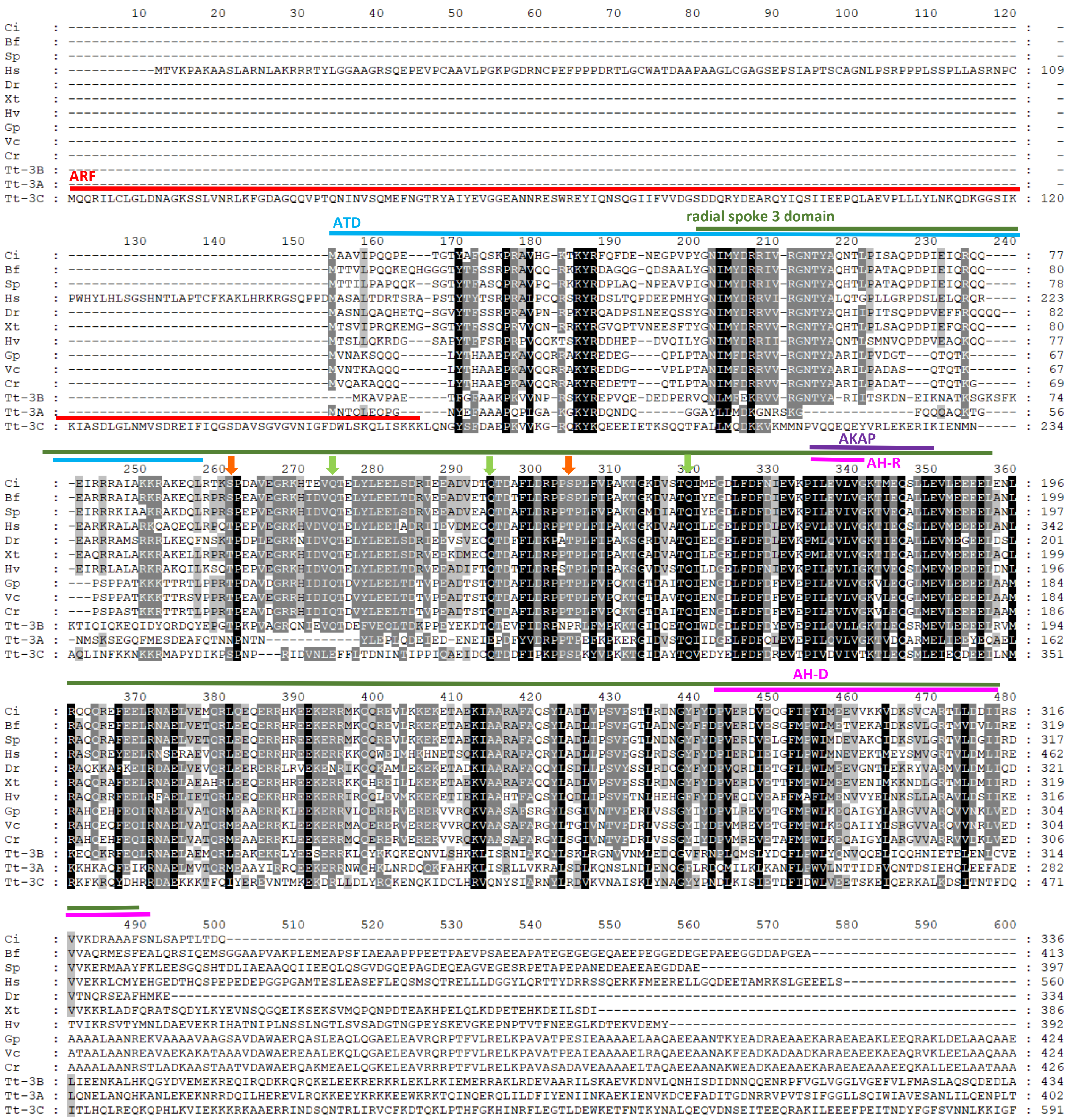

```
                 610       620       630       640       650       660       670       680       690       700       710       720
Ci     : ------------------------------------------------------------------------------------------------------------------------ :    -
Bf     : ------------------------------------------------------------------------------------------------------------------------ :    -
Sp     : ------------------------------------------------------------------------------------------------------------------------ :    -
Hs     : ------------------------------------------------------------------------------------------------------------------------ :    -
Dr     : ------------------------------------------------------------------------------------------------------------------------ :    -
Xt     : ------------------------------------------------------------------------------------------------------------------------ :    -
Hv     : ------------------------------------------------------------------------------------------------------------------------ :    -
Gp     : AAAERGEDPPEEPPQLDE--PLVDVEAVVAAAADAVQRPPVREVADIDILSYMLDKGIVTKDAIIQSLAVHALGDKAYVNHPAFADSS-------------------------------- :  510
Vc     : EAEERGEEPPEEPPTLDE--PLVDVEAVVAAAVEAVVKPPVKEVEDIDIISYMLDQGIITKDAIIQSLAVHALGDKAYINHPAFAEGA-------------------------------- :  510
Cr     : EAEERGEEPPAEPPSLPDGVEPVDVEAEVAKAVEAVPKPPVKEVTDIDILSYMMDKGAITKDAIIQALAVHALGDKAYTNHPAFAEAEGA------------------------------ :  516
Tt-3B  : SGARELINNQEAILNSFKELLTSVLSEGSIDLPINSQIDALLAQKGEDLSLATLATAQLGDAQDDVIDLIVSNVSSQILNAIREHSNEFGFPQETIDQLIYLIVQSYFSEESLQAKIKFS :  554
Tt-3A  : QNQIRDIVDLVLREFKHGLTIKFSESQKEFYDELVKQLSEATENIPEAREQFINNILENEEIYINDLDTISYLRNAIEKRNGSTQTFKEILKILLEYHAKNIKPDPIDEQKSKEEHNSSI :  522
Tt-3C  : TLAHSQFYVQTKKDRLLELVAFAIQKDGKIISRIDKDNLRPNNQNPSLNKYIRYKRNIRDLTKKASDDEAIVVDFSKVPDDVWGFFFFVKLPNLNLLADKLEKENEVLKNSRMGIEDYSN :  711

                 730       740       750       760       770       780       790       800       810       820       830       840
Ci     : ------------------------------------------------------------------------------------------------------------------------ :    -
Bf     : ------------------------------------------------------------------------------------------------------------------------ :    -
Sp     : ------------------------------------------------------------------------------------------------------------------------ :    -
Hs     : ------------------------------------------------------------------------------------------------------------------------ :    -
Dr     : ------------------------------------------------------------------------------------------------------------------------ :    -
Xt     : ------------------------------------------------------------------------------------------------------------------------ :    -
Hv     : ------------------------------------------------------------------------------------------------------------------------ :    -
Gp     : ------------------------------------------------------------------------------------------------------------------------ :    -
Vc     : ------------------------------------------------------------------------------------------------------------------------ :    -
Cr     : ------------------------------------------------------------------------------------------------------------------------ :    -
Tt-3B  : PVKPVKGFSQAGIAYLRPDLESLTINTRSPKFGQEPEIGEDEEPIKVVPENLVESEFSNKVSLLNPKDEDLEVLVQHAVAEQVVREDILRVIAGSIKGVDGHEYPILYEALSKTQEYQRK :  674
Tt-3A  : SHQRGSMNDLDDKLYRQKIRILFENTNEYIRNHANVIIRLRVPFLTQQEMEKIRQEAEKQSQEEQKSQNNSQLQQEQEEGSNQVQENQKEQEAQGVQEQLEQGQEQEQEQQPEEEKCKYD :  642
Tt-3C  : NIAIDGVNLMEKIQFDQLIQQPPSEESPTIQNSYYLGYSLTKRPITGWYLENIKIASKGMFEQAEEQEYLNKLVSFWTSTEGIPPKNIEASQVADGEEANVNGSPAQANNLRVMENQSQM :  831

                 850       860       870       880       890       900       910       920       930       940       950       960
Ci     : ------------------------------------------------------------------------------------------------------------------------ :    -
Bf     : ------------------------------------------------------------------------------------------------------------------------ :    -
Sp     : ------------------------------------------------------------------------------------------------------------------------ :    -
Hs     : ------------------------------------------------------------------------------------------------------------------------ :    -
Dr     : ------------------------------------------------------------------------------------------------------------------------ :    -
Xt     : ------------------------------------------------------------------------------------------------------------------------ :    -
Hv     : ------------------------------------------------------------------------------------------------------------------------ :    -
Gp     : ------------------------------------------------------------------------------------------------------------------------ :    -
Vc     : ------------------------------------------------------------------------------------------------------------------------ :    -
Cr     : ------------------------------------------------------------------------------------------------------------------------ :    -
Tt-3B  : VLDSIHNDFPVLDMKSY------------------------------------------------------------------------------------------------------- :  691
Tt-3A  : VFVETFDGEGYKRNLIENLNNSQRRPLNEDHPSQINIWVNEERRFLYFDERTLFYLEAVLHKVIRDVLGNKVEPINASQFSQYFDEIQDLVFENFLGDVYEEEKIEKIPIFDIDFGFNLP :  762
Tt-3C  : SLSNGTQFFQKCYKEYKVEFAKNENGELTANGKNYEEFIEELSQYIQSQDAALDALCQWGFEFYFRGKLLKRLQTLKKCKNAFEFEVKKKPEPIVEKVENGEEGEEGEGEEGEEEPEED- :  950

Ci     : -- :    -
Bf     : -- :    -
Sp     : -- :    -
Hs     : -- :    -
Dr     : -- :    -
Xt     : -- :    -
Hv     : -- :    -
Gp     : -- :    -
Vc     : -- :    -
Cr     : -- :    -
Tt-3B  : -- :    -
Tt-3A  : SE :  764
Tt-3C  : -- :    -
```

**Figure S1.** Domain analyses and multiple alignment of Rsp3 orthologs.

(A) Domain organization of *Tetrahymena* Rsp3 paralogs as predicted using SMART (http://smart.embl-heidelberg.de/) [1,2], showing radial spoke 3 domain (green rectangle)[3] and ARF (ADP-ribosylation factor) domain (red rectangle), uniquely present at the Rsp3C N-termini. (B) Multiple alignment of Rsp3 orthologs. The amino acid sequences of RSP3 orthologs were obtained from the NCBI protein database (National Center for Biotechnology Information) using Blastp search and human RSPH3 as a bait. The *Tetrahymena* Rsp3 orhologs were obtained from Tetrahymena Genome Database (https://tet.ciliate.org/). Protein amino acid sequences were aligned using ClustalX2 software [4] and edited using SeaView [5]. The identical and similar amino acid residues were shaded using GenaDoc [6] and the following color code: white letter on the black background (100% conserved residues); white letter on a grey background (80% conservation); black letter on a light grey background (60% conservation). The color lines indicate the position of the following domains: red: ARF domain in *T.thermophila RSP3C* (1-164 aa), please not that there is no homology to the N-termini of human RSPH3; green: SMART predicted radial spoke 3 domain (position:188-470 aa in HsRSPH3); blue: region corresponding to *Chlamydomonas* RSP3 axoneme targeting domain (ATD) located within amino acids 1-85 aa [7]; purple: AKAP domain, in Hs 317-332 aa [8]; pink: amphipathic helix (AH-R, 161-178 aa) binding RIIa domain and amphipathic helix (AH-D, 269-316 aa) binding DPY-30 domain in *Chlamydomonas* RSP3 [9]; green arrow: TQT-like motifs mediating interactions with LC8 [10]; orange arrow: position of the threonine residues, T243 and T286, phosphorylated in RSPH3 [8],

**Abbreviations** in the alphabetical order with the accession number.
*Branchiostoma floridae* (Bf, XP_035686603.1), *Chlamydomonas reinhardtii* (Cr, XP_001695406.1), *Ciona intestinalis* (Ci, NP_001027624.1), *Danio rerio* (Dr, AAI34868.1), *Gonium pectorale* (Gp, KXZ49444.1), *Homo sapiens* (Hs, BAB71615.1), *Hydra vulgaris* (Hv, XP_047128590.1), *Strongylocentrotus purpuratus* (Sp, XP_783727.2), *Tetrahymena thermophila* (Tt, 3A, TTHERM_01044600; 3B, TTHERM_00566810; 3C, TTHERM_00418270), *Volvox carteri f. nagariensis* (Vc, XP_002953748.1), *Xenopus tropicalis* (Xt, NP_998863.1).

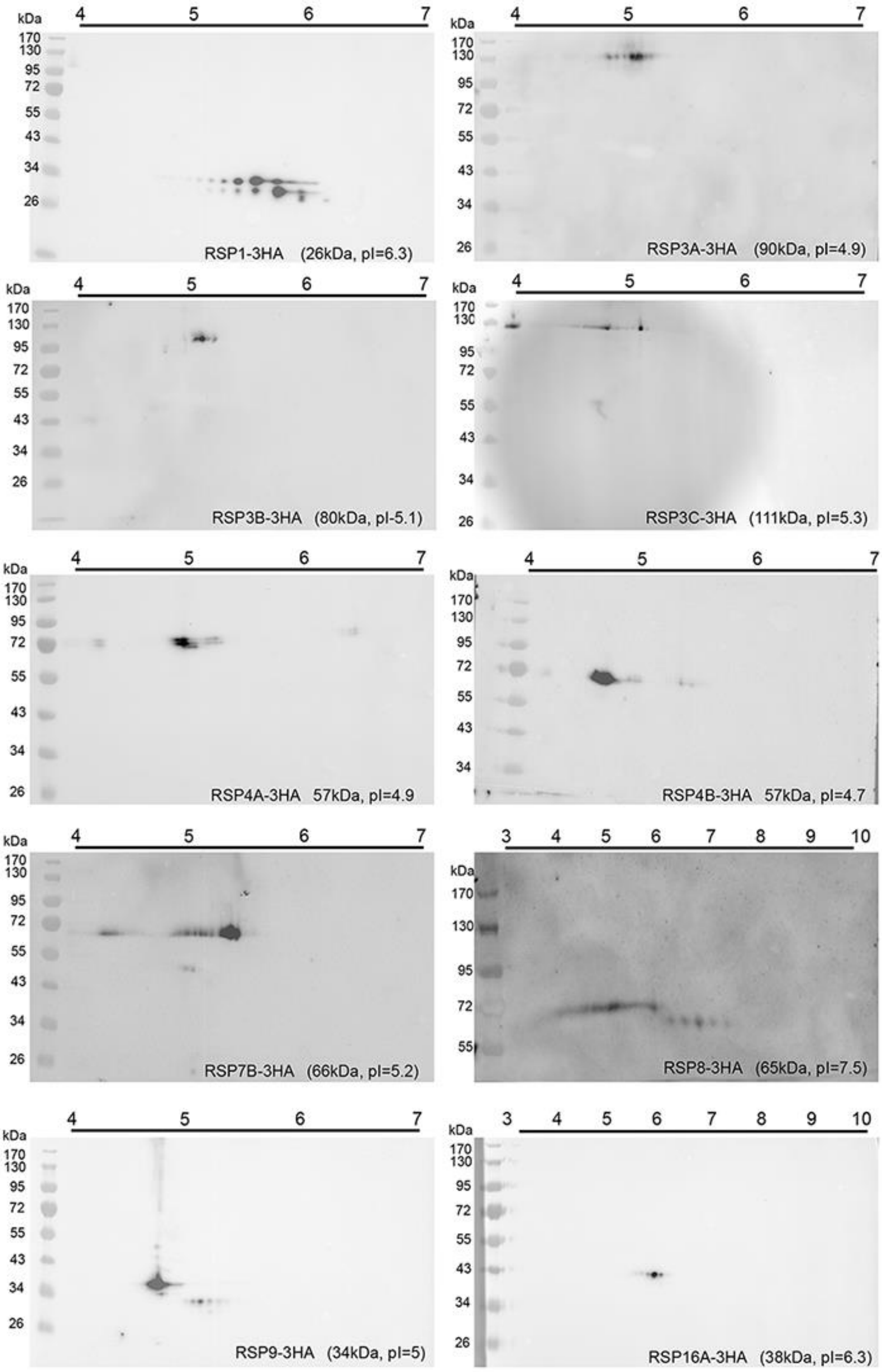

**Figure S2.** Two-dimensional analyses of the selected *Tetrahymena* Rsp showing presence of more than one isoform. The Rsp-3HA fusions were expressed under the control of the transcriptional promoters. The isoelectric focusing of the ciliary proteins (30 μg) was performed using 7 cm 4-7 or 3-10 ready strips. The theoretical pI and Mw values were calculated using https://web.expasy.org/compute_pi/. The Rsp isoforms were detected using anti-HA antibodies.

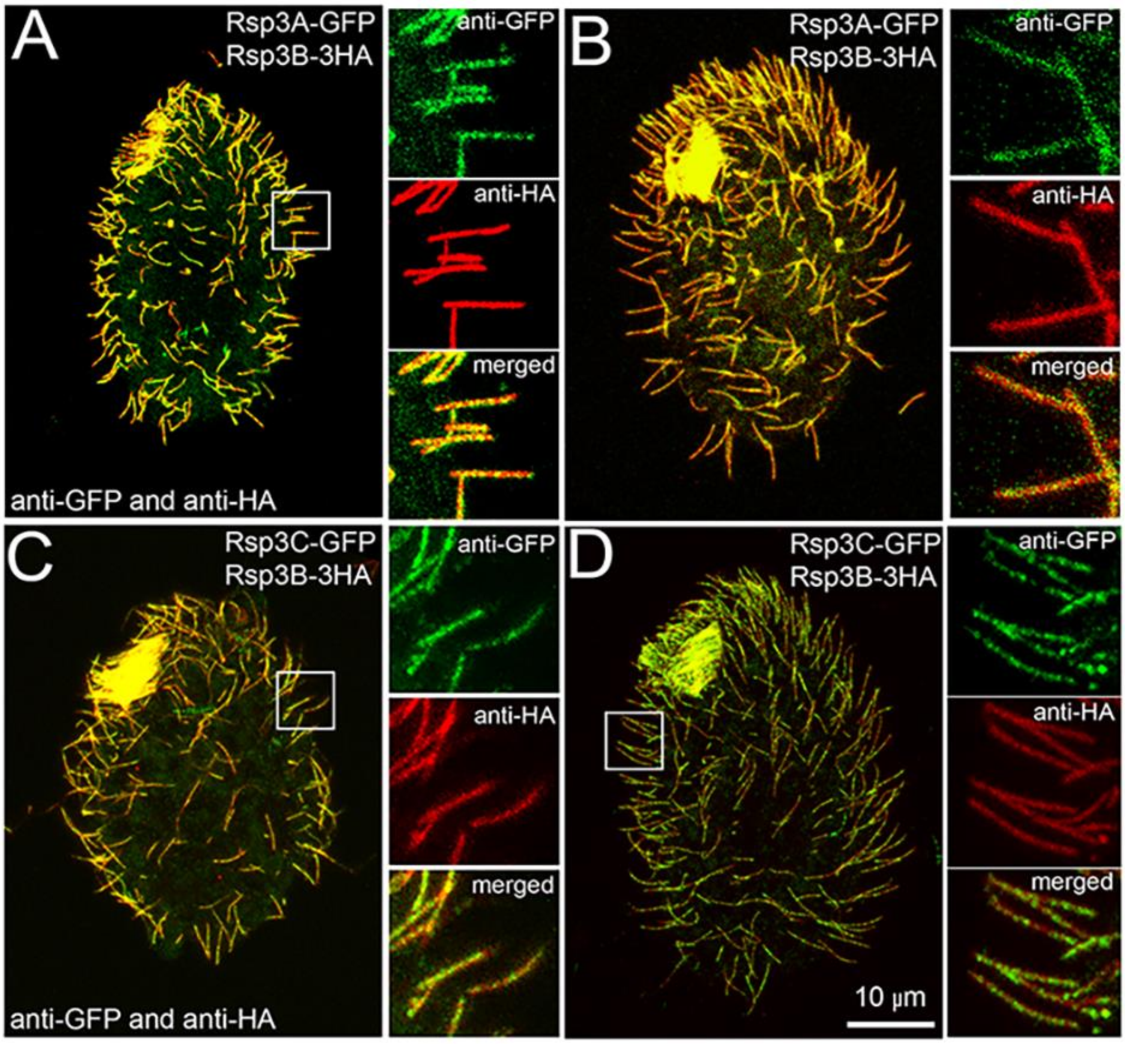

**Figure S3**. Rsp3 paralogs are simultaneously incorporated into regenerating cilia.

(A-D) Confocal immunofluorescence images of *Tetrahymena* cells co-expressing Rsp3B-3HA and either Rsp3A-GFP (A-B) or Rsp3C-GFP (C-D) under the control of transcriptional promoters, stained with anti-HA and anti-GFP antibodies. Cells were experimentally deciliated and grown in medium to enable cilia regeneration for 30 min (A, C) or 60 min (B, D). To the right, the magnified areas marked in the main image (white frame).

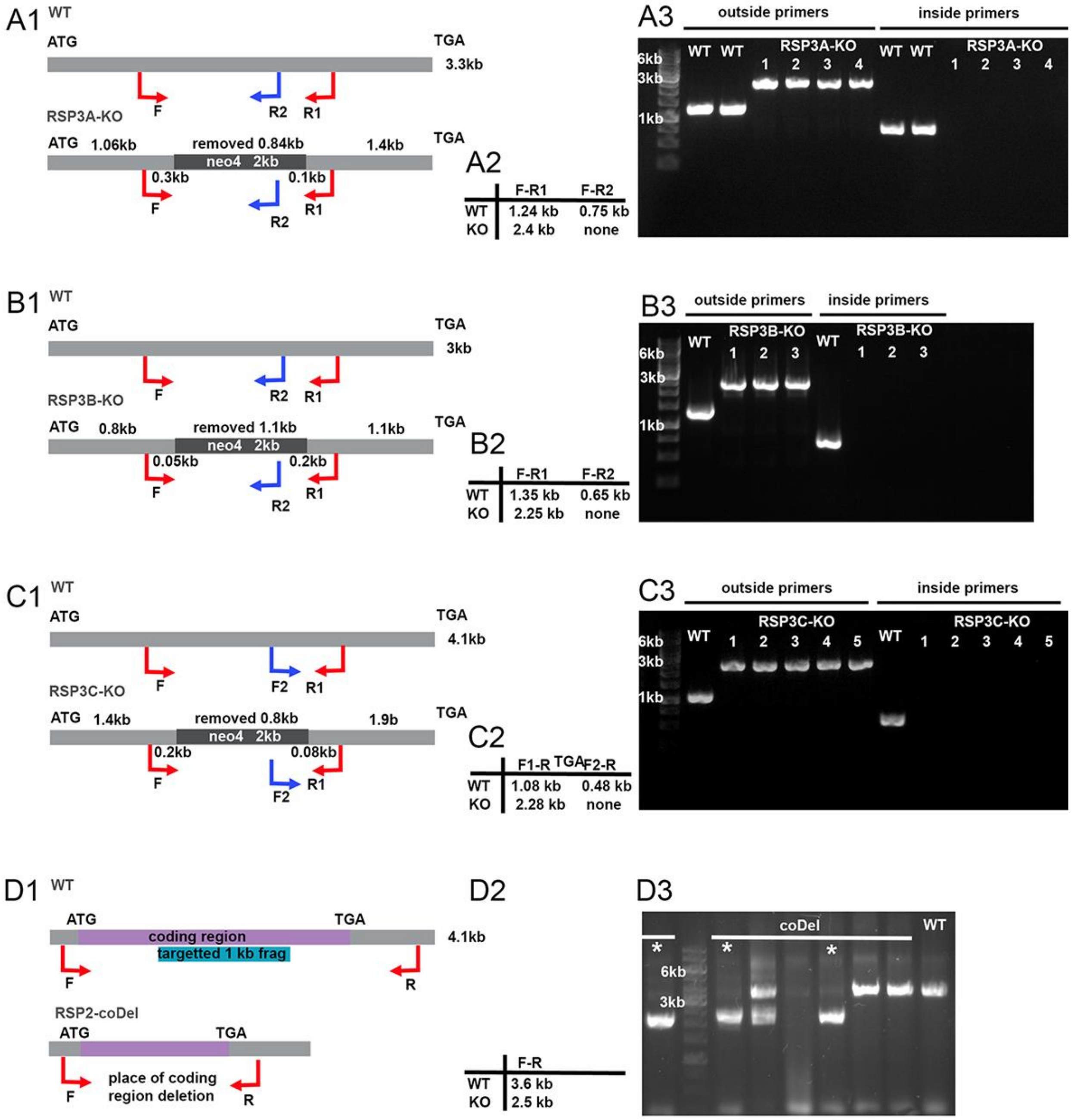

**Figure S4**. Alterations in *RSP3A*, *RSP3B*, *RSP3C*, and *RSP2* loci in engineered *Tetrahymena* knockout mutants. (A1, B1, C1, D1) Schematic representations of the *RSP3A* (A), *RSP3B* (B), *RSP3C* (C), and *RSP2* (D) loci in the wild-type genome (WT, upper gray bar) and obtained knockout cells (RSP mutants, lower gray bar) with the size of the fragment of the open reading frame removed in the knockout cells indicated and the position of the annealing of primers (blue and red arrows) used to test changes in the analyzed loci marked. Numbers near the red arrows indicate the distance of the primer annealing site from the neo4 cassette that replaced the removed fragment of the open reading frame in mutated loci. Note that the primer represented by a blue arrow recognizes the nucleotide sequence deleted in mutant cells. Numbers above the gray bars representing the open reading frame indicate the distance between either the start or stop codons and the neo4 cassette insertion site. (A2, B2, C2, D2) A table showing the expected size of the PCR products obtained using genomic DNA purified from wild-type (WT) and knockout cells (KO) and primers as indicated. (A3, B3, C3, D3) PCR analyses of the RSP3 loci *RSP3A* (A3), *RSP3B* (B3), and *RSP3C* (C3), and *RSP2* (D3) locus in independently obtained mutants using, in the case of RSP3 mutants, either both primers annealing outside the neo4 cassette or one of the primers recognizing the deleted gene fragment as indicated in A1, B1, and C1, respectively. (D3) star indicates clones with deletion (RSP2-coDel mutants).

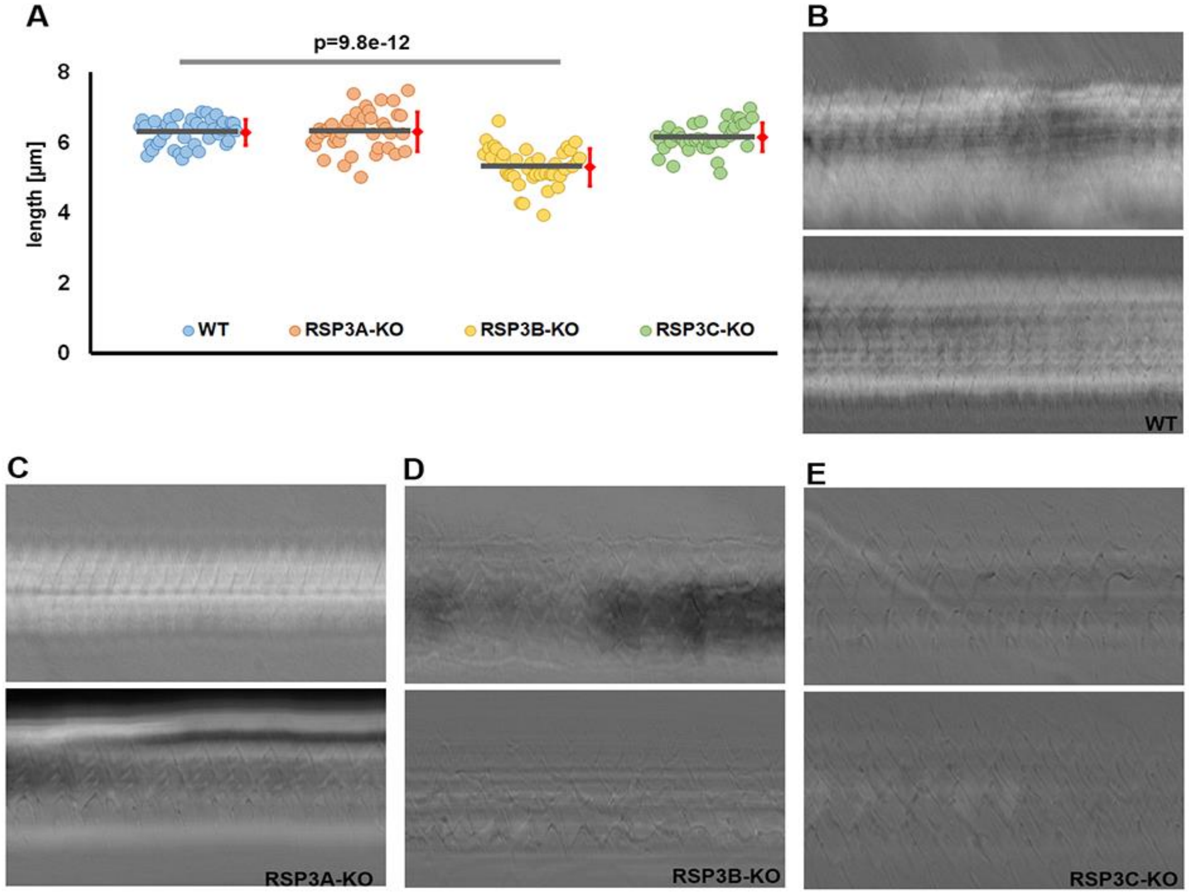

**Figure S5**. Deletion of a single *RSP3* gene had no effect on *Tetrahymena* cilia length but affected cilia beating. (A) Graphical representation of cilia length measurements. Wild-type and *RSP3* knockouts cells were double labeled with anti-HA and anti-acetylated tubulin antibodies (polyG in the case of RSP3C-KO), and cilia length was measured in confocal microscopy images using ImageJ program (N=40 cilia). Red bars represent standard deviation. Note that only deletion of *RSP3B* slightly affects cilia length (B-E) but the detected cilia shortening is statistically significant (t-test). Examples of the kymographs showing cilia beating synchrony. Moving cells were recorded using fast speed camera and x40 objective, and analyzed using ImageJ. Note that only knockout of *RSP3B* affects cilia beating synchrony.

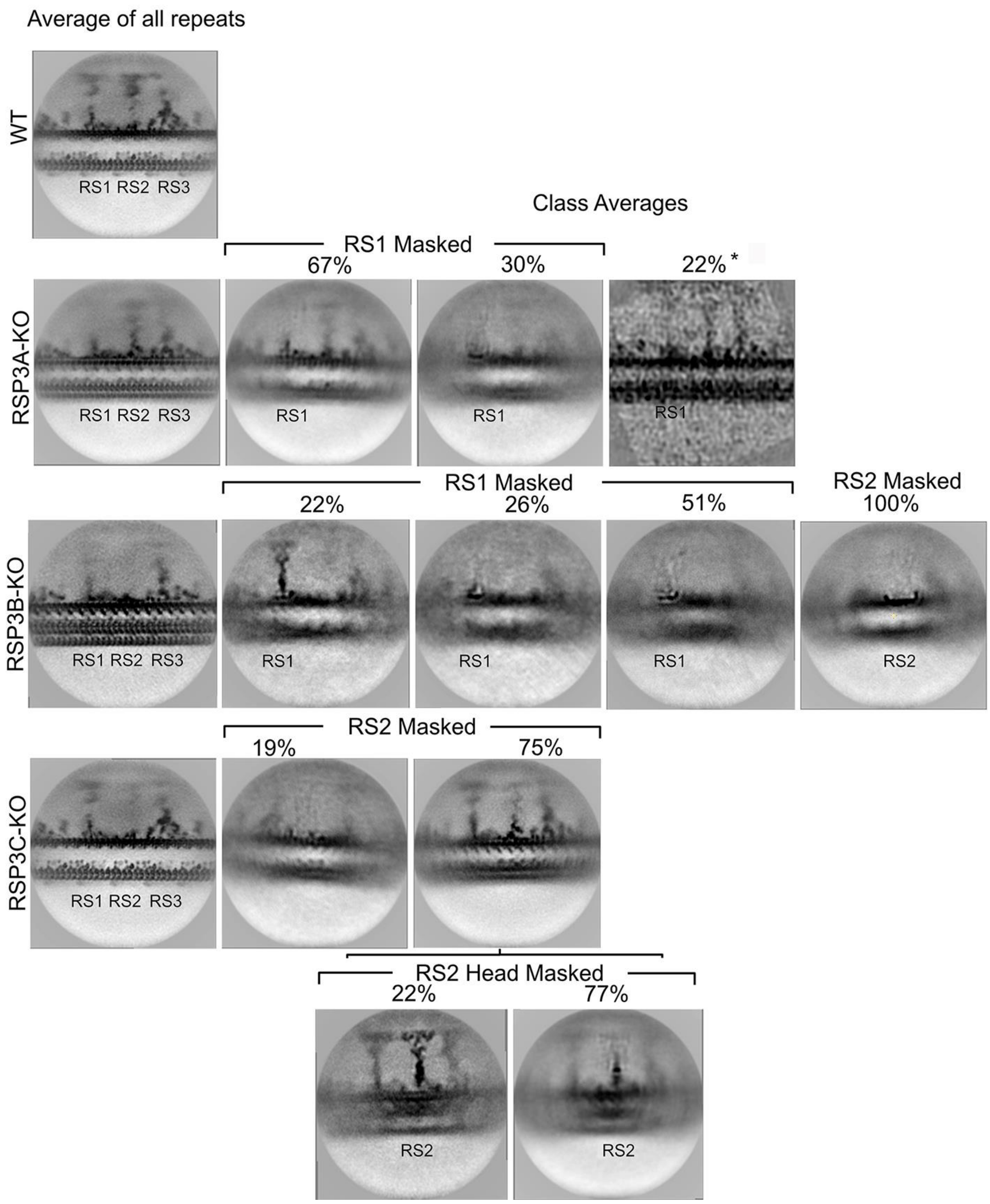

**Figure S6**. Consensus subtomogram averages and three-dimensional classification of the RSs in *Tetrahymena* RSP3 knockout mutants. The 3D classification in Relion of n=2099 axonemal repeats revealed either the lack of the entire RS1 (30%, n=639 units) or RS1 except for the RS1 base (67%, n=1417). Three percent of the recorded repeats were not classified due to the low number of particles. *3D classification for intact RS1 in Relion was unsuccessful; therefore, we performed manual classification using visual inspection in ArtiaX. Subtomogram coordinates were mapped onto the denoised tomogram, and those containing intact RS1 were manually selected in ArtiaX. Due to the limitation of missing wedge, we can only inspect two doublets showing RS1 in side view of each tomogram. On average, each doublet contained three intact particles, leading to an estimated 13 intact particles across all nine doublets. With 35 tomograms analyzed, the dataset contained approximately 455 out of 2090 intact particles (~22%). The selected subtomograms were then rotated based on a reference and averaged using a Python script. In RSP3B-KO mutant cilia, all analyzed axonemal repeats (n=2092) lacked RS2. Additionally, 77% of the axonemal repeats (n=1622) had RS1 spokes defects. The RS1 spokes were either completely missing (51%, n=1078) or only their base was well visible (26%, n=544). We collected n=2790 axonemal repeats of RSP3C-KO mutant cilia. Among them 173 axonemal repeats (6%) were unclassified, ~19% (n=527) lacked the entire RS2. The 3D classification of the remaining repeats using the RS2 mask covering an RS2 head and stalk showed two RS2 categories: (i) intact RS2 structure (~17%, n=469) and (ii) only well-visible RS2 base (~58%, n=1621).

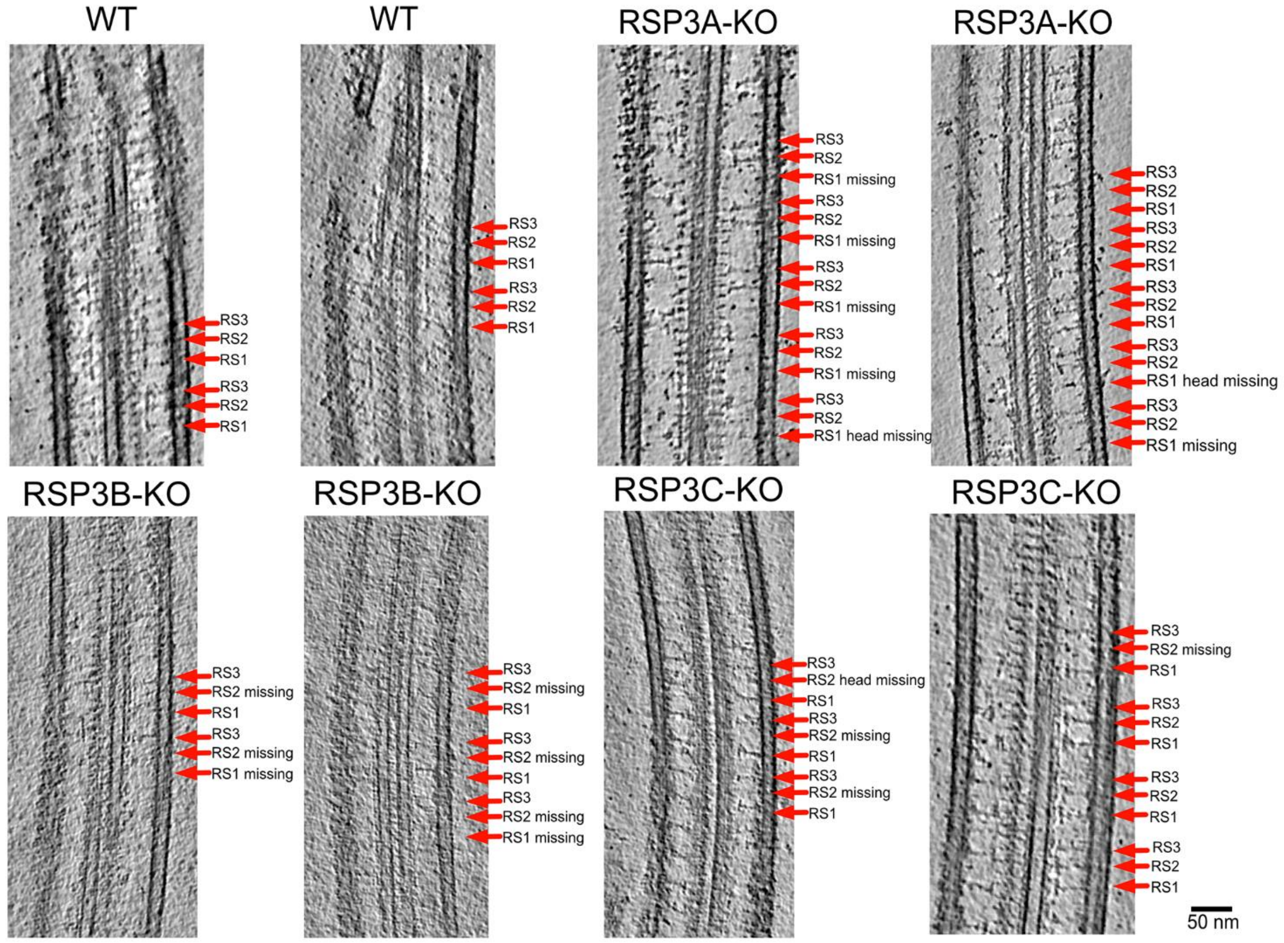

**Figure S7**. Tomographic slices from WT, RSP3A-KO, RSP3B-KO, and RSP3C-KO cilia showing the heterogeneity of distributions of RS with structural defects. Scale bar: 50 nm.

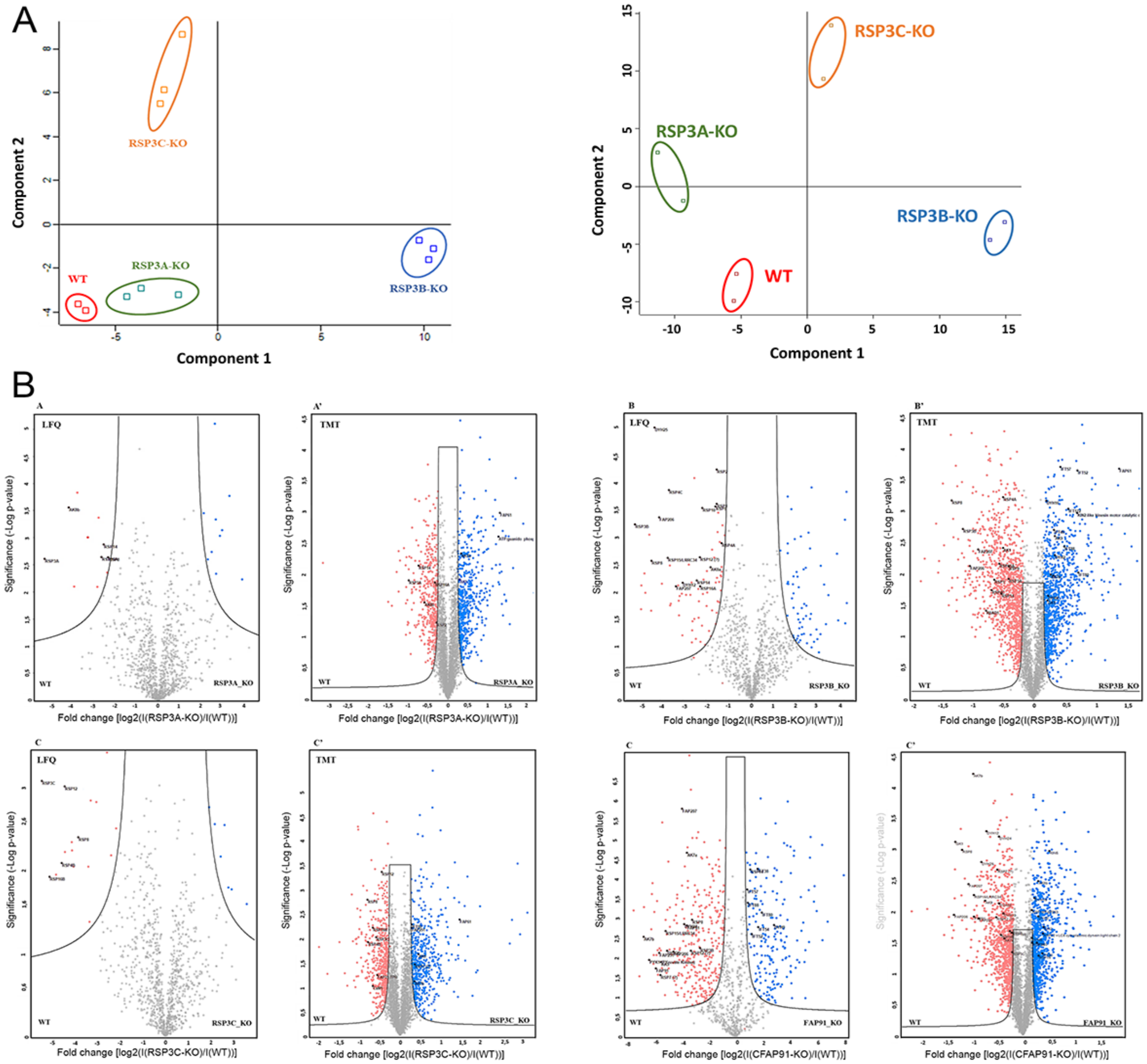

**Figure S8**. Graphical illustration of quantitative proteomics data. (A) Principal component analysis (PCA) score plot from fold-change values of the differentially expressed proteins showing similar grouping of ciliomes identified by mass spectrometry using either LFQ (graphs to the left) or TMT (graphs to the right) strategy based on two independent variables. The analyzed ciliary protein samples were color-coded as follows: WT - red, RSP3A-KO - green, RSP3B-KO - blue, and RSP3C-KO – orange. (B) Volcano plots of wild-type and RS mutant ciliary proteins identified by mass spectrometry using either LFQ (graphs to the left) or TMT (graphs to the right) strategy (N=3) showing changes in mutant ciliomes. FDR=0.05. Proteins Red dots represent proteins that are at a significantly lower level in RSP mutants compared to wild-type cells, while blue dots represent proteins that are more abundant in mutant cilia. The statistical significance of the difference in protein enrichment between wild-type and RS mutant ciliomes was calculated using Student’s t-test.

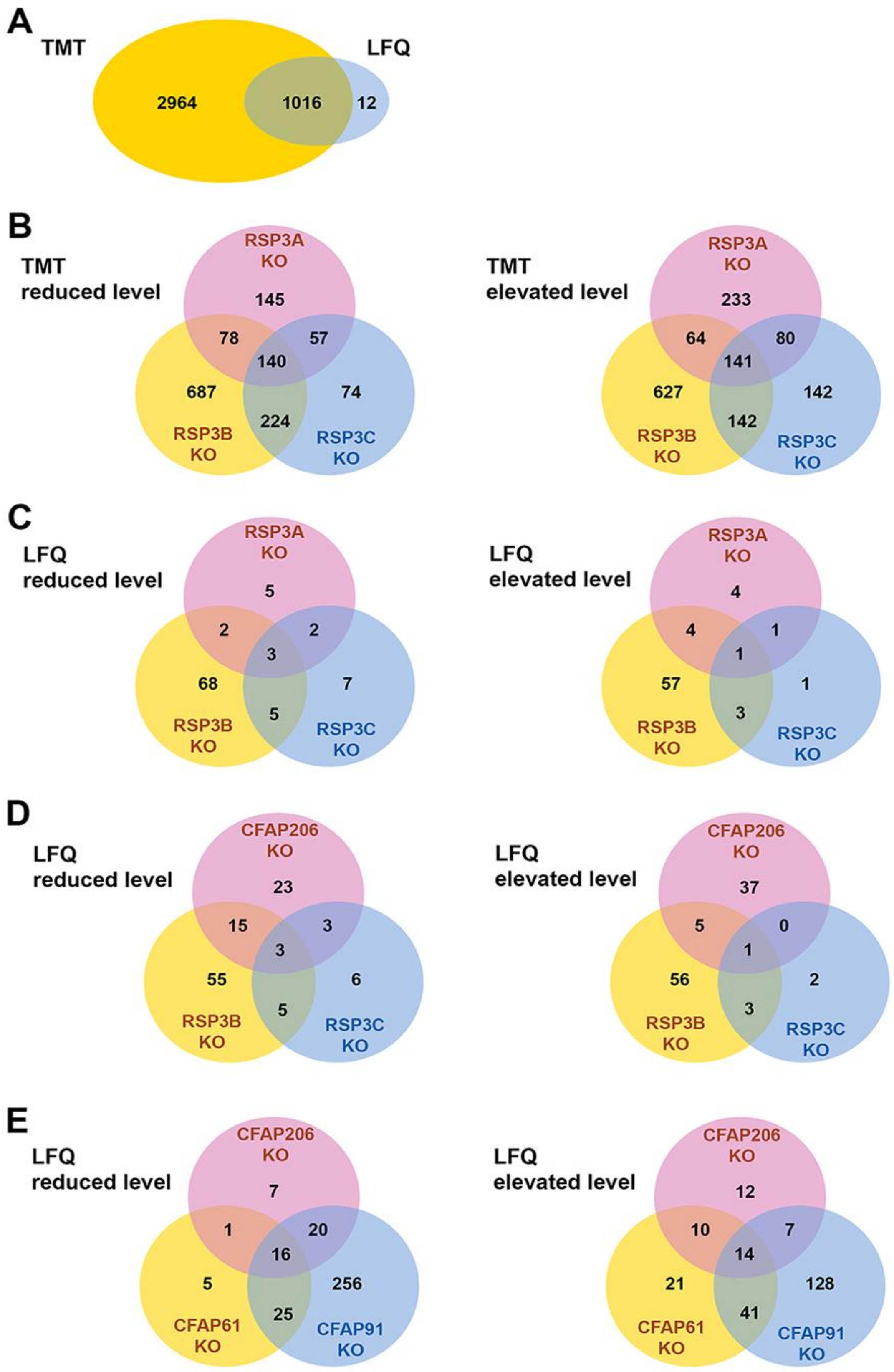

**Figure S9.** Venn diagrams indicating the overlap of differentially expressed proteins between different RS mutants based on ciliomes obtained either by LFQ or TMT approaches. (A) Comparison of the total number of proteins identified using LFQ and TMT approaches. (B-E) The overlap of proteins that were either reduced (to the left) or elevated (to the right) in different RS mutants as detected using (B) TMT and (C-E) LFQ methods. Images were prepared in Adobe Photoshop based on data sorted using Microsoft Excel program.

#### RSP1 and RSP10

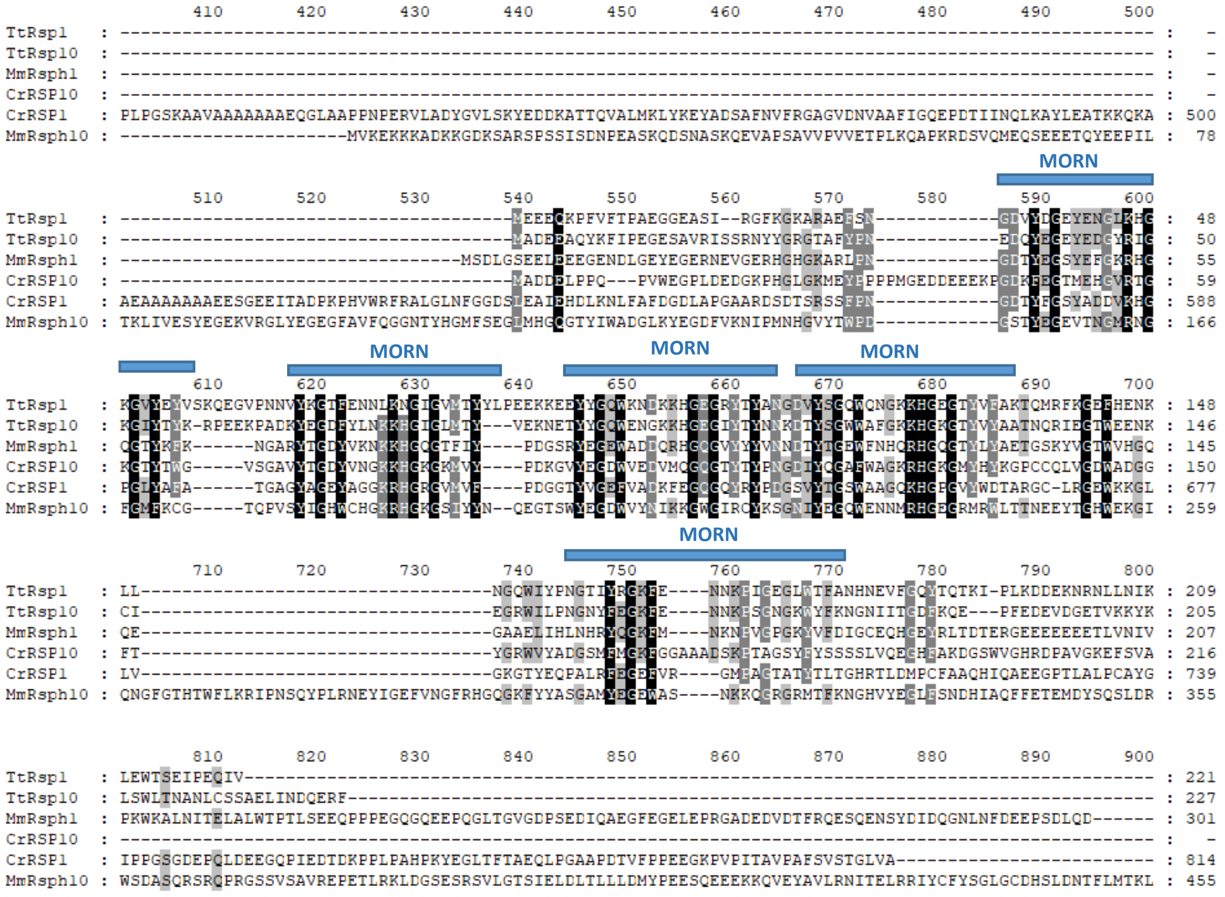

The following fragments: CrRSP1 M1-A400 and MmRsph10 H456-K876 are not shown.

Domain position in *Tetrahymena* orthologs.

| | domain | aa position | e-value |
|---|---|---|---|
| **TtRsp1** | **domain** | **aa position** | **e-value** |
| | MORN | 35-56 | 0.00000506 |
| | MORN | 64-85 | 0.172 |
| | MORN | 91-112 | 0.00000236 |
| | MORN | 114-135 | 1.95e-7 |
| | MORN | 160-181 | 2.35 |

| | domain | aa position | e-value |
|---|---|---|---|
| **TtRsp10** | **domain** | **aa position** | **e-value** |
| | MORN | 37-58 | 0.108 |
| | MORN | 65-86 | 0.0547 |
| | MORN | 89-110 | 0.0000134 |
| | MORN | 112-133 | 0.00298 |
| | MORN | 158-179 | 3.1 |

**RSP2**

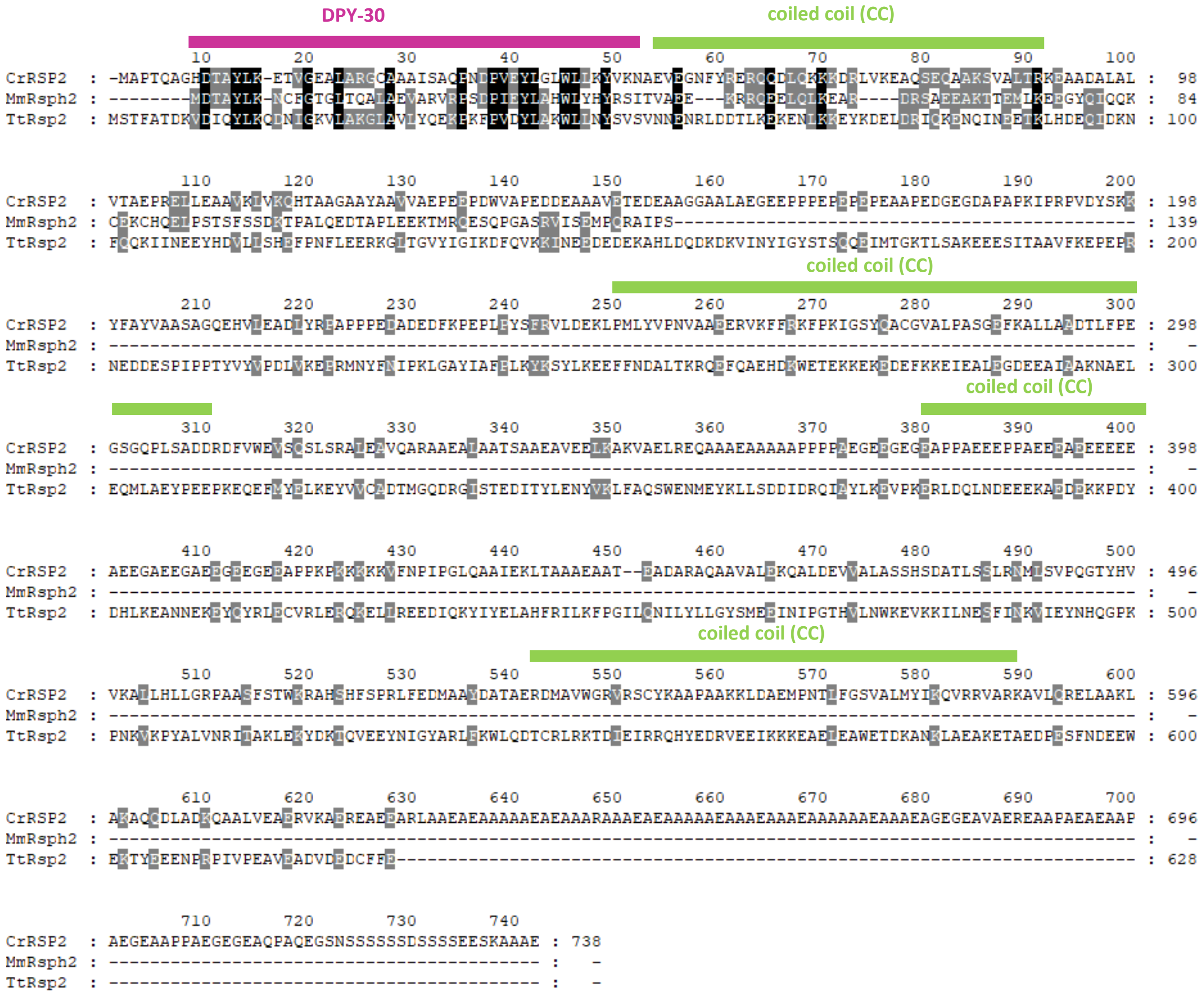

| TtRsp2 | domain | aa position | e-value |
|---|---|---|---|
| | DPY-30 | 10-51 | 9.6e-9 |
| | CC | 52-92 | n/a |
| | CC | 251-310 | n/a |
| | CC | 379-399 | n/a |
| | CC | 544-591 | n/a |

**RSP4/6**

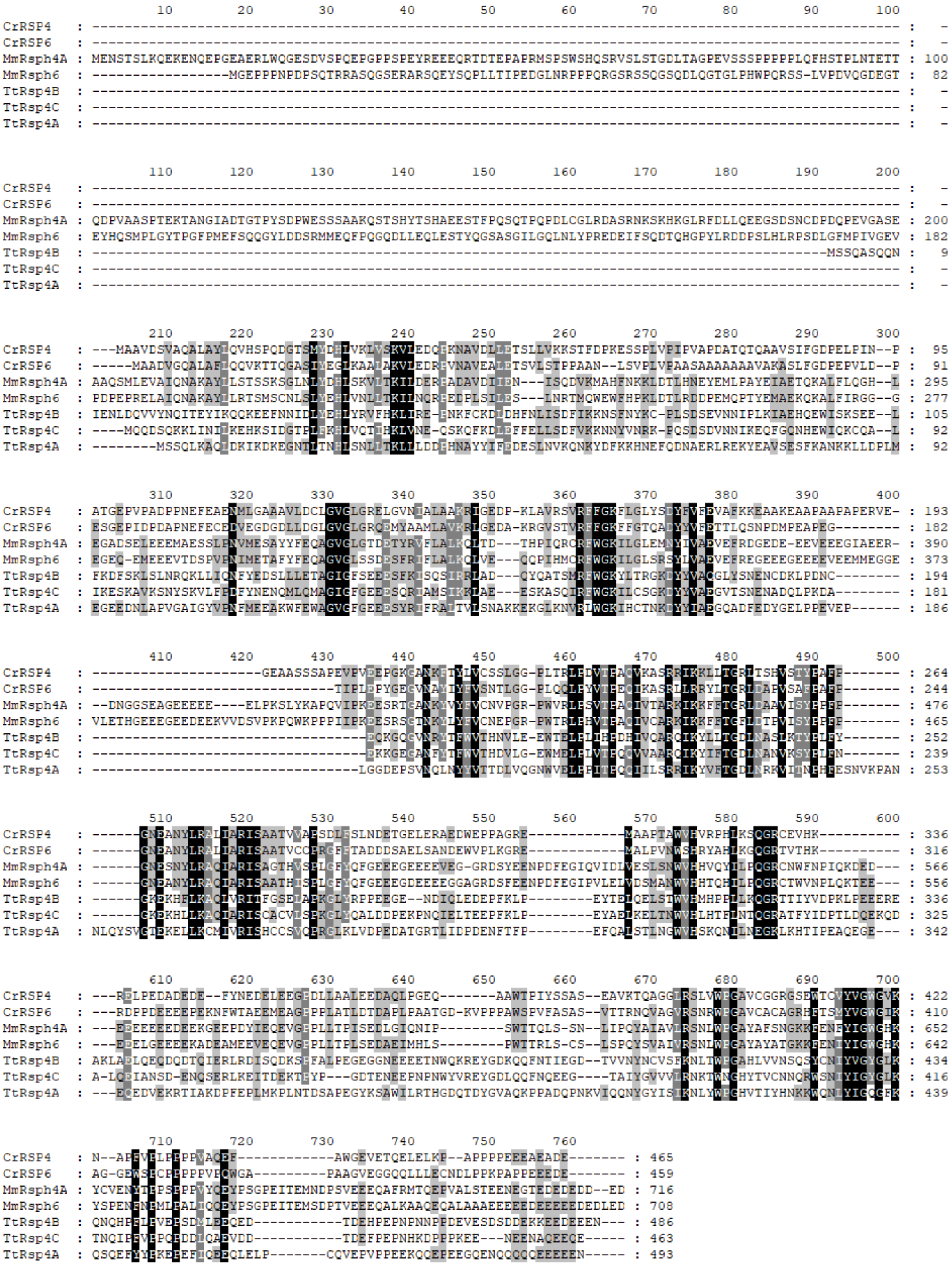

#### RSP7

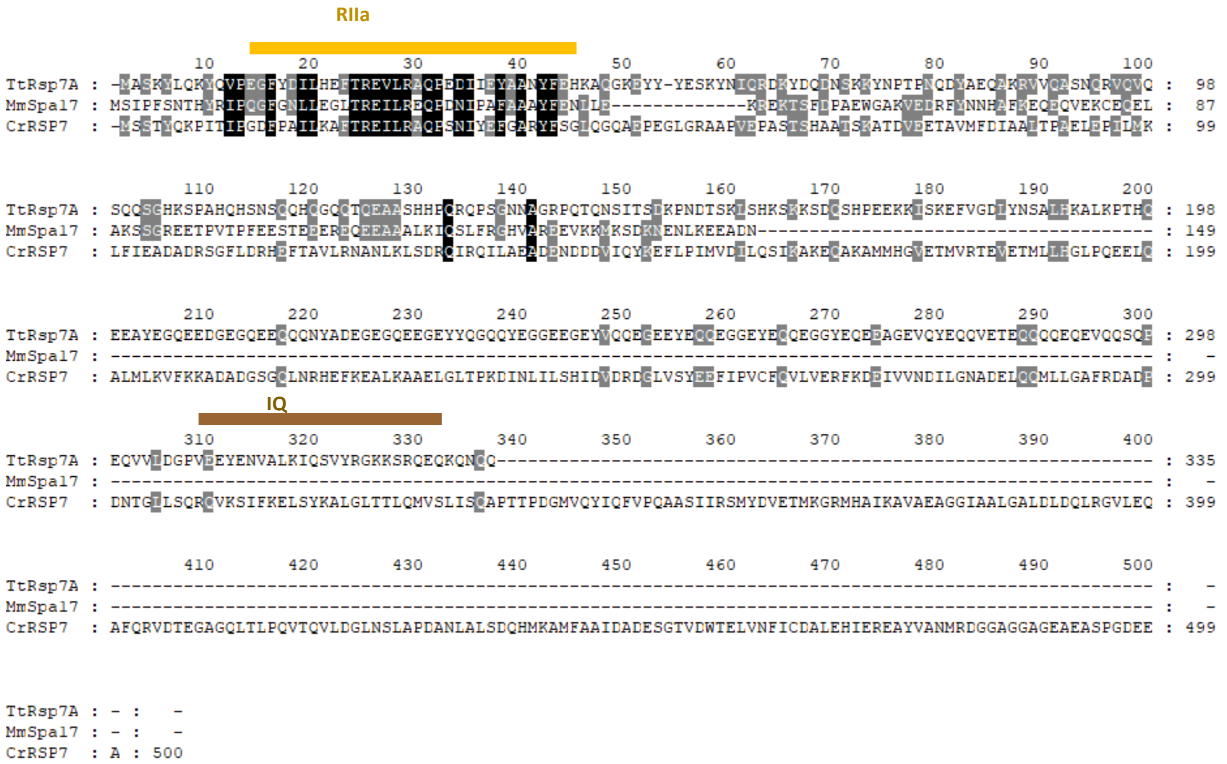

| TtRsp7A | domain | aa position | e-value |
|---|---|---|---|
| | RIIa | 13-45 | 1.4e-11 |
| | IQ | 309-331 | 6.06 |

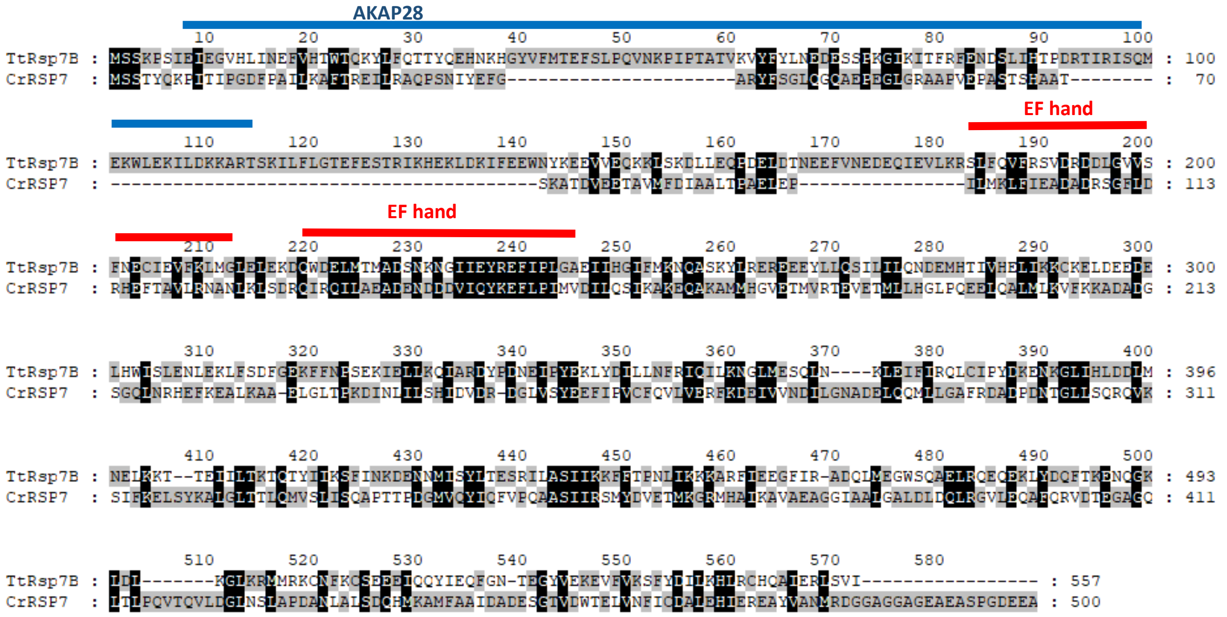

| TtRsp7B | domain | aa position | e-value |
|---|---|---|---|
| | AKAP28 | 8-115 | 5.9e-19 |
| | EF hand | 183-211 | 0.65 |
| | EF hand | 219-247 | 0.614 |

#### RSP8

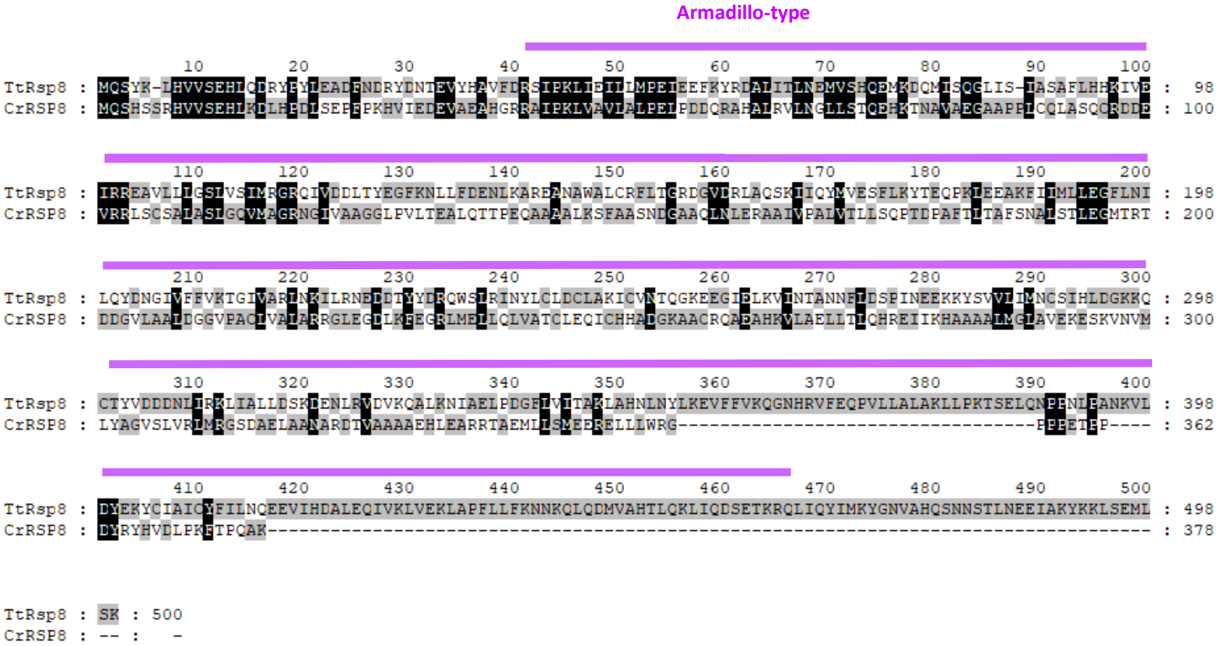

| TtRsp8 | domain | aa position | e-value |
|---|---|---|---|
| | Armadillo-type | 42-468 | n/a |

#### RSP9

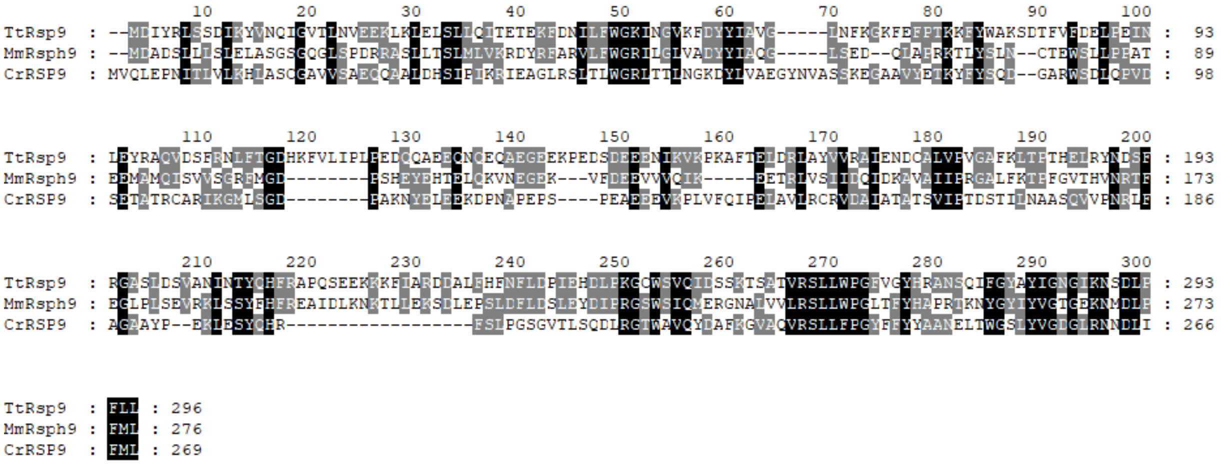

#### RSP11

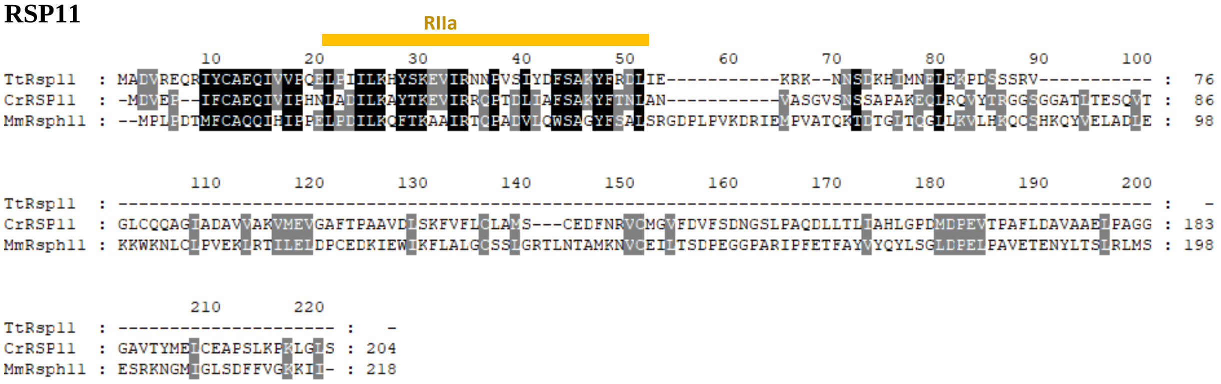

| TtRsp11 | domain | aa position | e-value |
|---|---|---|---|
| | RIIa | 21-56 | 2e-7 |

**RSP12**

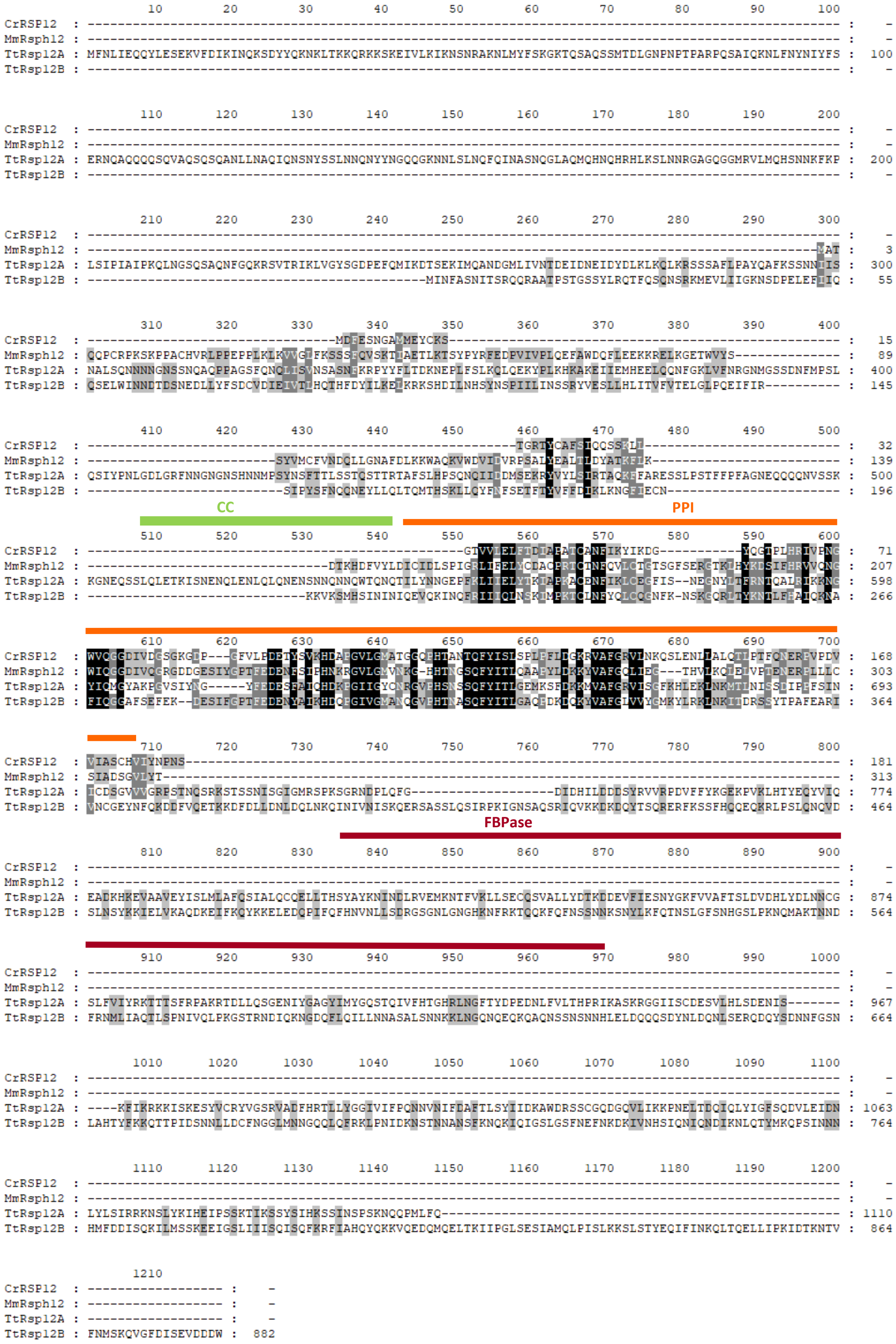

| | domain | aa position | e-value | |
|---|---|---|---|---|
| **TtRsp12A** | PPI | 544-697 | 1.4e-27 | |
| | FBPase | 809-941 | 3.8e-18 | |
| | CC | 509-542 | n/a | |
| | **TtRsp12B** | PPI | 209-369 | 6.3e-35 |

#### RSP14

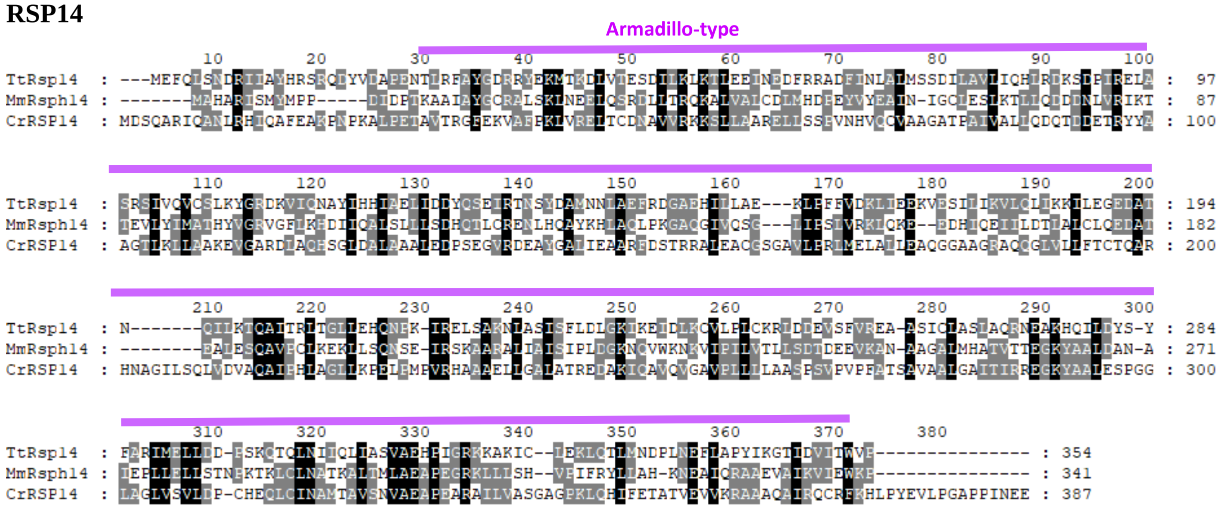

| TtRsp14 | domain | aa position | e-value |
|---|---|---|---|
| | Armadillo-type | 26-347 | n/a |

#### RSP15

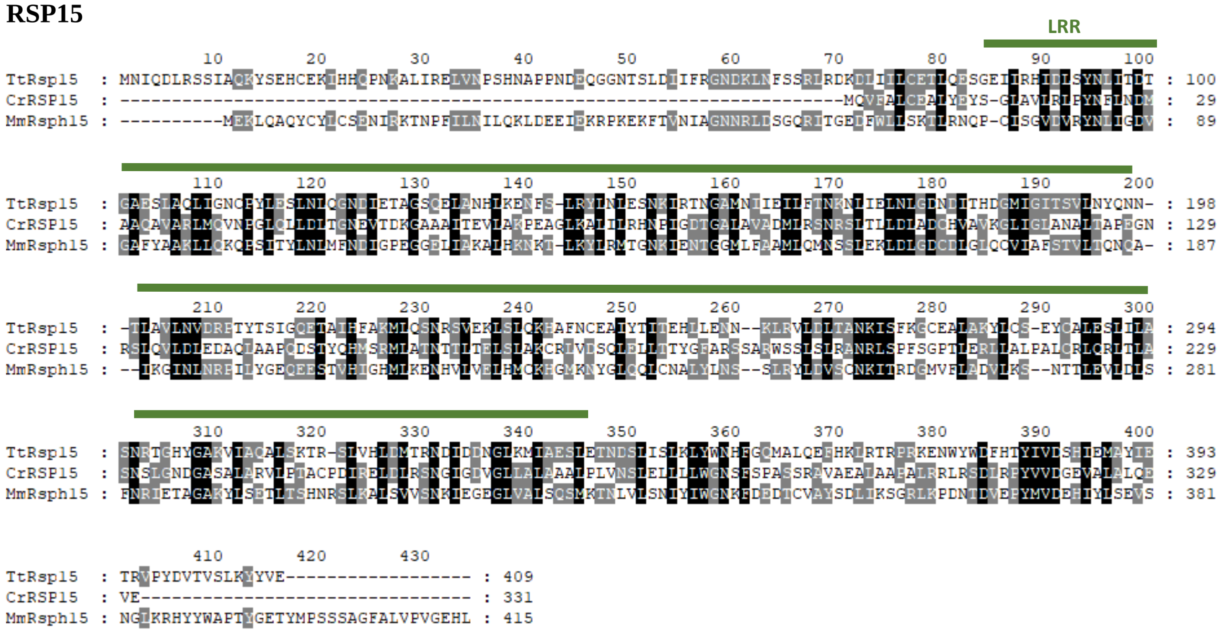

| TtRsp15 | domain | aa position | e-value |
|---|---|---|---|
| | LRR | 84-111 | 5.96 |
| | LRR | 112-139 | 1.73 |
| | LRR | 140-167 | 6.57 |
| | LRR | 168-195 | 1.78 |
| | LRR | 228-255 | 112 |
| | LRR | 256-283 | 1.33 |
| | LRR | 285-312 | 3.1 |
| | LRR | 313-340 | 5.06 |

#### RSP16

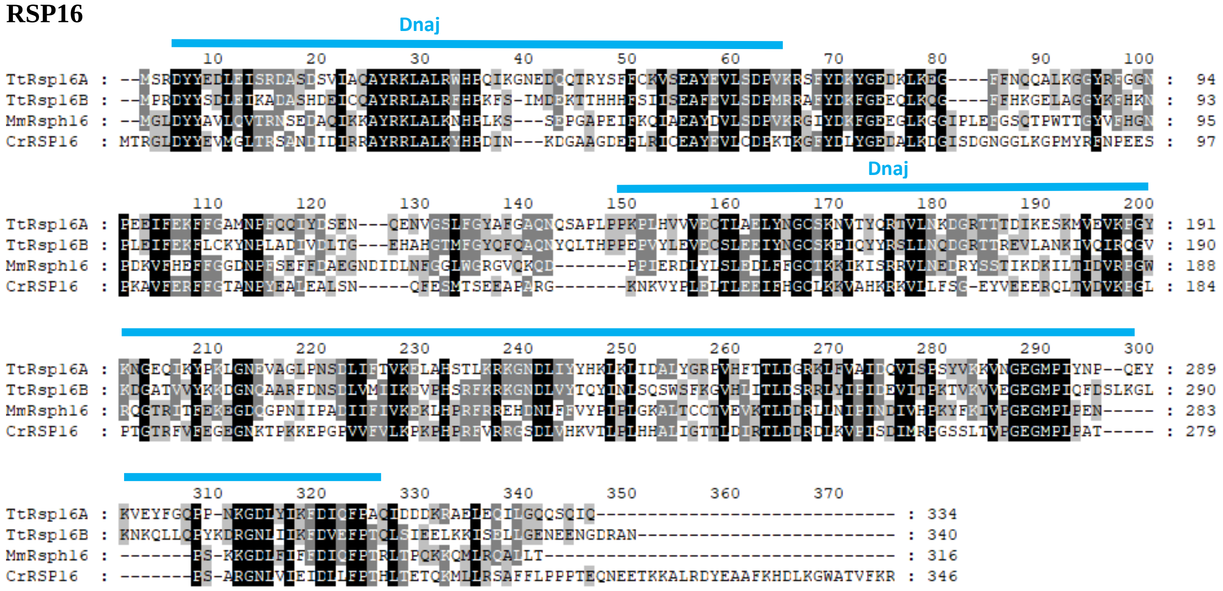

| | domain | aa position | e-value |
|---|---|---|---|
| **TtRsp16A** | Hsp40/Dnaj | 3-63 | 1.16e-14 |
| | Hsp40/Dnaj | 143-312 | 1.8e-36 |
| **TtRsp16B** | Hsp40/Dnaj | 3-62 | 1.31e-8 |
| | Hsp40/Dnaj | 142-314 | 1e-31 |

#### RSP20

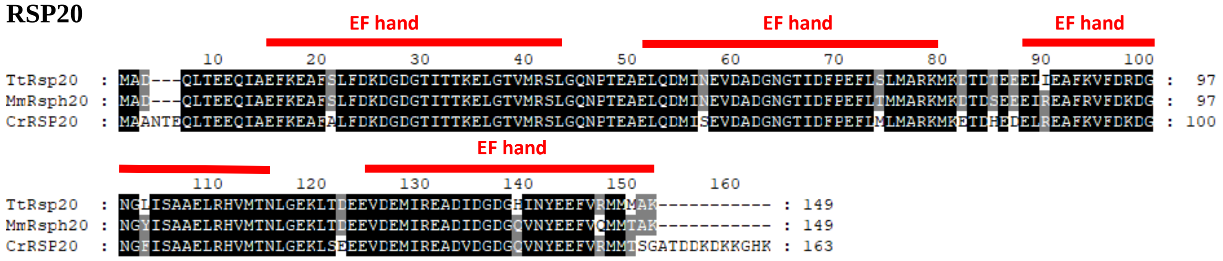

| **TtRsp20** | domain | aa position | e-value |
|---|---|---|---|
| | EF hand | 12-40 | 6.39e-9 |
| | EF hand | 48-76 | 2.39e-8 |
| | EF hand | 85-113 | 5.04e-7 |
| | EF hand | 121-149 | 3.67e-9 |

#### RSP22

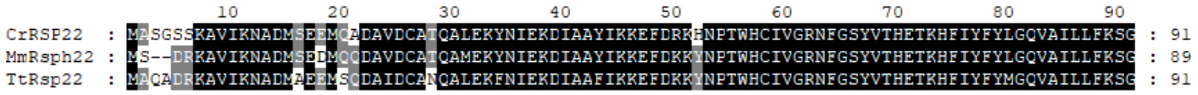

#### RSP23

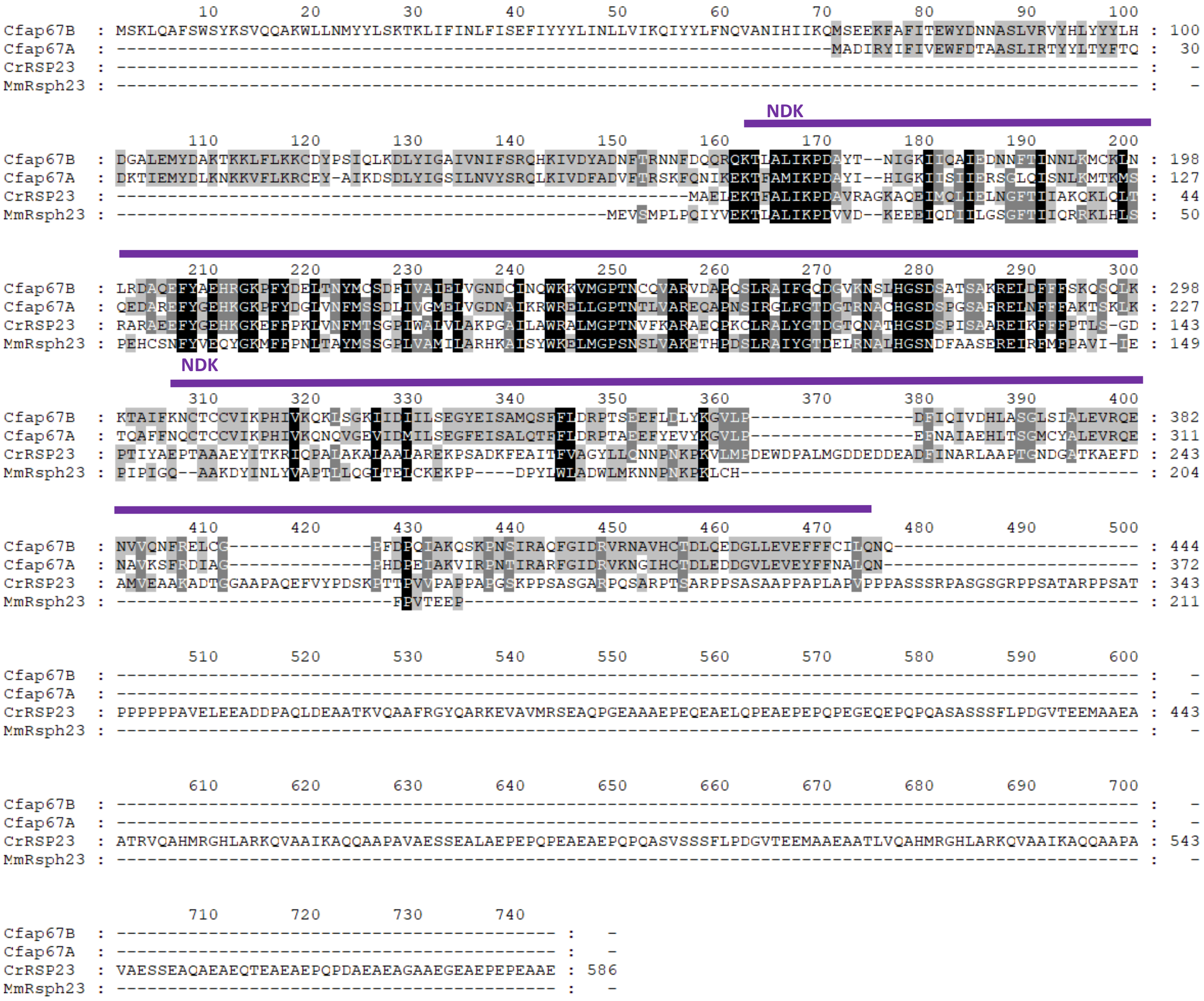

| TtRsp23 | domain | aa position | e-value |
|---|---|---|---|
| | NDK | 89-227 | 4.94e-68 |
| | NDK | 233-372 | 1.56e-38 |

#### FAP198

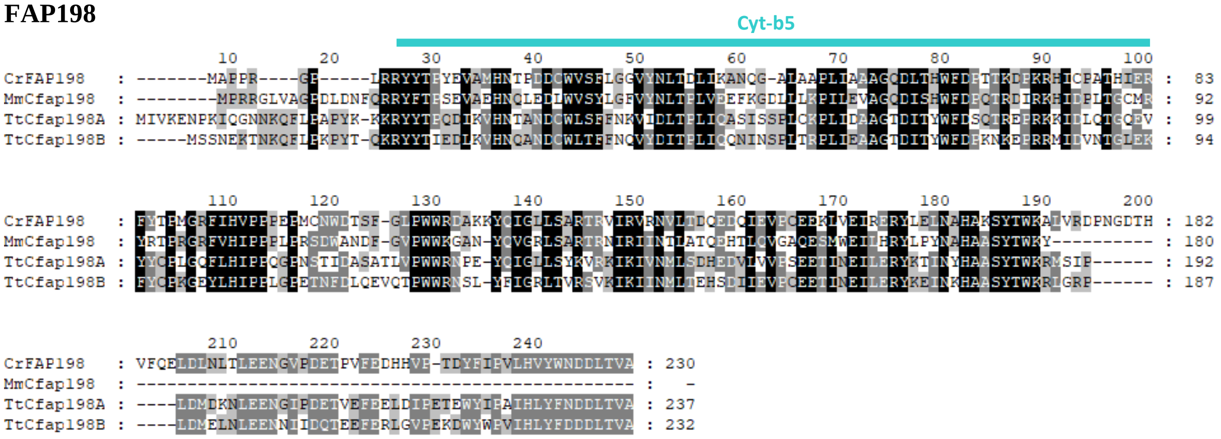

| | domain | aa position | e-value |
|---|---|---|---|
| **TtCfap198A** | Cyt-b5 | 27-99 | 0.0288 |
| **TtCfap198B** | Cyt-b5 | 22-103 | 0.0102 |

**FAP207**

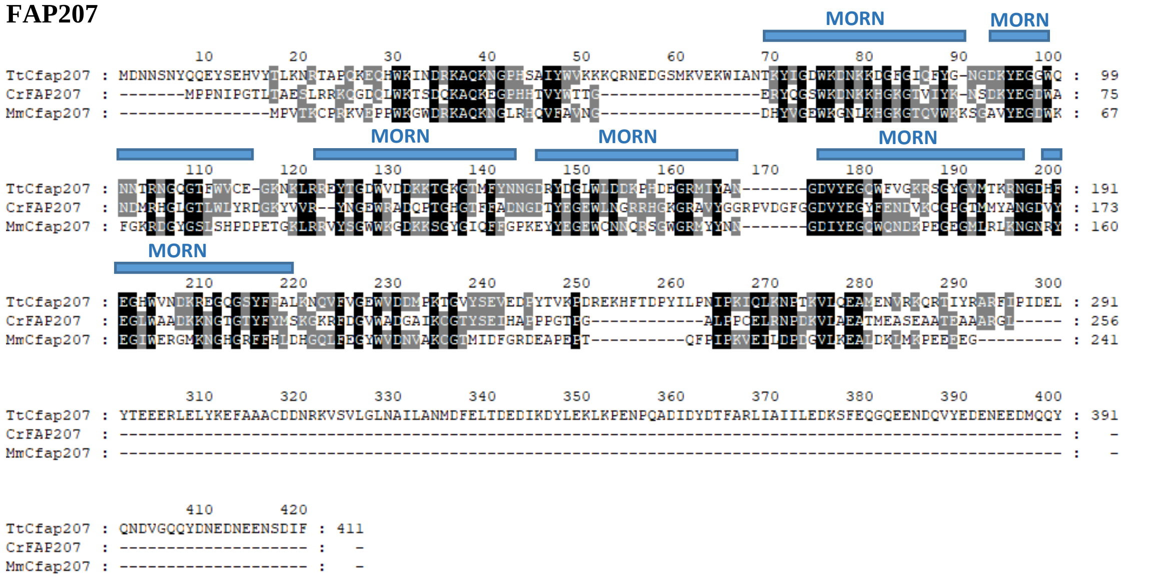

| **TtCfap207** | **domain** | **aa position** | **e-value** |
|---|---|---|---|
| | MORN | 69-90 | 0.0314 |
| | MORN | 92-113 | 0.148 |
| | MORN | 120-141 | 0.0274 |
| | MORN | 143-164 | 0.0293 |
| | MORN | 166-187 | 0.0000405 |
| | MORN | 189-210 | 0.22 |

**FAP253**

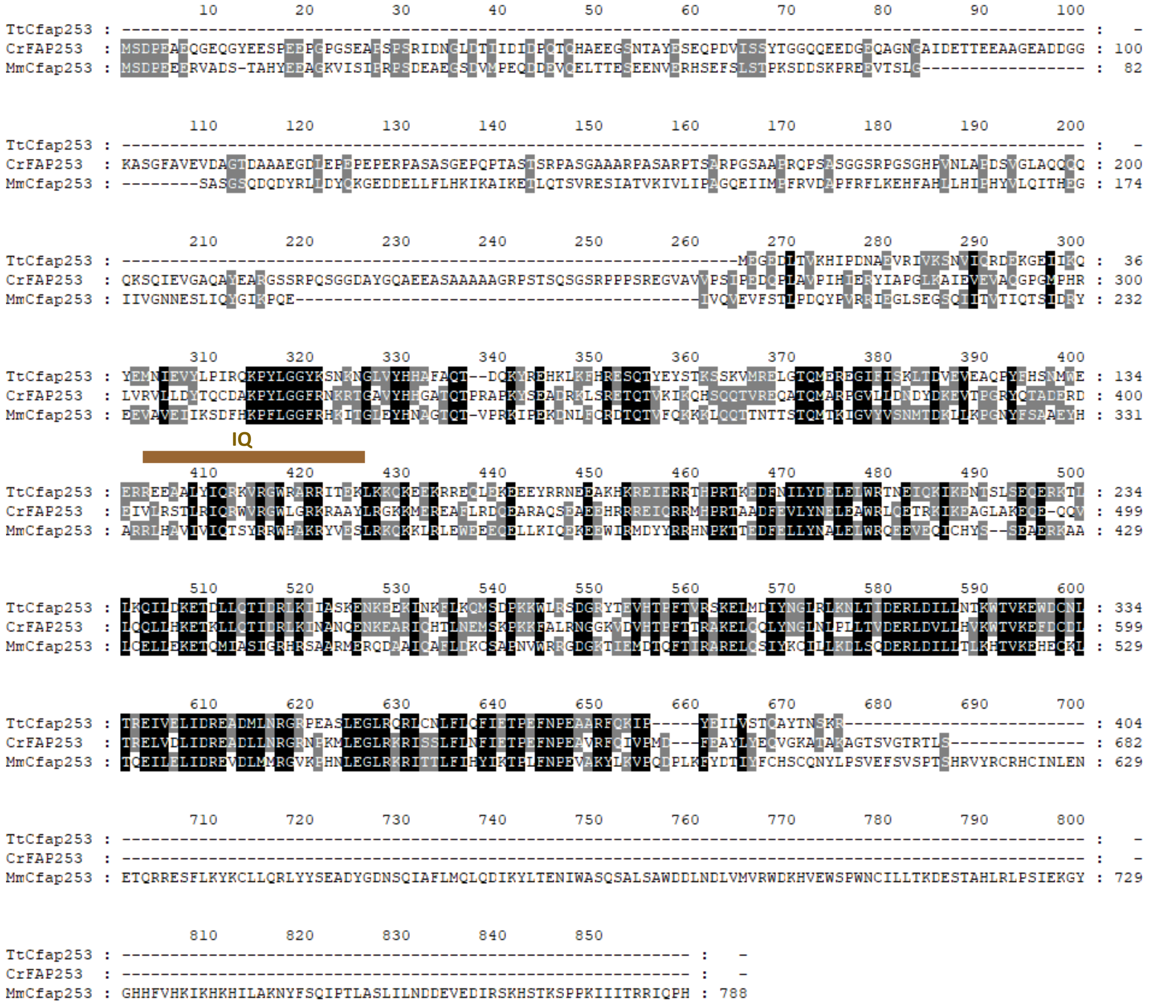

| **TtCfap253** | **domain** | **aa position** | **e-value** |
|---|---|---|---|
| | IQ | 136-158 | 1.3 |

**Figure S10**. Multiple alignments and domain analyses of RSP orthologs. Domain organization is predicted using SMART (http://smart.embl-heidelberg.de/) [1,2] or UniProt (htttps://www.uniprot.org) [11]. Domains are marked according to position in *Tetrahymena* proteins. The amino acid sequences of RSP orthologs were obtained from the NCBI protein database (National Center for Biotechnology Information). The accession numbers are provided in Table S3. The *Tetrahymena* Rsp orhologs were obtained from Tetrahymena Genome Database (https://tet.ciliate.org/). Protein amino acid sequences were aligned using ClustalX2 software [4] and edited using SeaView [5]. The identical and similar amino acid residues were shaded using GeneDoc [6] and the following color code: white letter on the black background (100% conserved residues); white letter on a grey background (80% conservation); black letter on a light grey background (60% conservation). The color lines indicate the position of indicated domains.

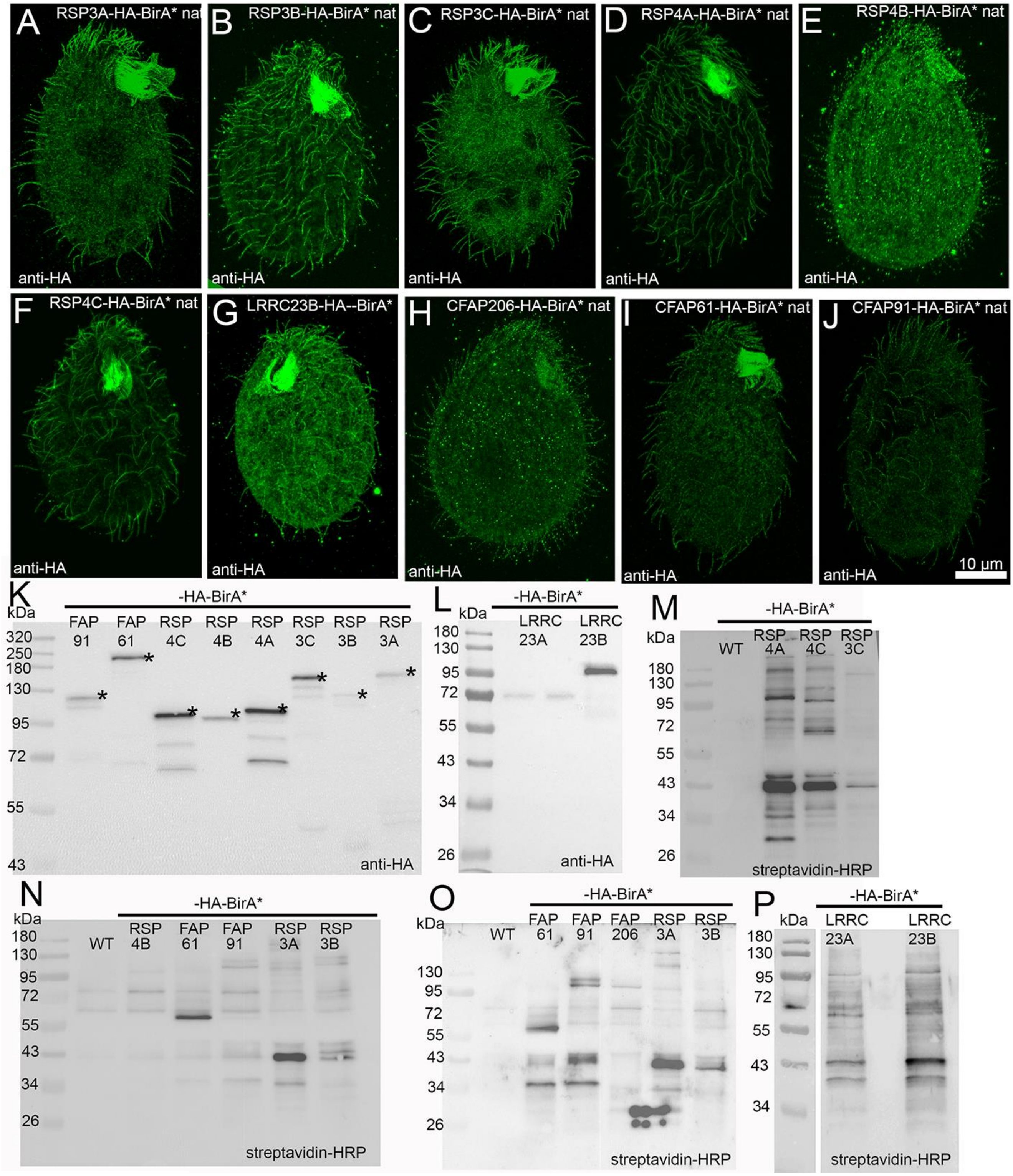

**Figure S11.** (A-J) Immunofluorescence confocal images of *Tetrahymena* cells expressing RS proteins as C-terminal –HA-BirA* fusions under the control of respective native promoters detected using anti-HA antibodies. (A) Rsp3A-HA-BirA*, (B) Rsp3B-HA-BirA*, (C) Rsp3C-HA-BirA*, (D) Rsp4A-HA-BirA*, (E) Rsp4B-HA-BirA*, (F) Rsp4C-HA-BirA*, (G) Lrrc23B-HA-BirA* (H) Cfap206-HA-BirA*, (I) Cfap61-HA-BirA*, (J) Cfap91-HA-BirA*. (K-P) Western blot analyses of the ciliary proteins isolated from *Tetrahymena* cells expressing RS proteins as –HA-BirA* fusions under the control of respective native promoters. A star indicates a band corresponding to the position of fusion proteins. (K-M) Western blot-based identification of the Rsp-HA-BirA* fusion proteins. (N-P) Western blot-based analyses of the biotinylated ciliary proteins in either WT cells or cells expressing Rsp-HA-BirA* under the control of the respective native promoter, all grown for 4 hrs in a 10 mM Tris-HCl buffer (pH 7.4) supplemented with biotin, detected using HRP-conjugated streptavidin.

### LRRC23

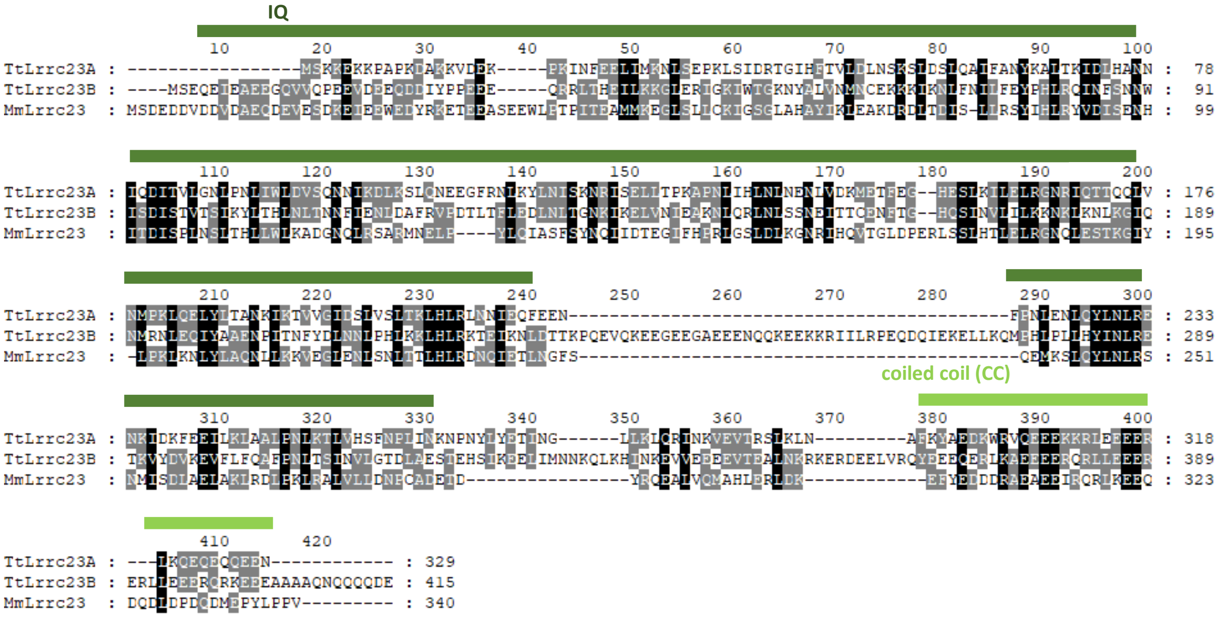

### STPG2

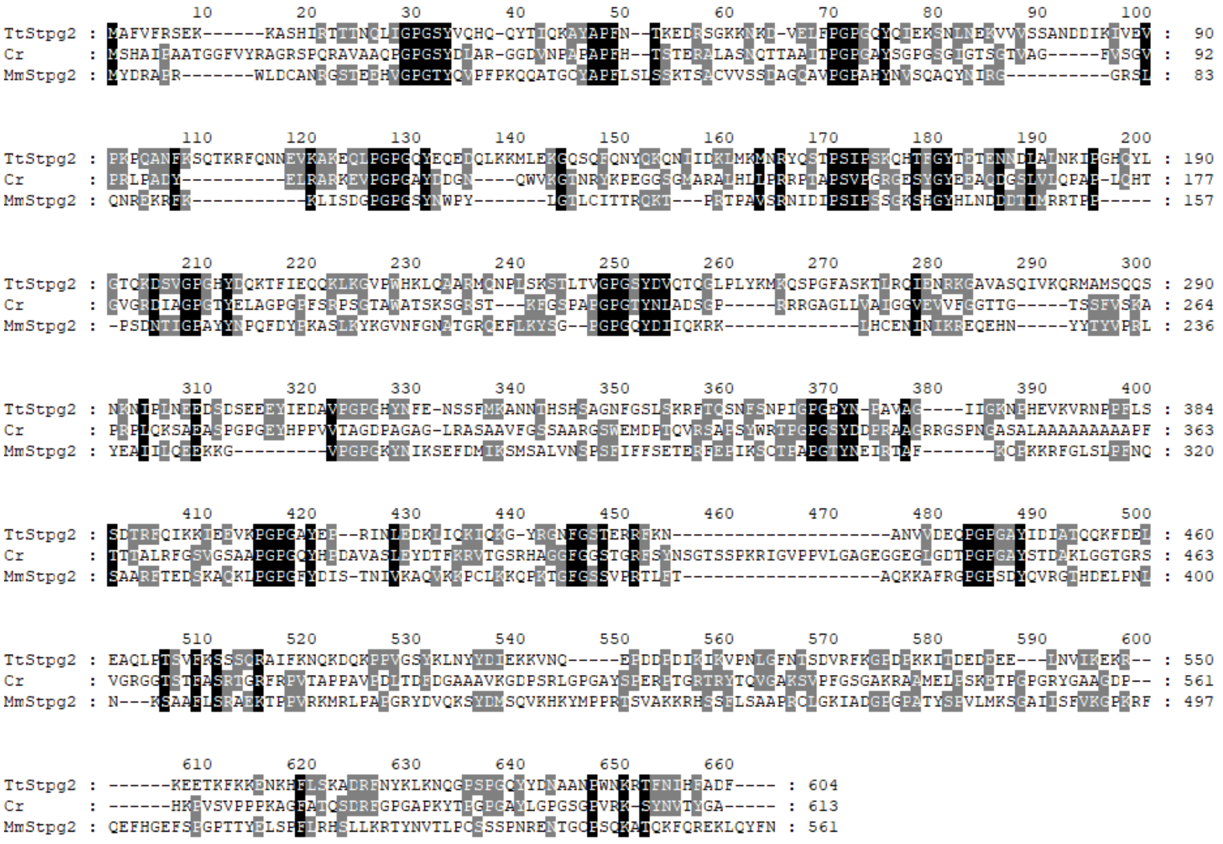

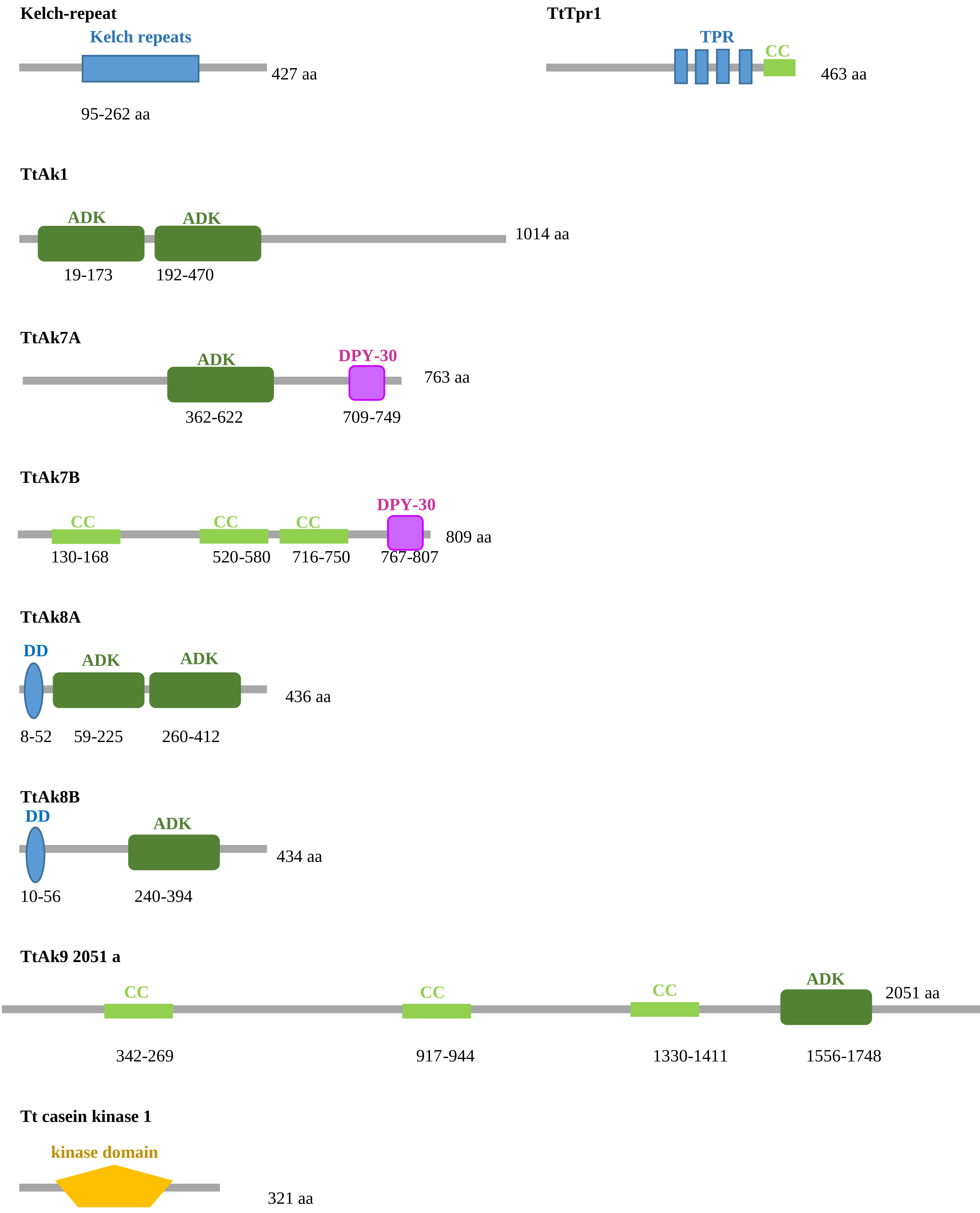
Kelch-repeat
Kelch repeats
427 aa
95-262 aa
TtTpr1
TPR
CC
463 aa
TtAk1
ADK
ADK
1014 aa
19-173
192-470
TtAk7A
ADK
DPY-30
763 aa
362-622
709-749
TtAk7B
CC
CC
CC
DPY-30
809 aa
130-168
520-580
716-750
767-807
TtAk8A
DD
ADK
ADK
436 aa
8-52
59-225
260-412
TtAk8B
DD
ADK
434 aa
10-56
240-394
TtAk9 2051 a
CC
CC
CC
ADK
2051 aa
342-269
917-944
1330-1411
1556-1748
Tt casein kinase 1
kinase domain
321 aa

**Figure S12**. Domain analyses of RS candidate proteins. Multiple alignments and domain analyses of RSP orthologs. Domain organization is predicted using SMART (http://smart.embl-heidelberg.de/) [1,2] or UniProt (htttps://www.uniprot.org) [11]. Domains are marked according to position in *Tetrahymena* proteins. The amino acid sequences of RSP orthologs were obtained from the NCBI protein database (National Center for Biotechnology Information). The accession numbers are provided in Table S3. The *Tetrahymena* Rsp orhologs were obtained from Tetrahymena Genome Database (https://tet.ciliate.org/). Protein amino acid sequences were aligned using ClustalX2 software [4] and edited using SeaView [5]. The identical and similar amino acid residues were shaded using GeneDoc [6] and the following color code: white letter on the black background (100% conserved residues); white letter on a grey background (80% conservation); black letter on a light grey background (60% conservation). The color lines indicate the position of indicated domains**.**

#### Supplementary Tables

**Table S1.** Data collection and analysis parameters for cryo-ET and subtomogram averaging.

Data collection and analysis parameters for cryo-ET and subtomogram averaging.

| **Method** | **Subtomogram Averaging** | | | |
|---|---|---|---|---|
| **Dataset** | ***WT*** | ***RSP3A-KO*** | ***RSP3B-KO*** | ***RSP3C-KO*** |
| **Microscope** | Titan Krios | Titan Krios | Titan Krios | Titan Krios |
| **Electron Detector** | Gatan K3 | Gatan K3 | Gatan K3 | Gatan K3 |
| **Zero-loss filter (eV)** | 20 | 20 | 20 | 20 |
| **Magnification** | 42,000 | 42,000 | 42,000 | 42,000 |
| **Voltage (keV)** | 300 | 300 | 300 | 300 |
| **Exposure (e/A$^2$)** | 160 | 160 | 80 | 160 |
| **Defocus (µm)** | 1.0-3.0 | 3.5-5.0 | 3-4 | 2.5-4.0 |
| **Pixel size (Å)** | 2.12 | 2.12 | 2.12 | 2.12 |
| **Tilt range (increment)** | -60° - 60° (3°) | -60° - 60° (3°) | -60° - 60° (3°) | -60° - 60° (3°) |
| **Tilt scheme** | dose symmetric | dose symmetric | dose symmetric | dose symmetric |
| **Tilt series acquired** | 58 | 37 | 50 | 38 |
| **Subtomo averaged** | 2608 | 2099 | 2093 | 2790 |
| **Symmetry imposed** | C1 | C1 | C1 | C1 |
| **Map resolution (Å)** | 19 | 20 | 22 | 17 |

**Table S2.** Global mass spectrometry analyses (LFQ) of wild-type and RS mutant showing total number of peptides.

| | | Number of total and unique peptides (X/Y) in RS knock-out mutant cilia | | | | | | | | | | | | | | | | | | | | | | | | |
|---|---|---|---|---|---|---|---|---|---|---|---|---|---|---|---|---|---|---|---|---|---|---|---|---|---|---|
| Protein | TGD accession numbers | WT1 | WT2 | RSP 3A-1 | RSP 3A-2 | RSP 3A-3 | RSP 3B-1 | RSP 3B-2 | RSP 3B-3 | RSP 3C-1 | RSP 3C-2 | RSP 3C-3 | WT1 | WT2 | WT3 | CFAP 206 | CFAP 206 | CFAP 206 | CFAP 91 | CFAP 91 | CFAP 91 | CFAP 61 | CFAP 61 | CFAP 61 | WT | RSP 2 |
| Rsp1 | TTHERM_000196029 | 0 | 20 | 17 | 24 | 21 | 13 | 14 | 15 | 20 | 20 | 21 | 19 | 20 | 21 | 15 | 14 | 14 | 12 | 10 | 13 | 17 | 17 | 17 | 15 | 13 |
| Rsp2 | TTHERM_00394410 | 45 | 46 | 29 | 27 | 31 | 24 | 30 | 24 | 27 | 37 | 36 | 50 | 46 | 47 | 36 | 31 | 32 | 40 | 48 | 40 | 46 | 49 | 53 | 88 | 9 |
| Rsp3A | TTHERM_01044600 | 21 | 15 | 0 | 0 | 0 | 7 | 10 | 9 | 11 | 14 | 16 | 17 | 17 | 15 | 15 | 8 | 13 | 18 | 18 | 17 | 18 | 18 | 22 | 25 | 24 |
| Rsp3B | TTHERM_00566810 | 22 | 24 | 28 | 25 | 29 | 0 | 0 | 0 | 27 | 26 | 24 | 40 | 37 | 36 | 26 | 20 | 23 | 19 | 16 | 15 | 31 | 30 | 30 | 52 | 53 |
| Rsp3C | TTHERM_00418270 | 43 | 41 | 26 | 25 | 35 | 30 | 35 | 33 | 0 | 0 | 0 | 46 | 42 | 43 | 4 | 2 | 4 | 32 | 30 | 22 | 49 | 45 | 47 | 61 | 41 |
| Rsp4A | TTHERM_00427590 | 36 | 35 | 25 | 22 | 23 | 24 | 26 | 27 | 23 | 30 | 30 | 50 | 42 | 49 | 32 | 37 | 35 | 39 | 38 | 36 | 44 | 46 | 47 | 75 | 53 |
| Rsp4B | TTHERM_00502580 | 16 | 18 | 6 | 6 | 8 | 9 | 14 | 13 | 1 | 3 | 2 | 15 | 14 | 13 | 8 | 8 | 8 | 12 | 11 | 11 | 17 | 13 | 16 | 29 | 24 |
| Rsp4C | TTHERM_00444180 | 23 | 25 | 15 | 10 | 15 | 5 | 5 | 5 | 17 | 23 | 20 | 29 | 25 | 27 | 22 | 19 | 23 | 25 | 26 | 25 | 29 | 27 | 28 | 34 | 24 |
| Rsp7A | TTHERM_00670610 | 10 | 10 | 9 | 10 | 7 | 10 | 8 | 9 | 8 | 9 | 8 | 17 | 15 | 18 | 9 | 8 | 8 | 6 | 6 | 5 | 15 | 14 | 13 | 40 | 45 |
| Rsp7B | TTHERM_00492840 | 15 | 15 | 8 | 7 | 10 | 8 | 12 | 12 | 9 | 10 | 11 | 12 | 11 | 11 | 9 | 7 | 9 | 1 | 1 | 0 | 13 | 9 | 12 | 43 | 30 |
| Rsp8 | TTHERM_00313520 | 15 | 14 | 5 | 3 | 6 | 1 | 1 | 1 | 1 | 2 | 2 | 8 | 9 | 10 | 2 | 1 | 2 | 2 | 2 | 1 | 13 | 11 | 11 | 35 | 25 |
| Rsp9 | TTHERM_00430020 | 15 | 14 | 18 | 17 | 19 | 12 | 11 | 11 | 20 | 20 | 15 | 23 | 25 | 22 | 17 | 19 | 18 | 16 | 17 | 19 | 21 | 23 | 19 | 54 | 41 |
| Rsp10 | TTHERM_00378600 | 25 | 23 | 10 | 10 | 12 | 7 | 11 | 11 | 12 | 15 | 12 | 23 | 20 | 21 | 18 | 18 | 20 | 21 | 19 | 19 | 23 | 24 | 24 | 33 | 30 |
| Rsp11 | TTHERM_00540090 | 5 | 4 | 3 | 2 | 3 | 3 | 4 | 2 | 2 | 5 | 4 | 5 | 5 | 7 | 2 | 2 | 1 | 2 | 4 | 2 | 6 | 5 | 6 | 8 | 4 |
| Rsp12A | TTHERM_01018330 | 24 | 20 | 15 | 18 | 12 | 7 | 9 | 11 | 1 | 1 | 1 | 26 | 24 | 23 | 6 | 6 | 8 | 2 | 2 | 0 | 22 | 23 | 24 | 16 | 3 |
| Rsp12B | TTHERM_00466090 | 8 | 2 | 0 | 1 | 0 | 0 | 0 | 0 | 3 | 5 | 3 | 7 | 4 | 3 | 5 | 2 | 4 | 3 | 5 | 5 | 9 | 8 | 9 | 2 | 0 |
| Rsp14 | TTHERM_000773538 | 17 | 18 | 8 | 3 | 6 | 6 | 11 | 9 | 10 | 17 | 14 | 18 | 15 | 15 | 12 | 14 | 14 | 15 | 12 | 12 | 13 | 13 | 15 | 29 | 29 |
| Rsp15 | TTHERM_00695730 | 13 | 14 | 7 | 5 | 8 | 0 | 0 | 0 | 4 | 9 | 8 | 13 | 9 | 10 | 0 | 0 | 0 | 1 | 1 | 0 | 11 | 12 | 12 | 14 | 10 |
| Rsp16A | TTHERM_00471260 | 30 | 30 | 15 | 15 | 13 | 16 | 18 | 16 | 16 | 25 | 23 | 28 | 25 | 29 | 25 | 22 | 20 | 30 | 29 | 29 | 32 | 34 | 36 | 41 | 35 |
| Rsp16B | TTHERM_00238810 | 21 | 17 | 18 | 21 | 22 | 19 | 16 | 17 | 8 | 7 | 5 | 19 | 22 | 19 | 8 | 9 | 9 | 15 | 16 | 13 | 19 | 18 | 17 | 24 | 18 |
| Rsp20/CaM1 | TTHERM_00630500 | 3 | 2 | 1 | 1 | 0 | 1 | 1 | 1 | 0 | 0 | 2 | 1 | 1 | 1 | 2 | 2 | 1 | 6 | 7 | 4 | 2 | 1 | 2 | 40 | 39 |
| Rsp22 | TTHERM_000649439 | 9 | 10 | 7 | 10 | 8 | 7 | 7 | 7 | 6 | 8 | 8 | 17 | 11 | 13 | 9 | 8 | 10 | 9 | 7 | 8 | 12 | 10 | 14 | 21 | 13 |
| Rsp23/Cfap67A | TTHERM_000372529 | 18 | 16 | 12 | 13 | 12 | 12 | 14 | 12 | 11 | 15 | 15 | 18 | 15 | 14 | 14 | 18 | 17 | 13 | 13 | 13 | 14 | 13 | 13 | 39 | 25 |
| Cfap61 | TTHERM_00641200 | 61 | 64 | 45 | 42 | 51 | 54 | 59 | 59 | 49 | 64 | 51 | 64 | 53 | 59 | 56 | 54 | 51 | 11 | 9 | 6 | 0 | 0 | 0 | 90 | 76 |
| Cfap251 | TTHERM_01262850 | 51 | 47 | 41 | 41 | 46 | 37 | 37 | 44 | 42 | 45 | 41 | 47 | 45 | 51 | 39 | 40 | 43 | 0 | 0 | 0 | 46 | 49 | 45 | 54 | 53 |
| Cfap91 | TTHERM_00578560 | 33 | 34 | 28 | 27 | 32 | 33 | 26 | 29 | 31 | 33 | 23 | 36 | 27 | 33 | 25 | 21 | 26 | 0 | 0 | 0 | 34 | 35 | 40 | 52 | 42 |
| Cfap206 | TTHERM_00820660 | 31 | 35 | 20 | 21 | 24 | 7 | 2 | 4 | 19 | 21 | 17 | 28 | 28 | 27 | 0 | 0 | 0 | 1 | 1 | 1 | 36 | 34 | 34 | 53 | 40 |
| Cfap253 | TTHERM_00316930 | 26 | 25 | 12 | 10 | 12 | 19 | 22 | 22 | 11 | 15 | 16 | 21 | 18 | 24 | 19 | 16 | 19 | 19 | 19 | 19 | 23 | 25 | 23 | 26 | 24 |
| Cfap207 | TTHERM_00529880 | 20 | 18 | 17 | 17 | 21 | 8 | 4 | 7 | 21 | 21 | 15 | 22 | 21 | 21 | 2 | 0 | 2 | 1 | 1 | 2 | 19 | 21 | 21 | 19 | 9 |
| Cfap198A | TTHERM_01092470 | 9 | 8 | 0 | 0 | 1 | 0 | 0 | 0 | 7 | 10 | 8 | 9 | 9 | 10 | 6 | 6 | 7 | 7 | 7 | 4 | 7 | 8 | 8 | 10 | 11 |
| Cfap198B | TTHERM_00476710 | 3 | 5 | 4 | 4 | 4 | 2 | 3 | 3 | 2 | 2 | 3 | 8 | 7 | 6 | 3 | 4 | 2 | 4 | 4 | 5 | 7 | 6 | 6 | 17 | 11 |
| TtTpr | TTHERM_00623040 | 11 | 12 | 3 | 6 | 10 | 8 | 11 | 9 | 0 | 0 | 0 | 10 | 9 | 10 | 1 | 1 | 2 | 5 | 5 | 5 | 9 | 10 | 9 | 19 | 20 |
| Kelch-motif | TTHERM_00760390 | 15 | 13 | 11 | 9 | 14 | 12 | 11 | 13 | 13 | 16 | 15 | 19 | 13 | 17 | 15 | 19 | 20 | 1 | 2 | 1 | 15 | 17 | 19 | 25 | 20 |
| Lrrc23A | TTHERM_000703669 | 11 | 11 | 11 | 12 | 6 | 9 | 12 | 10 | 5 | 9 | 11 | 11 | 13 | 13 | 12 | 7 | 10 | 0 | 0 | 1 | 10 | 10 | 11 | 23 | 24 |
| Lrrc23B | TTHERM_00105260 | 13 | 11 | 8 | 6 | 8 | 7 | 7 | 8 | 5 | 10 | 9 | 15 | 14 | 13 | 10 | 7 | 9 | 0 | 1 | 0 | 12 | 11 | 13 | 31 | 24 |
| Lrrc | TTHERM_01084360 | 14 | 10 | 5 | 3 | 6 | 1 | 0 | 0 | 6 | 3 | 4 | 9 | 8 | 10 | 2 | 4 | 4 | 0 | 0 | 0 | 2 | 5 | 6 | 10 | 10 |
| Stpg1 | TTHERM_00549640 | 12 | 8 | 3 | 7 | 2 | 1 | 0 | 1 | 8 | 7 | 5 | 12 | 8 | 11 | 3 | 0 | 2 | 0 | 0 | 0 | 6 | 3 | 6 | 4 | 2 |
| Stpg2 | TTHERM_00420710 | 19 | 16 | 15 | 16 | 10 | 7 | 9 | 7 | 22 | 14 | 19 | 30 | 26 | 28 | 18 | 15 | 19 | 11 | 9 | 8 | 25 | 27 | 25 | 7 | 16 |
| Ak1 | TTHERM_00317200 | 78 | 76 | 59 | 50 | 64 | 54 | 59 | 55 | 61 | 69 | 61 | 84 | 73 | 80 | 61 | 60 | 65 | 59 | 68 | 62 | 83 | 78 | 79 | 175 | 152 |
| Ak7A | TTHERM_00558060 | 28 | 29 | 23 | 24 | 24 | 26 | 27 | 24 | 22 | 23 | 25 | 42 | 37 | 38 | 28 | 31 | 33 | 24 | 22 | 21 | 35 | 35 | 38 | 67 | 74 |
| Ak7B | TTHERM_000569069 | 18 | 21 | 13 | 15 | 13 | 12 | 15 | 14 | 14 | 17 | 16 | 19 | 21 | 19 | 15 | 15 | 17 | 1 | 0 | 0 | 16 | 16 | 17 | 35 | 28 |
| Ak8A | TTHERM_00227800 | 36 | 35 | 27 | 26 | 31 | 21 | 23 | 24 | 26 | 33 | 30 | 40 | 39 | 38 | 28 | 29 | 27 | 27 | 29 | 29 | 35 | 31 | 33 | 78 | 68 |
| Ak8B | TTHERM_00455540 | 21 | 19 | 0 | 0 | 2 | 13 | 11 | 13 | 13 | 15 | 16 | 20 | 19 | 16 | 17 | 16 | 16 | 16 | 15 | 17 | 22 | 21 | 19 | 40 | 40 |
| Ak9 | TTHERM_00148750 | 84 | 89 | 55 | 53 | 59 | 65 | 70 | 67 | 54 | 68 | 62 | 94 | 82 | 88 | 74 | 61 | 58 | 1 | 0 | 0 | 77 | 81 | 75 | 137 | 109 |
| Ck1 | TTHERM_00938880 | 11 | 11 | 6 | 6 | 7 | 7 | 10 | 8 | 6 | 8 | 8 | 9 | 8 | 8 | 7 | 6 | 8 | 0 | 0 | 0 | 9 | 8 | 8 | 15 | 16 |
| PKA catalytic sub | TTHERM_00433420 | 7 | 6 | 8 | 7 | 5 | 2 | 6 | 3 | 4 | 5 | 11 | 23 | 19 | 20 | 12 | 12 | 12 | 10 | 9 | 11 | 21 | 22 | 21 | 18 | 30 |
| PKA catalytic sub | TTHERM_00658860 | 5 | 5 | 5 | 3 | 2 | 1 | 5 | 0 | 2 | 4 | 7 | 14 | 12 | 13 | 8 | 9 | 7 | 8 | 7 | 7 | 13 | 13 | 17 | 20 | 23 |
| PKA regulatory sub | TTHERM_00623090 | 36 | 32 | 34 | 34 | 23 | 7 | 5 | 5 | 26 | 29 | 36 | 64 | 63 | 68 | 46 | 39 | 42 | 25 | 26 | 24 | 55 | 52 | 54 | 61 | 76 |

**Table S3**. *Tetrahymena* RSP ortholog

| Protein | TGD accession numbers | UniProt / NCBI accession numbers | Bait<br>e-value | Length (residues) | Mass (kDa) | pI | Domains<br>in Tt orthologs | C.reinchardtii ortholog | C.reinchardtii UniProt/NCBI Accession number | M.musculus Ortholog | M.musculus UniProt/NCBI Accession number |
|---|---|---|---|---|---|---|---|---|---|---|---|
| Rsp1 | TTHERM_000196029<br>TTHERM_00196025 | W7X469 /<br>XP_012653365.1 | CrRSP10<br>1.78444e-19 | 221 | 26 | 6.3 | MORN | RSP1 | Q27YU0 /<br>XP_001693353.1 | Rsph1 | Q8VIG3 /<br>NP_079566.1 |
| Rsp2 | TTHERM_00394410 | Q233B2 /<br>XP_001011913.1 | MmRsph2<br>2.0127e-05 | 628 | 75 | 4.7 | DPY-30<br>CC | RSP2 | Q6UBQ3 /<br>XP_001702718.1 | Rsph2<br>Dydc2 | Q9D3X8 /<br>NP_081993.1 |
| Rsp3A | TTHERM_01044600 | Q22CF9 /<br>XP_001030633.2 | CrRSP3<br>2.61477e-19 | 764 | 90 | 4.9 | radial spoke 3 | RSP3 | A8J2J7 /<br>XP_001695406.1 | Rsph3b | Q9DA80 /<br>NP_001077414.1 |
| Rsp3B | TTHERM_00566810 | I7M308 /<br>XP_001022082.2 | CrRSP3<br>2.4593e-41 | 691 | 80 | 5.1 | Radial spoke 3 | | | | |
| Rsp3C | TTHERM_00418270 | Q22NW8 /<br>XP_001007288.2 | CrRSP3<br>8.91033e-20 | 950 | 111 | 5.3 | ARF<br>radial spoke 3 | | | | |
| Rsp4A | TTHERM_00427590 | Q23A99 /<br>XP_001013838.2 | CrRSP4<br>1.56162e-13<br>CrRSP6<br>2.18558e-12 | 493 | 57 | 4.9 | Radial spoke 4/6 | RSP4 | A8I550 /<br>XP_001700728.1 | Rsph4a | Q8BYM7 /<br>NP_001156429.1 |
| Rsp4B | TTHERM_00502580 | I7M9L5 /<br>XP_001022312.3 | CrRSP4<br>5.89158e-26<br>CrRSP6<br>2.55436e-20 | 486 | 57 | 4.7 | Radial spoke 4/6 | | | | |
| Rsp4C | TTHERM_00444180 | I7M3F3 /<br>XP_001023280.2 | CrRSP4<br>2.26135e-31<br>CrRSP6<br>3.55275e-33 | 463 | 54 | 4.9 | Radial spoke 4/6 | RSP6 | A8I554 /<br>XP_001700729.1 | Rsph6a | Q8CDR2/<br>NP_112545.2 |
| Rsp7A | TTHERM_00670610 | I7MMP3 /<br>XP_001026382.1 | CrRSP7<br>3.69931e-05 | 335 | 39 | 4.7 | RIIa<br>IQ | RSP7 | PNW81221.1<br>(A0A2K3DL29) | SPA17 | Q62252 /<br>NP_035579.1 |
| Rsp7B | TTHERM_00492840 | I7M9W2 /<br>XP_001023176.2 | CrRSP7<br>7.40409e-05 | 557 | 66 | 5.2 | AKAP28<br>Ef hands | | | | |
| Rsp8 | TTHERM_00313520 | Q22KC9 /<br>XP_001033533.2 | CrRSP8<br>4.83879e-16 | 500 | 58 | 6 | Armadillo | RSP8 | Q27YU6 /<br>XP_001701869.1 | nd | nd |
| Rsp9 | TTHERM_00430020 | Q231F8 /<br>XP_001011326.2 | CrRSP9<br>7.0382e-12 | 296 | 34 | 5 | nd | RSP9 | Q27YU5 /<br>XP_001690441.1 | Rsph9 | Q9D9V4 /<br>NP_083614.1 |
| Rsp10 | TTHERM_00378600 | Q23FF7<br>XP_001015441.2 | CrRSP10<br>1.90468e-19 | 227 | 27 | 5.3 | MORN | RSP10 | Q27YU4 /<br>XP_001702125.1 | Rsp10b | E9PYQ0 |
| Rsp11 | TTHERM_00540090 | I7MDI8 /<br>XP_001007934.2 | CrRSP11<br>1.85137e-11 | 76 | 9 | 7.9 | RIIa | RSP11 | Q27YU3 /<br>XP_001698630.1 | Rsph11<br>Ropn1l | Q9EQ00 /<br>NP_665851.2 |
| Rsp12A | TTHERM_01018330 | Q22XP3 /<br>XP_001010281.2 | CrRSP12<br>3.42243e-19 | 1110 | 126 | 9.3 | CC, PPI,<br>FBPase | RSP12 | A8I2U9 /<br>XP_001699890.1 | Rsph12<br>Ppil6 | Q9D6D8 /<br>NP_082706.1 |
| Rsp12B | TTHERM_00466090 | I7MIU8 /<br>XP_001025007.2 | CrRSP12<br>4.4317e-18 | 882 | 102 | 9.0 | PPI | | | | |
| Rsp14 | TTHERM_000773538<br>TTHERM_00773533 | W7XHA5 /<br>XP_012655051.1 | CrRSP14<br>0.535804 | 354 | 41 | 5.7 | Armadillo | RSP14 | A8HNV0 /<br>XP_001690282.1 | Rsph14 | Q9D3W1 /<br>NP_001157006.1 |
| Rsp15 | TTHERM_00695730 | Q24C99 /<br>XP_001025584.1 | CrRSP15<br>5.73734e-19 | 409 | 47 | 5.7 | LRR | RSP15 | A0A2K3DEQ5 /<br>PNW79021 | Rsph15<br>LRRC34 | Q9DAM1 /<br>NP_082217.1 |
| Rsp16A | TTHERM_00471260 | I7MGN3 /<br>XP_001033013.2 | CrRSP16<br>1.38908e-43 | 334 | 38 | 6.3 | Dnaj | RSP16 | A8IKR9 /<br>XP_001690875.1 | Rsph16<br>Dnajb13 | Q80Y75 /<br>NP_705755.2 |
| Rsp16B | TTHERM_00238810 | I7M3Y8 /<br>XP_001024806.2 | CrRSP16<br>4.01687e-40 | 340 | 39 | 6.2 | Dnaj | | | | |
| Rsp20/CaM1 | TTHERM_00630500 | Q241P0 /<br>XP_001022775.2 | CrRSP20<br>7.97936e-92 | 149 | 17 | 4 | EF hand | RSP20<br>calmodulin | A8IDP6 /<br>XP_001703420.1 | Rsph20<br>Calm1 | P0DP26 /<br>NP_033920.1 |

| | | | | | | | | | | | |
|---|---|---|---|---|---|---|---|---|---|---|---|
| Rsp22/LC8<br>Wrong number | TTHERM_000649439<br>TTHERM_00649435 | W7X4R1 /<br>XP_012653183.1 | CrRSP22<br>7.5e-56 | 91 | 10 | 6.8 | LC | RSP22<br>LC8 | A8JH45 /<br>XP_001702907.1 | Rsph22<br>Dynll2 | Q9D0M5 /<br>NP_080832.1 |
| Rsp23/<br>Cfap67A | TTHERM_000372529<br>TTHERM_00372533 | W7XGD1 /<br>XP_012651451.1 | CrRSP23<br>9.32902e-33 | 372 | 43 | 6.2 | NDK | FAP67 | Q69B19 /<br>XP_001698136.1 | Rsph23<br>Nme5 | Q99MH5 /<br>NP_542368.2 |
| Cfap61 | TTHERM_00641200 | Q23F13 /<br>XP_001015337.1 | CrFAP61<br>5.54686e-36 | 1699 | 198 | 5.4 | DUF4821, CC<br>FAD/NAD(P) binding | FAP61 | A8IF44 /<br>XP_001703513.1 | Cfap61 | Q8CEL2 /<br>XP_030108106.1 |
| Cfap251 | TTHERM_01262850 | Q24DE2 /<br>XP_001026044.2 | CrFAP251<br>3.4439e-75 | 996 | 109 | 5 | WD40 | FAP251 | A8IRK7 /<br>XP_001691834.1 | Cfap251<br>WDR66 | E9Q743 /<br>NP_001357769.1 |
| Cfap91 | TTHERM_00578560 | I7LWP7 /<br>XP_001022857.1 | CrFAP91<br>1.59648e-19 | 644 | 76 | 7.3 | PaaSYMP<br>CC | FAP91 | A8IH47 /<br>XP_001690436.1 | Cfap91<br>MAATS1 | Q8BRC6<br>NM_001081025.1 |
| Cfap206 | TTHERM_00820660 | Q23H79 /<br>XP_001016174.1 | MmCfap206<br>3.71623e-63 | 635 | 73 | 6 | FAP206 domain | FAP206 | A0A2K3DUY6 /<br>PNW84338.1 | CFap206 | Q6PE87 /<br>NP_001074494.1 |
| Cfap253 | TTHERM_00316930 | I7M989 /<br>XP_001021374.2 | CrFAP253<br>1.1375e-102 | 404 | 49 | 9 | IQ | FAP253 | A0A2K3D359 /<br>PNW74974.1 | Iqub | Q8CDK3 /<br>NP_766123.2 |
| Cfap207 | TTHERM_00529880 | I7MCW2 /<br>XP_001032699.2 | CrFAP207<br>2.1404e-46 | 411 | 48 | 4.9 | MORN | FAP207 | A0A2K3DJP7 /<br>PNW80755.1 | Morn3 | Q8C5T4 /<br>NP_083388.1 |
| Cfap198A | TTHERM_01092470 | Q24BN4 /<br>XP_001025454.2 | CrFAP198<br>1.34852e-59 | 237 | 28 | 5 | Cyt-b5 | FAP198 | A0A2K3DCN8 /<br>PNW78294.1 | Cyb5d1 | Q5NCY3 /<br>NP_001038990.1 |
| Cfap198B | TTHERM_00476710<br>TTHERM_00476715 | I7MEL3 /<br>XP_001017378.2 | CrFAP198<br>2.05639e-56 | 232 | 17 | 8 | Cyt-b5 | | | | |
| TtTpr | TTHERM_00623040 | Q240Y0 /<br>XP_001022529.2 | - | 463 | 54 | 6 | TPR | nd | nd | nd | nd |
| Kelch-motif | TTHERM_00760390 | I7M609 /<br>XP_001031655.2 | - | 427 | 48 | 5.3 | Kelch repeat | nd | nd | nd | nd |
| Lrrc23A | TTHERM_000703669<br>TTHERM_00703665 | W7X6G9 /<br>XP_012655513.1 | MmLrrc23<br>5.55138e-27 | 329 | 38 | 8.4 | LRR | nd | nd | Lrrc23 | O35125 /<br>NP_001289484.1 |
| Lrrc23B | TTHERM_00105260 | Q234H2 /<br>XP_001012276.1 | MmLrrc23<br>2.11729e-11 | 415 | 49 | 4.98 | LRR | nd | nd | | |
| Lrrc | TTHERM_01084360 | Q22BT2 /<br>XP_001030431.1 | MmLrrc74B<br>9.70234e-18 | 1014 | 117 | 6.4 | LRR | nd | nd | Lrrc74B | Q14BP6<br>NP._001138907.1 |
| Lrrc | TTHERM_00046820 | Q23DK9<br>XP_001014677.2 | - | 401 | 46 | 5.6 | LRR | nd | nd | nd | nd |
| RIIa domain-containing protein 1 | TTHERM_00537370 | I7MLY6<br>XP_001023547.1 | Mm RIID<br>2.1451e-13 | 137 | 16 | 5.8 | DD-RIIAD1 | nd | nd | RIIa domain-containing protein 1 | Q3KNY5.1<br>NP_001404721.1 |
| MRNN04 | TTHERM_00324550 | Q237E4<br>XP_001013042.2 | MmMORN5<br>1.01265e-39 | 197 | 23 | 7 | MORN | nd | nd | MORN5 | Q9DAI9<br>NP_083585.1 |
| Stpg1 | TTHERM_00549640 | I7M0B4 /<br>XP_976712.2 | - | 349 | 39 | 9.8 | PGP motif | nd | nd | nd | nd |
| Stpg2 | TTHERM_00420710 | I7M6R0 /<br>XP_001033326.2 | MmStpg2<br>1.7267e-05 | 604 | 69 | 9.4 | PGP motif | CHLRE_09g41<br>5650v5 | A0A2K3DFZ1<br>XP_001696826.2 | Stpg2 | Q8C8J0<br>NP._941061.1 |
| adenylate kinase 1, Ak1 | TTHERM_00317200 | I7M2R5 /<br>XP_001021401.2 | Cr<br>5.58533e-48 | 1014 | 119 | 5.4 | ADK | CHLRE_01g02<br>9750v5 | A0A2K3E6L7<br>XP_042928532.1 | Ak1 | Q9R0Y5 /<br>NP_001185719.1 |
| adenylate kinase 7A, Ak7A | TTHERM_00558060 | I7MLH8 /<br>XP_001022370.2 | MmAk7<br>3.68184e-72 | 763 | 88 | 5 | ADK<br>DPY-30 | - | - | Ak7 | Q9D2H2 |
| adenylate kinase 7B, Ak7B | TTHERM_000569069 | W7WZL8 /<br>XP_012655235.1 | MmAk7<br>5e-22 | 809 | 95 | 4.6 | DPY-30<br>CC | | | | |
| adenylate kinase 8A, Ak8A | TTHERM_00227800 | Q23BQ2 /<br>XP_001014311.2 | MmAk8<br>7.20501e-37 | 436 | 50 | 6.6 | DD<br>ADK | nd | nd | Ak8 | Q32M07 /<br>NP_001029046.2 |
| adenylate kinase 8B, Ak8B | TTHERM_00455540 | I7MA87 /<br>XP_001024160.1 | MmAK8<br>5.04591e-17 | 434 | 51 | 8.5 | DD<br>ADK | | | | |
| adenylate kinase 9, Ak9 | TTHERM_00148750 | I7LWA5 /<br>XP_001021510.2 | MmAk9<br>8.11956e-41 | 2051 | 240 | 5.6 | CC<br>ADK | nd | nd | Ak9 | G3UYQ4<br>NP._001357742.1 |

| | | | | | | | | | | | |
|---|---|---|---|---|---|---|---|---|---|---|---|
| caseine kinase 1, Ck1, | TTHERM_00938880 | Q22DQ4 / XP_001031063.1 | MmCk1 1.69772e-139 | 321 | 38 | 6 | Kinase domain | CHLRE_12g549750v5 | A8iYG8 XP_001693999.1 | Ck1 | Q07954 NP._620690.1 |
| Guanylate kinase, Gk1 | TTHERM_00781030 | I7MMN2 XP_001026259.1 | MmGuk1 6.7636e-40 | 264 | 31 | 5.9 | GuKc | CHLRE_09g394102v5 | A0A2K3DEF1 XP_042921222.1 | Guk1 | Q64520 NP_001390764.1 |
| Phosphodiesterase PDE | TTHERM_00293350 | I7M0W2 XP_001013067.3 | MmPDE 5.6219e-70 | | 111 | 6.3 | | CHLRE_13g605100v5 | A0A2K3D1E3 XP_042917831.1 | PDE1 | A2ASF9 NP_001342073.1 |
| PKA catalytic subunit-1 | TTHERM_00433420 | Q231B5 / XP_001011369.2 | Cr 2.23002e-94 | 319 | 36 | 9 | Kinase domain | CHLRE_16g669800v5 | XP_001699230.2 | PKA catalytic subunit | P05132 NP_032880.1 |
| PKA catalytic subunit-2 | TTHERM_00658860 | I7MM26 / XP_001024087.2 | Cr 2.24894e-89 | 321 | 37 | 9 | Kinase domain | | | PKA catalytic subunit | P22735 NP._058675.1 |
| PKA regulatory subunit | TTHERM_00623090 | Q240X5 / XP_001022534.2 | Mm 4.05793e-63 | 414 | 48 | 5 | cNMP | CHLRE_03g199050v5 | A0A2K3DZH4 XP_042926603.1 | PKA regulatory subunit | Q9DBC7 NP_001300902.1 |

**Table S4.** Summary of co-immunoprecipitation experiments showing number of total and unique peptides

| Protein | TGD accession numbers | RSP-3HA (bait) | | | | |
|---|---|---|---|---|---|---|
| | | WT | Rsp3A | Rsp4A | Rsp4B | Rsp4C |
| Rsp1 | TTHERM_000196029 | 0 | 1/1 | 0 | 0 | 0 |
| Rsp2 | TTHERM_00394410 | 3/2 | 15/13 | 10/9 | 9/8 | 1/1 |
| Rsp3A | TTHERM_01044600 | 0 | 10/9 | 6/6 | 5/5 | 1/1 |
| Rsp3B | TTHERM_00566810 | 0 | 7/7 | 6/6 | 2/2 | 0 |
| Rsp3C | TTHERM_00418270 | 0 | 11/8 | 6/5 | 2/2 | 1/1 |
| Rsp4A | TTHERM_00427590 | 0 | 21/13 | 22/17 | 6/6 | 4/4 |
| Rsp4B | TTHERM_00502580 | 0 | 4/4 | 5/5 | 2/2 | 2/2 |
| Rsp4C | TTHERM_00444180 | 0 | 18/11 | 8/5 | 2/2 | 1/1 |
| Rsp7A | TTHERM_00670610 | 0 | 8/5 | 2/2 | 2/2 | 1/1 |
| Rsp7B | TTHERM_00492840 | 0 | 4/3 | 4/3 | 1/1 | 0 |
| Rsp8 | TTHERM_00313520 | 0 | 10/8 | 6/6 | 3/3 | 1/1 |
| Rsp9 | TTHERM_00430020 | 1/1 | 31/11 | 23/11 | 9/7 | 0 |
| Rsp10 | TTHERM_00378600 | 0 | 1/1 | 5/5 | 2/2 | 0 |
| Rsp11 | TTHERM_00540090 | 0 | 1/1 | 2/2 | 1/1 | 0 |
| Rsp12A | TTHERM_01018330 | 0 | 1/1 | 0 | 0 | 0 |
| Rsp12B | TTHERM_00466090 | 0 | 0 | 0 | 0 | 0 |
| Rsp14 | TTHERM_000773538 | 0 | 9/5 | 4/3 | 0 | 0 |
| Rsp15 | TTHERM_00695730 | 0 | 4/4 | 0 | 0 | 0 |
| Rsp16A | TTHERM_00471260 | 0 | 4/4 | 4/3 | 2/2 | 3/2 |
| Rsp16B | TTHERM_00238810 | 0 | 0 | 4/4 | 0 | 0 |
| Rsp20/CaM1 | TTHERM_00630500 | 0 | 0 | 0 | 0 | 0 |
| Rsp22/LC8 | TTHERM_000649439 | 0 | 11/5 | 6/4 | 0 | 0 |
| Rsp23/Cfap67A | TTHERM_000372529 | 0 | 14/10 | 4/4 | 4/4 | 3/3 |
| Cfap61 | TTHERM_00641200 | 0 | 19/15 | 15/13 | 2/2 | 4/4 |
| Cfap251 | TTHERM_01262850 | 0 | 7/7 | 11/10 | 9/8 | 4/3 |
| Cfap91 | TTHERM_00578560 | 0 | 2/2 | 7/6 | 5/5 | 1/1 |
| Cfap206 | TTHERM_00820660 | 0 | 2/2 | 5/5 | 0 | 0 |
| Cfap253 | TTHERM_00316930 | 0 | 5/4 | 7/7 | 3/3 | 1/1 |
| Cfap207 | TTHERM_00529880 | 0 | 2/1 | 4/4 | 3/3 | 0 |
| Cfap198A | TTHERM_01092470 | 0 | 1/1 | 0 | 0 | 0 |
| Cfap198B | TTHERM_00476710 | 0 | 0 | 0 | 0 | 0 |
| TtTpr | TTHERM_00623040 | 0 | 1/1 | 0 | 0 | 0 |
| Kelch-motif protein | TTHERM_00760390 | 0 | 4/4 | 6/6 | 3/3 | 1/1 |
| Lrrc23A | TTHERM_000703669 | 0 | 4/3 | 3/3 | 0 | 0 |
| Lrrc23B | TTHERM_00105260 | 0 | 4/4 | 0 | 0 | 0 |
| Lrr-containing | TTHERM_01084360 | 0 | 0 | 0 | 0 | 0 |
| Stpg1 | TTHERM_00549640 | 0 | 0 | 0 | 0 | 0 |
| Stpg2 | TTHERM_00420710 | 0 | 0 | 0 | 0 | 0 |
| adenylate kinase 1, Ak1 | TTHERM_00317200 | 0 | 76/43 | 35/28 | 18/18 | 17/16 |
| adenylate kinase 7A, Ak7A | TTHERM_00558060 | 0 | 7/7 | 8/6 | 4/3 | 0 |
| adenylate kinase 7B, Ak7B | TTHERM_000569069 | 0 | 7/7 | 9/9 | 5/5 | 2/2 |
| adenylate kinase 8A, Ak8A | TTHERM_00227800 | 0 | 12/10 | 9/8 | 7/7 | 5/5 |
| adenylate kinase 8B, Ak8B | TTHERM_00455540 | 0 | 11/8 | 5/5 | 4/4 | 1/1 |
| adenylate kinase 9, Ak9 | TTHERM_00148750 | 0 | 47/37 | 16/16 | 4/4 | 9/9 |
| caseine kinase 1, Ck1, | TTHERM_00938880 | 0 | 6/5 | 2/2 | 3/3 | 1/1 |
| PKA catalytic subunit | TTHERM_00433420 | 0 | 4/3 | 0 | 0 | 2/2 |
| PKA catalytic subunit | TTHERM_00658860 | 0 | 0 | 0 | 0 | 0 |
| PKA regulatory protein | TTHERM_00623090 | 0 | 1/1 | 3/3 | 2/2 | 0 |

**Table S5.** Summary of BioID experiments showing number of total and unique peptides

| Protein | TGD accession numbers | Rsp 3A | Rsp 3A | Rsp 3A | Rsp 3B | Rsp 3B | Rsp 3B | Rsp 3B | Rsp 3B | Rsp 3C | Rsp 4A | Rsp 4A | Rsp 4A | Rsp 4A | Rsp 4B | Rsp 4C | Rsp 4C | Cfap 206 | Cfap 206 | Cfap 206 | Cfap 61 | Cfap 61 | Cfap 61 | Cfap 91 | Lrrc 23A | Lrrc 23B | Lrrc 23B |
|---|---|---|---|---|---|---|---|---|---|---|---|---|---|---|---|---|---|---|---|---|---|---|---|---|---|---|---|
| Rsp1 | TTHERM_000196029 | 2/2 | 2/2 | 1/1 | 1/1 | 5/4 | 0 | 0 | 5/4 | 0 | 0 | 16/9 | 2/2 | 16/9 | 2/2 | 0 | 2/2 | 0 | 0 | 0 | 0 | 0 | 0 | 0 | 7/3 | | 8/7 |
| Rsp2 | TTHERM_00394410 | 21/17 | 25/15 | 28/15 | 11/9 | 72/32 | 6/5 | 12/6 | 86/33 | 0 | 0 | 63/38 | 2/2 | 55/32 | 0 | 10/5 | 2/2 | 8/8 | 0 | 0 | 0 | 0 | 2/2 | 1/1 | 38/21 | | 44/20 |
| Rsp3A | TTHERM_01044600 | 16/12 | 16/9 | 21/11 | 6/6 | 46/26 | 1/1 | 2/1 | 52/28 | 0 | 1/1 | 27/18 | 0 | 25/17 | 0 | 3/2 | 1/1 | 0 | 0 | 0 | 0 | 0 | 0 | 0 | | | 5/5 |
| Rsp3B | TTHERM_00566810 | 14/11 | 2/2 | 5/4 | 12/9 | 42/20 | 1/1 | 1/1 | 49/23 | 9/7 | 0 | 32/20 | 1/1 | 25/16 | 1/1 | 6/5 | 1/1 | 6/6 | 7/5 | 7/5 | 4/4 | 0 | 0 | 1/1 | 32/20 | | 45/18 |
| Rsp3C | TTHERM_00418270 | 0 | 0 | 0 | 3/3 | 4/3 | 0 | 0 | 6/5 | 10/7 | 0 | 25/22 | 0 | 23/20 | 0 | 1/1 | 0 | 11/11 | 4/4 | 6/5 | 5/5 | 7/6 | 11/8 | 11/9 | 60/27 | | 77/32 |
| Rsp4A | TTHERM_00427590 | 3/3 | 14/8 | 12/7 | 0 | 36/18 | 1/1 | 3/2 | 39/19 | 0 | 5/4 | 76/27 | 15/13 | 67/25 | 9/9 | 13/10 | 20/16 | 1/1 | 0 | 0 | 0 | 0 | 0 | 0 | 15/13 | | 26/14 |
| Rsp4B | TTHERM_00502580 | 0 | 0 | 0 | 0 | 1/1 | 0 | 0 | 1/1 | 0 | 0 | 26/17 | 1/1 | 22/14 | 8/7 | 0 | 0 | 0 | 0 | 0 | 0 | 0 | 0 | 1/1 | 14/8 | | 11/7 |
| Rsp4C | TTHERM_00444180 | 6/6 | 6/5 | 7/5 | 4/4 | 36/18 | 0 | 0 | 40/20 | 0 | 2/2 | 51/23 | 3/3 | 43/20 | 0 | 11/9 | 24/16 | 0 | 0 | 0 | 0 | 0 | 0 | 0 | | | 2/2 |
| Rsp7A | TTHERM_00670610 | 5/2 | 0 | 1/1 | 4/1 | 12/5 | 0 | 0 | 13/5 | 11/3 | 0 | 16/4 | 0 | 14/3 | 0 | 3/2 | 1/1 | 4/2 | 0 | 0 | 0 | 0 | 0 | 0 | 27/4 | | 20/3 |
| Rsp7B | TTHERM_00492840 | 0 | 0 | 0 | 2/2 | 0 | 0 | 0 | 0 | 1/1 | 0 | 1/1 | 0 | 1/1 | 0 | 0 | 0 | 2/2 | 1/1 | 1/1 | 5/5 | 0 | 3/2 | 0 | 82/32 | | 88/29 |
| Rsp8 | TTHERM_00313520 | 0 | 0 | 0 | 3/3 | 1/1 | 0 | 0 | 1/1 | 5/4 | 2/2 | 18/3 | 0 | 14/11 | 0 | 0 | 0 | 9/9 | 4/2 | 6/4 | 1/1 | 4/3 | 5/4 | 5/3 | 37/21 | | 61/24 |
| Rsp9 | TTHERM_00430020 | 10/9 | 16/6 | 20/8 | 2/2 | 52/16 | 2/2 | 5/4 | 54/18 | 0 | 2/2 | 79/20 | 11/9 | 74/20 | 10/9 | 16/9 | 22/15 | 6/6 | 1/1 | 2/2 | 0 | 0 | 3/2 | 1/1 | 15/7 | 2/1 | 19/7 |
| Rsp10 | TTHERM_00378600 | 3/3 | 2/2 | 2/2 | 0 | 18/10 | 0 | 0 | 21/11 | 0 | 1/1 | 22/14 | 0 | 19/11 | 0 | 2/2 | 5/5 | 0 | 0 | 0 | 0 | 0 | 0 | 0 | 6/5 | | 18/11 |
| Rsp11 | TTHERM_00540090 | 1/1 | 0 | 2/1 | 0 | 7/2 | 0 | 0 | 8/3 | 0 | 0 | 3/2 | 0 | 3/2 | 0 | 0 | 0 | 2/2 | 0 | 1/1 | 0 | 0 | 0 | 2/1 | 8/3 | | 8/3 |
| Rsp12A | TTHERM_01018330 | 0 | 0 | 0 | 0 | 0 | 0 | 0 | 0 | 0 | 0 | 0 | 0 | 0 | 4/4 | 0 | 0 | 0 | 0 | 0 | 0 | 0 | 0 | 0 | | | |
| Rsp12B | TTHERM_00466090 | 0 | 0 | 0 | 0 | 0 | 0 | 0 | 0 | 0 | 1/1 | 0 | 0 | 0 | 0 | 0 | 0 | 0 | 0 | 0 | 0 | 0 | 0 | 0 | | | |
| Rsp14 | TTHERM_000773538 | 13/12 | 4/2 | 5/3 | 3/3 | 40/18 | 1/1 | 1/1 | 42/19 | 0 | 1/1 | 27/20 | 0 | 23/17 | 0 | 4/3 | 0 | 0 | 0 | 0 | 0 | 0 | 0 | 0 | | | 8/4 |
| Rsp15 | TTHERM_00695730 | 0 | 0 | 0 | 2/2 | 0 | 0 | 0 | 0 | 1/1 | 1/1 | 10/8 | 0 | 8/8 | 0 | 0 | 0 | 3/3 | 3/2 | 3/2 | 0 | 0 | 0 | 2/2 | 32/11 | | 31/10 |
| Rsp16A | TTHERM_00471260 | 6/5 | 6/4 | 8/5 | 3/2 | 19/11 | 1/1 | 1/1 | 21/12 | 0 | 0 | 16/11 | 0 | 14/10 | 0 | 1/1 | 0 | 2/2 | 0 | 0 | 0 | 0 | 0 | 0 | 18/11 | | 22/10 |
| Rsp16B | TTHERM_00238810 | 0 | 0 | 0 | 0 | 1/1 | 0 | 1/1 | 2/2 | 1/1 | 0 | 8/6 | 0 | 8/6 | 0 | 0 | 0 | 2/2 | 3/2 | 3/2 | 0 | 0 | 0 | 4/3 | 26/12 | | 31/13 |
| Rsp20/CaM1 | TTHERM_00630500 | 0 | 0 | 0 | 0 | 9/5 | 0 | 1/1 | 11/6 | 0 | 0 | 3/3 | 0 | 2/2 | 0 | 0 | 0 | 1/1 | 0 | 0 | 0 | 0 | 0 | 0 | | | |
| Rsp22 | TTHERM_000649439 | 1/1 | 5/2 | 6/2 | 1/1 | 26/3 | 0 | 0 | 28/4 | 3/1 | 2/2 | 20/5 | 0 | 19/15 | 1/1 | 2/2 | 1/1 | 2/2 | 4/2 | 4/2 | 0 | 0 | 0 | 7/2 | 26/3 | 1/1 | 33/3 |
| Rsp23/Cfap67A | TTHERM_000372529 | 0 | 0 | 0 | 0 | 2/2 | 0 | 0 | 3/3 | 0 | 0 | 9/9 | 0 | 8/8 | 0 | 0 | 0 | 0 | 0 | 0 | 0 | 0 | 0 | 0 | | | |
| Cfap61 | TTHERM_00641200 | 0 | 0 | 0 | 6/6 | 2/2 | 0 | 1/1 | 3/3 | 10/9 | 1/1 | 7/7 | 1/1 | 6/6 | 1/1 | 1/1 | 0 | 12/11 | 6/5 | 6/5 | 13/10 | 16/10 | 21/14 | 18/10 | 146/59 | | 160/60 |
| Cfap251 | TTHERM_01262850 | 0 | 0 | 0 | 2/2 | 1/1 | 0 | 0 | 1/1 | 4/2 | 0 | 17/15 | 0 | 16/14 | 0 | 0 | 0 | 8/8 | 2/2 | 2/2 | 6/6 | 10/6 | 13/9 | 26/14 | 96/42 | | 123/39 |
| Cfap91 | TTHERM_00578560 | 0 | 0 | 0 | 1/1 | 0 | 1/1 | 1/1 | 0 | 8/4 | 0 | 21/15 | 0 | 18/12 | 0 | 1/1 | 0 | 6/6 | 2/2 | 2/2 | 3/3 | 3/1 | 4/2 | 9/6 | 74/29 | | 84/28 |
| Cfap206 | TTHERM_00820660 | 0 | 0 | 0 | 3/3 | 0 | 0 | 0 | 1/1 | 7/4 | 1/1 | 15/10 | 0 | 10/7 | 1/1 | 0 | 0 | 12/12 | 9/6 | 10/6 | 8/8 | 2/1 | 3/2 | 5/2 | 60/29 | | 68/26 |
| Cfap253 | TTHERM_00316930 | 5/5 | 0 | 2/2 | 2/2 | 35/17 | 0 | 0 | 41/20 | 0 | 1/1 | 36/18 | 0 | 28/15 | 0 | 0 | 0 | 0 | 0 | 0 | 0 | 0 | 0 | 0 | | | 7/6 |
| Cfap207 | TTHERM_00529880 | 0 | 0 | 1/1 | 0 | 0 | 0 | 0 | 0 | 0 | 0 | 9/8 | 0 | 7/7 | 0 | 0 | 0 | 0 | 0 | 0 | 0 | 0 | 0 | 0 | 2/2 | | 3/3 |
| Cfap198A | TTHERM_01092470 | 2/2 | 1/1 | 1/1 | 1/1 | 11/5 | 0 | 0 | 11/5 | 0 | 0 | 10/9 | 0 | 8/5 | 0 | 0 | 0 | 0 | 0 | 0 | 0 | 0 | 0 | 0 | | | |
| Cfap198B | TTHERM_00476710 | 0 | 0 | 0 | 1/1 | 0 | 0 | 0 | 0 | 0 | 0 | 6/4 | 0 | 5/4 | 0 | 0 | 0 | 2/2 | 0 | 0 | 0 | 0 | 0 | 0 | 17/6 | | 19/6 |
| TtTpr | TTHERM_00623040 | 0 | 0 | 0 | 2/2 | 0 | 0 | 0 | 1/1 | 6/3 | 0 | 6/6 | 0 | 5/5 | 1/1 | 0 | 0 | 7/6 | 7/5 | 9/6 | 0 | 1/1 | 2/2 | 6/3 | 19/8 | | 24/12 |
| Kelch-motif | TTHERM_00760390 | 0 | 0 | 0 | 2/2 | 2/2 | 1/1 | 1/1 | 3/3 | 1/1 | 0 | 7/6 | 0 | 7/6 | 0 | 1/1 | 0 | 1/1 | 4/3 | 4/3 | 1/1 | 0 | 0 | 3/3 | 27/12 | | 20/11 |
| Lrrc23A | TTHERM_000703669 | 0 | 0 | 0 | 0 | 0 | 0 | 0 | 1/1 | 0 | 0 | 1/1 | 0 | 1/1 | 0 | 0 | 0 | 0 | 0 | 0 | 0 | 0 | 0 | 0 | 26/12 | 3/1 | 2/2 |
| Lrrc23B | TTHERM_00105260 | 0 | 0 | 0 | 0 | 1/1 | 0 | 0 | 1/1 | 0 | 0 | 1/1 | 0 | 1/1 | 0 | 0 | 0 | 0 | 0 | 0 | 0 | 0 | 0 | 0 | | | 49/13 |
| Lrrc | TTHERM_01084360 | 0 | 0 | 0 | 0 | 0 | 0 | 0 | 0 | 0 | 0 | 0 | 0 | 0 | 0 | 0 | 0 | 0 | 0 | 0 | 0 | 0 | 0 | 0 | | | |
| Stpg1 | TTHERM_00549640 | 0 | 0 | 0 | 0 | 0 | 0 | 0 | 0 | 0 | 0 | 0 | 0 | 0 | 0 | 0 | 0 | 0 | 0 | 0 | 0 | 0 | 0 | 0 | | | |
| Stpg2 | TTHERM_00420710 | 0 | 0 | 0 | 0 | 0 | 0 | 0 | 0 | 0 | 0 | 0 | 0 | 0 | 0 | 0 | 0 | 0 | 0 | 0 | 0 | 0 | 0 | 0 | | | |
| Ak1 | TTHERM_00317200 | 17/15 | 21/9 | 25/11 | 7/7 | 12/12 | 2/2 | 2/2 | 16/15 | 2/2 | 1/1 | 37/29 | 1/1 | 29/23 | 1/1 | 17/12 | 3/3 | 8/8 | 1/1 | 1/1 | 0 | 0 | 0 | 0 | 47/27 | 1/1 | 54/31 |
| Ak7A | TTHERM_00558060 | 0 | 0 | 0 | 0 | 0 | 0 | 0 | 0 | 0 | 0 | 2/2 | 0 | 2/2 | 0 | 0 | 0 | 0 | 0 | 0 | 0 | 0 | 0 | 0 | 11/9 | | 13/8 |
| Ak7B | TTHERM_000569069 | 0 | 0 | 0 | 0 | 1/1 | 0 | 0 | 1/1 | 0 | 0 | 3/3 | 0 | 3/3 | 0 | 0 | 0 | 0 | 0 | 0 | 0 | 0 | 0 | 0 | 28/17 | 25/13 | 9/7 |
| Ak8A | TTHERM_00227800 | 5/5 | 16/10 | 16/10 | 7/7 | 36/19 | 7/4 | 8/4 | 41/21 | 7/4 | 3/3 | 37/20 | 2/2 | 30/18 | 3/3 | 7/5 | 7/7 | 16/11 | 13/8 | 14/9 | 1/1 | 6/4 | 10/6 | 23/14 | 67/25 | | 69/23 |
| Ak8B | TTHERM_00455540 | 8/7 | 11/7 | 13/8 | 4/4 | 31/19 | 1/1 | 1/1 | 36/20 | 0 | 1/1 | 24/18 | 1/1 | 22/17 | 0 | 0 | 1/1 | 0 | 0 | 0 | 0 | 0 | 0 | 0 | 1/1 | | 16/9 |
| Ak9 | TTHERM_00148750 | 0 | 0 | 0 | 0 | 5/5 | 0 | 0 | 7/7 | 1/1 | 0 | 18/18 | 0 | 16/16 | 0 | 1/1 | 0 | 1/1 | 0 | 0 | 0 | 0 | 0 | 0 | 156/63 | | 13/10 |
| Ck1 | TTHERM_00938880 | 0 | 0 | 0 | 0 | 0 | 0 | 0 | 0 | 0 | 0 | 3/3 | 0 | 2/2 | 0 | 0 | 0 | 0 | 0 | 0 | 0 | 0 | 0 | 0 | | | 5/4 |
| PKA catalytic sub | TTHERM_00433420 | 0 | 0 | 0 | 0 | 0 | 0 | 0 | 0 | 0 | 0 | 0 | 0 | 0 | 0 | 0 | 0 | 0 | 0 | 0 | 0 | 0 | 0 | 0 | | | |
| PKA catalytic sub | TTHERM_00658860 | 0 | 0 | 0 | 0 | 0 | 0 | 0 | 0 | 0 | 0 | 0 | 0 | 0 | 0 | 0 | 0 | 0 | 0 | 0 | 0 | 0 | 0 | 0 | | | |
| PKA regulatory sub | TTHERM_00623090 | 0 | 0 | 0 | 0 | 0 | 0 | 0 | 0 | 0 | 0 | 0 | 0 | 0 | 0 | 0 | 0 | 0 | 0 | 0 | 0 | 0 | 0 | 0 | | | |

**Table S6.** *Tetrahymena* RSP orthologs identified with multiple hits in TGD. When numerous proteins were identified with similar scores in blastp search of *Tetrahymena* total proteome (TGD) using *Chlamydomonas* RSP as a bait, we analyzed which of those proteins are present in *Tetrahymena* ciliome (LFQ and TMT data) and either they co-immumoprecipitate (co-IP, Table S5) with known RSPs or are biotinylated (BioID, Table S6) in cells expressing known RS proteins as fusions with mutated BirA* ligase. Arrows indicate a significant change (↑ increase or ↓ decrease) in the protein level in studied knock-out mutants (based on TMT analyses). ns – change not significant, nd – protein not detected in the *Tetrahymena* ciliome.

| | | E-value (TGD blastp) | In TMT ciliome | In co-IP or BioID | RSP3A-KO | RSP3B-KO | RSP3C-KO | FAP91-KO |
|---|---|---|---|---|---|---|---|---|
| **Rsp1** | **TTHERM_000196029**<br>new number<br>**TTHERM_00196025** | 1.78444e-19 | yes | coIP 3A<br>Bir 3A, 3B, 4A, 4C | ns | ↓ | ns | ns |
| **Rsp10** | **TTHERM_00378600** | 1.90468e-19 | yes | coIP 4A<br>Bir 3A, 3B, 4A, 4C | ns | ↓ | ns | ns |
| | TTHERM_00338260 | 7.69708e-19 | no | - | - | - | - | - |
| | TTHERM_00561720 | 4.08366e-17 | yes | nd | ns | ns | ns | ns |
| | TTHERM_00086700 | 3.34742e-16 | yes | Bir 3B (1), 4A (1) | ns | ↓ | ↓ | ↓ |
| | TTHERM_00940320 | 4.65599e-16 | no | - | - | - | - | - |
| | TTHERM_00471440 | 3.45586e-15 | no | - | - | - | - | - |
| | TTHERM_00522150 | 6.02902e-15 | no | - | - | - | - | - |
| | TTHERM_00013150 | 6.59314e-15 | no | - | - | - | - | - |
| | TTHERM_00122390 | 1.03558e-14 | no | - | - | - | - | - |
| | TTHERM_01044720 | 4.35786e-14 | no | - | - | - | - | - |
| | TTHERM_00058629 | 8.15103e-14 | yes | nd | ns | ↓ | ns | ns |
| | TTHERM_00047490 | 8.87354e-14 | yes | nd | ↑ | ns | ns | ↓ |
| | TTHERM_00080020 | 1.83831e-13 | yes | nd | ns | ns | ns | ns |
| | TTHERM_00138540 | 1.85468e-13 | no | - | - | - | - | - |
| | TTHERM_00133460 | 4.62151e-13 | no | - | - | - | - | - |
| | TTHERM_00149910 | 5.33197e-13 | no | - | - | - | - | - |
| | TTHERM_00691730 | 6.19233e-13 | no | - | - | - | - | - |
| | TTHERM_00142460 | 6.24492e-13 | no | - | - | - | - | - |
| | TTHERM_01248870 | 6.40885e-13 | no | - | - | - | - | - |
| | TTHERM_00724730 | 1.06173e-12 | no | - | - | - | - | - |
| | TTHERM_00078940 | 1.13884e-12 | yes | nd | ns | ↓ | ns | ns |
| | TTHERM_00685930 | 1.18943e-12 | yes | nd | ns | ↓ | ↓ | ↓ |
| | TTHERM_00509110 | 2.45575e-12 | yes | nd | ns | ns | ↑ | ns |
| | TTHERM_00486410 | 4.33653e-12 | yes | nd | ns | ns | ns | ns |
| | TTHERM_00590130 | 5.60043e-12 | yes | nd | ns | ↓ | ↓ | ↓ |
| | TTHERM_00077350 | 6.44267e-12 | yes | nd | ns | ns | ns | ns |
| | TTHERM_00329860 | 6.74109e-12 | no | - | - | - | - | - |
| | TTHERM_01084140 | 8.55508e-12 | no | - | - | - | - | - |
| | TTHERM_00637100 | 8.74526e-12 | no | - | - | - | - | - |
| | TTHERM_00196080 | 1.13797e-11 | yes | nd | ↓ | ↓ | ↓ | ↓ |
| | TTHERM_00766490 | 1.31424e-10 | yes | nd | ↓ | ↓ | ↓ | ↓ |
| | TTHERM_00584930 | 1.95024e-10 | yes | nd | ↑ | ↑ | ns | ↓ |
| | TTHERM_00732890 | 3.04534e-10 | yes | nd | ns | ns | ns | ns |
| **Rsp5** | TTHERM_00962090 | none | yes | nd | ↑ | ns | ↑ | ↑ |
| | TTHERM_00338200 | none | yes | nd | ↑ | ns | ns | ↓ |
| | TTHERM_01080330 | none | yes | nd | ↑ | ↑ | ns | ns |
| | TTHERM_00058260 | none | yes | nd | ↑ | ↑ | ns | ns |
| | TTHERM_00526310 | none | yes | nd | ↑ | ↑ | ↑ | ns |
| | TTHERM_00300640 | none | yes | nd | ↑ | ns | ns | ns |
| | TTHERM_00300630 | none | yes | nd | ns | ↓ | ↓ | ns |
| | TTHERM_00526449 | none | yes | nd | ns | ns | ↑ | ↓ |
| | TTHERM_00274520 | none | yes | nd | ns | ↓ | ↓ | ↓ |
| | TTHERM_00266330 | none | yes | nd | ns | ↓ | ns | ns |
| | TTHERM_00300620 | none | yes | nd | ns | ns | ns | ↑ |
| | TTHERM_00703880 | none | yes | nd | ns | ↑ | ↑ | ns |
| | TTHERM_01002600 | none | yes | nd | ns | ns | ↓ | ↓ |
| **Rsp7** | TTHERM_00353490 | 7.66049e-13 | yes | nd | ns | ↑ | ns | ↑ |
| | TTHERM_00686010 | 2.22411e-09 | no | - | - | - | - | - |
| | TTHERM_00384910 | 5.51363e-09 | yes | nd | ↓ | ns | ns | ↑ |
| | TTHERM_00632950 | 1.57971e-08 | no | - | - | - | - | - |
| | TTHERM_00486220 | 8.14257e-08 | no | - | - | - | - | ↑ |
| | TTHERM_00523059 | 1.054e-07 | yes | nd | ↓ | ns | ↓ | - |

| | | | | | | | | |
|---|---|---|---|---|---|---|---|---|
| | TTHERM_00059190 | 1.12651e-07 | yes | nd | ↓ | ↓ | ↓ | ↓ |
| | TTHERM_00668030 | 2.30983e-07 | yes | nd | ns | ns | ns | ns |
| | TTHERM_000442899 (Stk36-like) | 4.65608e-07 | yes | co-IP RSP4C (1) Bir RSP4A (1-2), 3B (1) 4B (3/3) | ns | ↑ | ns | ↑ |
| | TTHERM_00300120 | 6.8876e-07 | yes | nd | ns | ↑ | ns | ns |
| | TTHERM_00630500 (CaM1) | 9.98653e-07 | yes | Bir 4A (2-3), 3B (1-11) F206 (1) | ns | ↓ | ↓ | ns |
| | TTHERM_00622900 | 2.83945e-06 | no | - | - | - | - | - |
| | TTHERM_00267880 (Bbc37) | 4.07889e-06 | yes | coIP 3A(1), 4A(4) | ns | ↓ | ↓ | ↑ |
| | TTHERM_00194410 | 5.09536e-06 | no | - | - | - | - | - |
| | TTHERM_00616570 | 5.2479e-06 | yes | nd | ↓ | ns | ns | ns |
| | TTHERM_00218310 | 9.61136e-06 | yes | nd | ns | ↓ | ↓ | ↓ |
| | TTHERM_00589960 | 1.36054e-05 | yes | nd | ns | ns | ns | ns |
| | TTHERM_00899430 | 1.57e-05 | no | - | - | - | - | - |
| | TTHERM_00495950 | 1.90483e-05 | yes | nd | ↑ | ns | ns | ↑ |
| | TTHERM_01052880 | 2.2429e-05 | yes | nd | ns | ↓ | ns | ↑ |
| **Rsp7A** | **TTHERM_00670610** | 3.31578e-05 | yes | co-IP 4A(2), 4B(2), 4C(1), 3A(8) Bir 3A(1-5), 3B(4-13), 3C(11), 4A(14-16), 4C(1-3), 206 (4) | ns | ↓ | ns | ↓ |
| | TTHERM_00637390 | 4.33175e-05 | yes | nd | ↑ | ↓ | ns | ↓ |
| | TTHERM_000841299 | 5.08855e-05 | no | - | - | - | - | - |
| | TTHERM_00147620 | 6.24849e-05 | yes | nd | ns | ns | ns | ns |
| **Rsp7B** | **TTHERM_00492840** | 6.65092e-05 | yes | coIP 3A (4), 4A(4), 4B (1) Bir 3B(2), 3C(1), 4A(1), 206(1), 61(3-5) | ↑ | ↑ | ↑ | ↓ |
| | TTHERM_00136380 | 0.00013603 | no | - | - | - | - | - |
| | TTHERM_00079490 | 0.00017402 | yes | nd | ns | ↑ | ns | ns |
| | TTHERM_00013440 | 0.00017893 | no | - | - | - | - | - |
| | TTHERM_00371010 | 0.00021959 | yes | nd | ns | ↓ | ns | ns |
| **Rsp12** | TTHERM_00051740 | 1.0883e-25 | yes | Bir 3 A(1 pep) | ns | ↑ | ns | ns |
| | TTHERM_00997720 | 1.64934e-25 | no | - | - | - | - | - |
| | TTHERM_00890120 | 2.17043e-25 | yes | nd | ↑ | ns | ↑ | ↑ |
| | TTHERM_00243800 | 1.07131e-22 | no | - | - | - | - | - |
| | TTHERM_00118710 | 5.99925e-22 | yes | nd | ns | ↓ | ↓ | ns |
| | TTHERM_00492720 | 6.01654e-22 | no | - | - | - | - | - |
| | TTHERM_00548130 | 4.03513e-21 | yes | nd | ns | ↓ | ↓ | ↑ |
| | **TTHERM_01018330** | 2.06701e-19 | yes | coIP 3a (1) BioID4B (4) | ns | ↓ | ↓ | ↓ |
| | TTHERM_00865350 | 5.9945e-19 | yes | nd | ↑ | ↓ | ns | ns |
| | TTHERM_00877050 | 1.9254e-18 | no | - | - | - | - | - |
| | **TTHERM_00466090** | 3.9832e-18 | yes | Bir 4a (1) | ↓ | ↓ | ns | ns |
| | TTHERM_00075590 | 7.37871e-18 | no | - | - | - | - | - |
| | TTHERM_00772040 | 1.51296e-17 | no | - | - | - | - | - |
| | TTHERM_00538480 | 7.1862e-15 | no | - | - | - | - | - |

**RSP1** and **RSP10** – proteins contain multiple MORN domains. Using *Chlamydomonas* RSP1 as a bait, *Tetrahymena* Rsp1 and Rsp10 were identified as a 9$^{th}$ and 36$^{th}$ hits, using *Chlamydomonas* RSP10 as a bait, *Tetrahymena* Rsp1 and Rsp10 were identified as a 1$^{st}$ and 2$^{nd}$ hits (shown in this Table).

**Table S7.** Bases of the identification of RS subunits. Cross-linking data [12]

| protein | TGD accession numbers | radial spoke | Mutant ciliomes |
|---|---|---|---|
| Rsp1 | TTHERM_000196029 | RS1<br>RS2 | Comparative global ciliome analyses:<br>• The level of Rsp1 is diminished in *RSP3B* knockout<br>Co-IP and BioID data:<br>• Rsp1 is effectively biotinylated in cells expressing BirA*-tagged protein: Rsp3A, Rsp3B, and Rsp4A<br>Cross-link data:<br>• Rsp1 cross-links with Rsp4A and Rsp2 |
| Rsp2 | TTHERM_00394410 | RS1<br>RS2 | Comparative global ciliome analyses:<br>• The level of Rsp2 is diminished in *RSP3B* knockouts<br>• The level of Rsp2 is unchanged in CFAP61-KO and CFAP91-KO<br>Co-IP and BioID data:<br>• Rsp2 co-precipitates with Rsp3A, Rsp4A, and Rsp4B<br>• Rsp2 is identified among proteins biotinylated in cells expressing BirA*-tagged protein: Rsp3A, Rsp3B, Rsp4A, Rsp4C<br>• Rsp2 is poorly or not biotinylated in cells expressing RS3 protein, Cfap61-HA-BirA*<br>Cross-link data:<br>• Rsp2 cross-links with Rsp1, Rsp4A, and Rsp4C |
| Rsp3A | TTHERM_01044600 | RS1 | Cryo-ET analyses:<br>• a subpopulation of RS1 spokes is completely or partly missing in RSP3A-KO cilia<br>• there are no defects in RS2 and RS3 spokes in RSP3A-KO cilia<br>Cross-link data:<br>• Rsp3A cross-links with Rsp3B and Rsp14 |
| Rsp3B | TTHERM_00566810 | RS1<br>RS2 | Cryo-ET analyses:<br>• a subpopulation of RS1 spokes is missing in RSP3B-KO cilia<br>• all RS2 spokes are eliminated in RSP3B-KO cilia<br>• there are no defects in RS3 spokes<br>Cross-link data:<br>• Rsp3B cross-links with Rsp3A, RSP9, Rsp22, and Cfap253 |
| Rsp3C | TTHERM_00418270 | RS2 | Cryo-ET analyses:<br>• a subpopulation of RS2 spokes is completely or partly missing in RSP3C-KO cilia<br>• there are no defects in RS1 and RS3 spokes |
| Rsp4A | TTHERM_00427590 | RS1<br>RS2 | Comparative global ciliome analyses:<br>• the level of Rsp4A is reduced in cells with knocked out *RSP3B* gene<br>Co-IP and BioID data:<br>• Rsp4A co-precipitates with Rsp2, Rsp3 paralogs, Rsp4A, Rsp4C, Rsp9, and Rsp10<br>• Rsp4A is biotinylated in cells expressing BirA*-tagged Rsp3A, Rsp3B, Rsp4B, Rsp4C<br>• Rsp4A-HA-BirA* biotinylates Rsp2, Rsp3 paralogs, Rsp4 paralogs, Rsp7A, Rsp8, Rsp9, Rsp10, Rsp14, Rsp15<br>Cross-link data:<br>• Rsp4A cross-links with Rsp1, Rsp2, Rsp4B, and Rsp4C |
| Rsp4B | TTHERM_00502580 | Rsp3C-containing RS2 subtype | Comparative global ciliome analyses:<br>• the level of Rsp4B is reduced in cells with knocked out *RSP3C* gene<br>Co-IP and BioID data:<br>• Rsp4B co-precipitates with Rsp2, Rsp3 paralogs, Rsp4A, Rsp4C, Rsp9, and Rsp10<br>• Rsp4B is biotinylated in Rsp4A-HA-BirA* cells<br>• Rsp4B-HA-BirA* biotinylates Rsp2, Rsp4A, Rsp9, Rsp12A |
| Rsp4C | TTHERM_00444180 | RS1<br>RS2 | Comparative global ciliome analyses:<br>• the level of Rsp4C is reduced in cells with knocked out *RSP3B* gene<br>Co-IP and BioID data:<br>• Rsp4C co-precipitates with Rsp3A and Rsp4A.<br>• Rsp4C is biotinylated in cells expressing BirA*-tagged Rsp3A, Rsp3B, and Rsp4A<br>• Rsp4C-HA-BirA* biotinylates Rsp2, Rsp4A, Rsp9, Rsp10, and Rsp14<br>Cross-link data:<br>• Rsp4C cross-links with Rsp2, Rsp4A, and Rsp16A |
| Rsp7A | TTHERM_00670610 | RS1<br>RS2 | Comparative global ciliome analyses:<br>• the level of Rsp7A is reduced in *RSP3B* and *CFAP91* knockouts<br>Co-IP and BioID data:<br>• Rsp7A co-precipitates with Rsp3A, Rsp4A, Rsp4B<br>• Rsp7A is identified among proteins biotinylated in cells expressing BirA*-tagged protein: Rsp3A, Rsp3B, Rsp3C, Rsp4A, Rsp4C<br>• Rsp7A is not biotinylated in cells expressing RS3 protein, Cfap61-HA-BirA* |

| | | | |
|---|---|---|---|
| | | | Cross-link data:<br>• Rsp7A cross-links with Rsp14 and Rsp22 |
| Rsp7B | TTHERM_00492840 | RS3 | Comparative global ciliome analyses:<br>• the level of Rsp7B is reduced only in *CFAP91* knockouts<br>Co-IP and BioID data:<br>• Rsp7B is biotinylated in cells expressing Cfap61-HA-BirA*, Lrrc23A, Lrrc23B |
| Rsp8 | TTHERM_00313520 | RS2 | Comparative global ciliome analyses:<br>• the level of Rsp8 is reduced in *RSP3B*, RSP3C, and *CFAP206* knockouts (mutants with RS2 defects) |
| Rsp9 | TTHERM_00430020 | RS1<br>RS2 | Comparative global ciliome analyses:<br>• the level of Rsp9 is reduced in *RSP3B* knockout<br>Co-IP and BioID data:<br>• Rsp9 is effectively biotinylated in cells expressing BirA*-tagged protein: Rsp3 and Rsp4 paralogs<br>• Rsp9 is not or poorly biotinylated in cells expressing RS3 protein, Cfap61-HA-BirA*<br>• Rsp9 co-precipitates with Rsp3A, Rsp4A, and Rsp4B<br>Cross-link data:<br>• Rsp9 cross-links with Rsp3B |
| Rsp10 | TTHERM_00378600 | RS1<br>RS2 | Comparative global ciliome analyses:<br>• the level of Rsp10 is reduced in *RSP3B* knockout<br>Co-IP and BioID data:<br>• Rsp10 is effectively biotinylated in cells expressing BirA*-tagged protein: Rsp3 and Rsp4 paralogs<br>• Rsp10 is not biotinylated in cells expressing RS3 protein, Cfap61-HA-BirA*<br>• Rsp10 co-precipitates with Rsp4A and Rsp4B |
| Rsp11 | TTHERM_00540090 | RS1<br>RS2 | Comparative global ciliome analyses:<br>• the level of Rsp11 is reduced in all *RSP3* knockout mutants<br>• the level of Rsp11 is unaffected in CFAP91-KO mutant<br>Co-IP and BioID data:<br>• Rsp11 is biotinylated in cells expressing BirA*-tagged protein: Rsp3A, Rsp3B, Rsp4A,<br>• Rsp11 is not biotinylated in cells expressing RS3 protein, Cfap61-HA-BirA* |
| Rsp12A | TTHERM_01018330 | RS2 | Comparative global ciliome analyses:<br>• The level of Rsp12A is reduced in *RSP3B*, *RSP3C*, and *CFAP206* knockouts (mutants with RS2 defects)<br>Co-IP and BioID data:<br>• Rsp12A is biotinylated in cells expressing Rsp4B-HA-BirA* |
| Rsp12B | TTHERM_00466090 | RS1<br>RS2 (?) | Comparative global ciliome analyses:<br>• The level of Rsp12B is reduced in *RSP3A* and *RSP3B* knockouts |
| Rsp14 | TTHERM_000773538 | RS1 | Comparative global ciliome analyses:<br>• The level of Rsp14 is reduced in *RSP3A* and *RSP3B* knockouts<br>• The level of Rsp14 is not significantly altered in *RSP3C* and *CFAP206* mutants<br>Co-IP and BioID data:<br>• Rsp14 co-precipitates with Rsp3A,<br>Cross-link data:<br>• Rsp14 cross-links with Rsp3A, Rsp7A, and Rsp22 |
| Rsp15 | TTHERM_00695730 | | Comparative global ciliome analyses:<br>• The level of Rsp15 is reduced in *RSP3B*, RSP3C, and *CFAP206* knockouts (mutants with RS2 defects), and in CFAP91-KO mutant (likely due to RS2 base defect)<br>Co-IP and BioID data:<br>• Rsp15 is biotinylated in cells expressing Rsp3B, Rsp4A, or Cfap206 tagged with BirA*<br>• Rsp15 is not biotinylated in Rsp3A-HA-BirA* and Cfap61-HA-BirA* cells |
| Rsp16A | TTHERM_00471260 | RS1<br>Rsp3B-containing RS2 subtype | Comparative global ciliome analyses:<br>• The level of Rsp16A is reduced in *RSP3A* and *RSP3B* knockouts<br>Co-IP and BioID data:<br>• Rsp16A is biotinylated in cells expressing BirA* tagged Rsp3A, Rsp3B, Rsp4A<br>Cross-link data:<br>• Rsp16A cross-links with Rsp4C |
| Rsp16B | TTHERM_00238810 | RS2 | Comparative global ciliome analyses:<br>• The level of Rsp16B is reduced in *RSP3C*, *CFAP206* (mutants with RS2 defects), and *CFAP91* knockouts (also defects in RS2 base)<br>Co-IP and BioID data:<br>• Rsp16B is biotinylated in cells expressing Rsp4A-HA-BirA* |
| Rsp20/CaM1 | TTHERM_00630500 | RS2 | Comparative global ciliome analyses:<br>• The level of Rsp20 is reduced in *RSP3B* and *RSP3C* knockouts<br>Co-IP and BioID data: |

| | | | |
|---|---|---|---|
| | | | • Rsp20 is biotinylated in cells expressing BirA*-tagged Rsp3B and Rsp4A |
| Rsp22/LC8 | TTHERM_000649439 | RS1<br>RS2 | Comparative global ciliome analyses:<br>• The level of Rsp22 is diminished in *RSP3B* and *RSP3C* knockouts<br>Co-IP and BioID data:<br>• Rsp22 co-precipitates with Rsp3A and Rsp4A<br>• Rsp22 is biotinylated in cells expressing BirA* tagged either Rsp3,<br>• Not biotinylated in cells expressing Cfap61-HA-BirA*<br>Cross-link data:<br>• Rsp22 cross-links with Rsp3B, Rsp7A, Rsp14, and Rsp15 |
| Rsp23/Cfap67A | TTHERM_000372529 | RS1<br>RS2 | Co-IP and BioID data<br>• Rsp23 is biotinylated in cells expressing BirA* tagged either Rsp3B, Rsp4A<br>• Not biotinylated in cells expressing BirA*-tagged Cfap61 or Cfap91<br>• Co-precipitates with Rsp3A, Rsp4A, Rsp4B |
| Cfap61 | TTHERM_00641200 | RS3 | Cryo-ET analyses:<br>– fragment of RS3 spoke is missing in CFAP61-KO cells (Urbanska et al., 2015) |
| Cfap251 | TTHERM_01262850 | RS3 | Cryo-ET analyses:<br>• fragment of the an arch-like structure at the RS3 base is missing in CFAP251-KO cells (Urbanska et al., 2015) |
| Cfap91 | TTHERM_00578560 | RS3 | TEM analyses:<br>of CFAP91-KO mutant showing a lack of some RS3 (Bicka et al., 2022) |
| Cfap206 | TTHERM_00820660 | RS2 | Cryo-ET analyses:<br>– either entire RS2 or is base and in some cases also RS3 base is missing in CFAP206-KO cells (Vasudevan et al., 2015) |
| Cfap253 | TTHERM_00316930 | RS1 | Co-IP and BioID data:<br>• Cfap253 co-precipitates with Rsp3A<br>• Cfap253 is biotinylated in cells expressing Rsp3A-HA-BirA* and Rsp3B-HA-BirA*<br>Cross-link data:<br>• Cfap253 cross-links with Rsp3B |
| Cfap207 | TTHERM_00529880 | RS2 | Comparative global ciliome analyses:<br>• The level of Cfap207 is reduced in *RSP3B, CFAP206*, and *CFAP91* knockouts |
| Cfap198A | TTHERM_01092470 | RS1, RS2 (?) | Comparative global ciliome analyses:<br>• Level diminished in *RSP3A* and *RSP3B* knockouts |
| Cfap198B | TTHERM_00476710 | Rsp3C-containing RS2 subtype | Comparative global ciliome analyses:<br>• Level diminished in *RSP3C* |
| TtTpr | TTHERM_00623040 | RS2 | Comparative global ciliome analyses:<br>• The level of TtTpr is reduced in *RSP3C* and *CFAP206* knockouts<br>Co-IP and BioID data:<br>• TtTpr is biotinylated in cells expressing Cfap206-HA-BirA*<br>Cross-link data:<br>• TtTpr cross-links with Ak8A assigned to RS2 |
| Kelch-motif protein | TTHERM_00760390 | RS3 | Comparative global ciliome analyses:<br>• The level of TtKelch is reduced in *CFAP91* knockout<br>Cross-link data:<br>• Kelch-motif proteins cross-links with Cfap91 |
| Lrrc23A | TTHERM_000703669 | RS3 | Comparative global ciliome analyses:<br>• The level of Lrrc23A is reduced in *CFAP91* knockout<br>BioID data:<br>• Lrrc23A-HA-BirA* biotinylates primary RS3 proteins Cfap61 and Cfap251 |
| Lrrc23B | TTHERM_00105260 | RS3 | Comparative global ciliome analyses:<br>• The level of Lrrc23B is reduced in *CFAP91* knockout<br>BioID data:<br>• Lrrc23B-HA-BirA* biotinylates primary RS3 proteins Cfap61 and Cfap251 |
| Lrrc | TTHERM_01084360 | RS3 vicinity? | Comparative global ciliome analyses:<br>• The level of Lrrc is reduced in *RSP3B* and *CFAP91* knockouts |
| Lrrc | TTHERM_00046820 | RS3 base? | Comparative global ciliome analyses:<br>• The level of Lrrc is reduced in *CFAP61* and *CFAP91* knockouts<br>Cross-link data:<br>• protein cross-links with Ak7A, Ak9, and DYH24 |
| RIIa domain | TTHERM_00537370 | RS3? | Comparative global ciliome analyses:<br>• The level of RIIa domain is reduced in *CFAP91* knockout<br>Co-IP and BioID data:<br>• Protein is biotinylated in cells expressing BirA*-tagged, Lrrc23A, Lrrc23B |
| MRNN04 | TTHERM_00324550 | RS3 | Comparative global ciliome analyses:<br>• The level of MRNN04 is reduced in *CFAP91* knockout<br>Co-IP and BioID data:<br>• MRNN04 is biotinylated in cells expressing BirA*-tagged Cfap61, Cfap91, Lrrc23A, Lrrc23B |
| Stpg1 | TTHERM_00549640 | RS1<br>RS2 | Comparative global ciliome analyses: |

| | | | |
|---|---|---|---|
| | | | • The level of Stpg1 is reduced in *RSP3A, RSP3B, CFAP206,* and *CFAP91* knockouts |
| Stpg2 | TTHERM_00420710 | RS2 | Comparative global ciliome analyses:<br>• The level of Stpg2 is reduced in *RSP3B* and *CFAP206* knockouts |
| adenylate kinase 1, Ak1 | TTHERM_00317200 | RS1<br>RS2 | Comparative global ciliome analyses:<br>• The level of AK1 is reduced in *RSP3B* and *CFAP91* knockouts<br>Co-IP and BioID data:<br>• AK1 co-precipitates with Rsp3A, Rsp4A, Rsp4B, and Rsp4C<br>• AK1 is biotinylated in cells expressing BirA*-tagged Rsp3 and Rsp4 paralogs<br>Cross-link data:<br>• AK1 cross-links with Rsp4B, Rsp4C, and Rsp10 |
| adenylate kinase 7A, Ak7A | TTHERM_00558060 | RS3 | Comparative global ciliome analyses:<br>• The level of AK7A is reduced in *CFAP91* knockout<br>Cross-link data:<br>• AK7A cross-links with AK9 |
| adenylate kinase 7B, Ak7B | TTHERM_000569069 | RS3 | Comparative global ciliome analyses:<br>• The level of AK7B is reduced in *CFAP91* knockout<br>Cross-link data:<br>• AK7B cross-links with AK9 |
| adenylate kinase 8A, Ak8A | TTHERM_00227800 | R2 | Comparative global ciliome analyses:<br>• The level of AK8A is reduced in *RSP3B* knockout<br>Cross-link data:<br>• AK8A cross-links with Rsp3B, Rsp16A, TtTpr |
| adenylate kinase 8B, Ak8B | TTHERM_00455540 | RS1<br>RS2 | Comparative global ciliome analyses:<br>• The level of AK8B is reduced in *RSP3A* knockout<br>Co-IP and BioID data:<br>• AK8B is biotinylated in cells expressing BirA*-tagged Rsp3A and Rsp3B<br>Cross-link data:<br>• AK8B cross-links with Rsp16A |
| adenylate kinase 9, Ak9 | TTHERM_00148750 | RS3 | Comparative global ciliome analyses:<br>• The level of AK9 is reduced in *CFAP91* knockout<br>Cross-link data:<br>• AK9 cross-links with AK7A and AK7B |
| caseine kinase1, Ck1, | TTHERM_00938880 | RS3 | Comparative global ciliome analyses:<br>• The level of CK1 is reduced in *CFAP61* and *CFAP91* knockout |
| Guanylate kinase, Gk1 | TTHERM_00781030 | RS3 | Comparative global ciliome analyses:<br>• The level of Gk1 is reduced in *CFAP91* knockout<br>Co-IP and BioID data:<br>• GK1 is biotinylated in cells expressing BirA*-tagged Cfap61 and Cfap91 |
| Phosphodiesterase, Pde | TTHERM_00293350 | RS3 | Comparative global ciliome analyses:<br>• The level of Pde is reduced in *CFAP61* and *CFAP91* knockouts<br>Co-IP and BioID data:<br>• PdeB is biotinylated in cells expressing BirA*-tagged Lrrc23B |
| PKA catalytic subunit-1 | TTHERM_00433420 | RS2? | Comparative global ciliome analyses:<br>• The level of PKA catalytic sub-1 is reduced in *RSP3B, RSP3C,* and *CFAP91* knockout<br>Cross-link data:<br>• PKA catalytic sub-1 cross-links with PKA catalytic sub-1 and PKA regulatory sub |
| PKA catalytic subunit-2 | TTHERM_00658860 | RS2 | Comparative global ciliome analyses:<br>• The level of PKA catalytic sub-2 is reduced in *RSP3B* knockout<br>Cross-link data:<br>• PKA catalytic sub-2 cross-links with PKA catalytic sub-2, PKA regulatory sub, and Rsp15 |
| PKA regulatory protein | TTHERM_00623090 | RS2 | Comparative global ciliome analyses:<br>• The level of PKA regulatory sub is reduced in *RSP3B, RSP3C,* and *CFAP91* knockout<br>Cross-link data:<br>• PKA regulatory sub cross-links with PKA catalytic sub-1 and -2, and AK8B |

**Table S8.** Comparative analyses of the levels of the RS proteins in wild-type cells and RS mutants

| Radial spoke proteins | Acc numer in TGD | RSP3A-KO/WT | | | | RSP3B-KO/WT | | | | RSP3C-KO/WT | | | | CFAP206-KO/WT | | CFAP61-KO/WT | | CFAP91-KO/WT | | | |
|---|---|---|---|---|---|---|---|---|---|---|---|---|---|---|---|---|---|---|---|---|---|
| | | LFQ | | TMT | | LFQ | | TMT | | LFQ | | TMT | | LFQ | | LFQ | | LFQ | | TMT | |
| | | q-value | diffe-rence | q-value | diffe-rence | q-value | diffe-rence | q-value | diffe-rence | q-value | diffe-rence | q-value | diffe-rence | q-value | diffe-rence | q-value | diffe-rence | q-value | diffe-rence | q-value | diffe-rence |
| Rsp1 | TTHERM_000196029 | 0.7795 | -0.15 | 0.2168 | -0.13 | 0.0171 | -1.54 | 0.0016 | -0.64 | 0.8424 | -0.12 | 0.5512 | -0.07 | 0.3602 | -0.37 | 0.9789 | -0.01 | 0.6253 | 0.21 | 0.1372 | -0.13 |
| Rsp2 | TTHERM_00394410 | 0.4621 | -0.50 | 0.0414 | -0.29 | 0.0196 | -1.54 | 0.0012 | -0.59 | 0.3111 | -0.74 | 0.3073 | -0.13 | 0.3215 | -0.42 | 0.4980 | 0.24 | 0.6309 | -0.21 | 0.5120 | -0.05 |
| Rsp3A | TTHERM_01044600 | 0.0025 | -5.23 | 0.0124 | -0.99 | 0.0748 | -1.12 | 0.0029 | -0.40 | 0.7104 | -0.22 | 0.6405 | 0.05 | 0.3774 | -0.41 | 0.9426 | 0.02 | 0.8836 | -0.08 | 0.0548 | 0.18 |
| Rsp3B | TTHERM_00566810 | 0.4992 | 0.41 | 0.3772 | -0.09 | 0.0010 | -5.10 | 0.0019 | -1.15 | 0.6048 | 0.30 | 0.8883 | 0.02 | 0.0768 | -1.05 | 0.5058 | -0.24 | 0.0040 | -2.61 | 0.0041 | -0.43 |
| Rsp3C | TTHERM_00418270 | 0.9072 | -0.06 | 0.8769 | 0.02 | 0.2197 | -0.53 | 0.1760 | -0.10 | 0.0009 | -5.47 | 0.0044 | -0.59 | 0.0085 | -3.39 | 0.3490 | 0.43 | 0.0554 | -1.69 | 0.0399 | -0.21 |
| Rsp4A | TTHERM_00427590 | 0.7013 | -0.20 | 0.3759 | -0.08 | 0.0320 | -1.32 | 0.0011 | -0.49 | 0.9359 | -0.05 | 0.4944 | 0.07 | 0.1033 | -0.83 | 0.7104 | 0.12 | 0.2889 | -0.40 | 0.2864 | -0.08 |
| Rsp4B | TTHERM_00502580 | 0.3863 | -0.80 | 0.9895 | 0.002 | 0.2212 | -0.75 | 0.9626 | 0.00 | 0.0082 | -4.58 | 0.0125 | -0.79 | 0.0954 | -0.94 | 0.9786 | 0.01 | 0.0484 | -2.05 | 0.0219 | -0.26 |
| Rsp4C | TTHERM_00444180 | 0.3744 | -0.74 | 0.1361 | -0.18 | 0.0003 | -3.62 | 0.0017 | -0.68 | 0.5952 | -0.27 | 0.5776 | 0.07 | 0.9839 | -0.01 | 0.9724 | 0.01 | 0.8082 | -0.11 | 0.6438 | -0.03 |
| Rsp7A | TTHERM_00670610 | 0.4480 | 0.61 | 0.0700 | -0.29 | 0.0441 | -2.04 | 0.0019 | -0.77 | 0.4556 | 0.57 | 0.8805 | 0.02 | 0.0963 | -1.34 | 0.8033 | -0.08 | 0.0248 | -1.16 | 0.0044 | -0.48 |
| Rsp7B | TTHERM_00492840 | 0.5596 | -0.30 | 0.0471 | 0.25 | 0.7444 | -0.16 | 0.0031 | 0.35 | 0.6300 | -0.27 | 0.0271 | 0.36 | 0.0815 | -3.00 | 0.3705 | -0.42 | 0.0043 | -5.51 | 0.0020 | -0.51 |
| Rsp8 | TTHERM_00313520 | 0.0444 | -2.31 | 0.2889 | -0.11 | 0.0180 | -4.38 | 0.0006 | -1.34 | 0.0040 | -3.86 | 0.0017 | -0.80 | 0.0247 | -2.51 | 0.0381 | 1.12 | 0.0056 | -3.19 | 0.0006 | -1.21 |
| Rsp9 | TTHERM_00430020 | 0.7701 | -0.15 | 0.4773 | -0.08 | 0.0190 | -1.59 | 0.0024 | -0.40 | 0.6139 | -0.27 | 0.1883 | 0.16 | 0.4841 | -0.30 | 0.3612 | -0.37 | 0.1901 | -0.57 | 0.5680 | 0.04 |
| Rsp10 | TTHERM_00378600 | 0.3694 | -0.78 | 0.4403 | -0.08 | 0.0148 | -2.16 | 0.0012 | -0.58 | 0.3030 | -0.79 | 0.7289 | -0.04 | 0.5293 | -0.23 | 0.7611 | 0.10 | 0.6808 | -0.18 | 0.2148 | -0.11 |
| Rsp11 | TTHERM_00540090 | 0.2115 | -1.53 | 0.0307 | -0.31 | 0.0111 | -4.64 | 0.0011 | -0.49 | 0.0869 | -2.04 | 0.0367 | -0.35 | 0.5919 | 1.14 | 0.1085 | 2.76 | 0.6309 | 1.01 | 0.0548 | -0.20 |
| Rsp12A | TTHERM_01018330 | 0.9256 | 0.05 | 0.2392 | 0.11 | 0.0161 | -2.27 | 0.0079 | -0.29 | 0.0010 | -4.46 | 0.0118 | -0.45 | 0.0096 | -2.20 | 0.5560 | -0.26 | 0.0036 | -3.28 | 0.0019 | -0.53 |
| Rsp12B | TTHERM_00466090 | nd | | 0.0102 | -0.48 | nd | | 0.0000 | -0.68 | nd | | 0.0807 | 0.24 | nd | | nd | | nd | | 0.5789 | 0.04 |
| Rsp14 | TTHERM_000773538 | 0.0377 | -2.51 | 0.0045 | -0.74 | 0.0146 | -2.40 | 0.0010 | -0.52 | 0.3733 | -0.65 | 0.8355 | 0.02 | 0.6305 | -0.24 | 0.1018 | 0.72 | 0.0130 | 1.08 | 0.0172 | 0.31 |
| Rsp15 | TTHERM_00695730 | 0.2194 | -1.21 | 0.5601 | -0.06 | 0.0120 | -3.67 | 0.0019 | -0.86 | 0.1759 | -1.49 | 0.2321 | -0.17 | 0.0080 | -2.91 | 0.3356 | -0.88 | 0.0042 | -5.13 | 0.0038 | -0.97 |
| Rsp16A | TTHERM_00471260 | 0.0406 | -2.62 | 0.0253 | -0.33 | 0.0216 | -2.25 | 0.0013 | -0.56 | 0.0968 | -1.59 | 0.5844 | -0.07 | 0.3365 | -0.51 | 0.2292 | 0.54 | 0.4597 | 0.35 | 0.4636 | 0.05 |
| Rsp16B | TTHERM_00238810 | 0.8796 | -0.08 | 0.3494 | 0.10 | 0.3395 | -0.40 | 0.6381 | 0.03 | 0.0121 | -5.13 | 0.0064 | -0.67 | 0.0092 | -1.92 | 0.8685 | -0.05 | 0.0078 | -1.24 | 0.0354 | -0.22 |
| Rsp20/CaM1 | TTHERM_00630500 | nd | | 0.1719 | -0.20 | nd | | 0.0240 | -0.32 | nd | | 0.0117 | -0.67 | 0.1659 | -2.02 | 0.0406 | 1.00 | 0.0049 | -3.10 | 0.3203 | 0.10 |
| Rsp22/Lc8 | TTHERM_000649439 | 0.0666 | -2.53 | 0.4513 | -0.08 | 0.0439 | -1.62 | 0.0027 | -0.42 | 0.0461 | -3.36 | 0.2872 | 0.11 | 0.0952 | -1.00 | 0.2773 | 0.5 | 0.4440 | -0.38 | 0.2457 | 0.08 |
| Rsp23/Cfap67A | TTHERM_000372529 | 0.6087 | -0.29 | 0.9553 | -0.01 | 0.7482 | -0.16 | 0.6381 | 0.03 | 0.6874 | -0.26 | 0.6799 | 0.04 | 0.5172 | -0.25 | 0.5812 | 0.19 | 0.9600 | 0.03 | 0.0188 | 0.27 |
| Cfap61 | TTHERM_00641200 | 0.5334 | -0.39 | 0.0010 | 1.32 | 0.5872 | -0.23 | 0.0002 | 1.41 | 0.5468 | -0.36 | 0.0044 | 1.46 | 0.6809 | -0.16 | 0.0128 | -3.60 | 0.0044 | -3.56 | 0.1182 | 0.13 |
| Cfap251 | TTHERM_01262850 | 0.7834 | -0.15 | 0.3688 | 0.09 | 0.6187 | -0.23 | 0.4797 | 0.04 | 0.9708 | -0.02 | 0.0437 | 0.28 | 0.3754 | -0.34 | 0.9176 | -0.03 | 0.0049 | -5.61 | 0.0095 | -0.88 |
| Cfap91 | TTHERM_00578560 | 0.6777 | -0.21 | 0.3425 | 0.10 | 0.2540 | -0.54 | 0.1045 | -0.13 | 0.5640 | -0.34 | 0.2188 | 0.14 | 0.3221 | -0.40 | 0.7473 | -0.10 | 0.0035 | -5.85 | 0.0089 | -0.97 |
| Cfap206 | TTHERM_00820660 | 0.8793 | -0.08 | 0.4950 | 0.07 | 0.0008 | -4.04 | 0.0074 | -1.04 | 0.6660 | -0.22 | 0.3175 | -0.16 | 0.0139 | -4.67 | 0.2723 | -0.46 | 0.0041 | -4.55 | 0.0083 | -1.37 |
| Cfap253 | TTHERM_00316930 | 0.6415 | -0.24 | 0.3314 | -0.09 | 0.8457 | -0.09 | 0.1084 | 0.12 | 0.5416 | -0.36 | 0.4070 | 0.09 | 0.5152 | -0.45 | 0.3267 | 0.55 | 0.6564 | 0.27 | 0.0557 | 0.18 |
| Cfap207 | TTHERM_00529880 | 0.8223 | -0.12 | 0.5520 | 0.06 | 0.0137 | -3.31 | 0.0040 | -0.90 | 0.5665 | -0.32 | 0.9967 | 0.00 | 0.0157 | -3.76 | 0.4082 | -0.29 | 0.0177 | -3.93 | 0.0024 | -1.08 |
| Cfap198A | TTHERM_01092470 | 0.8513 | -0.09 | 0.0085 | -0.55 | 0.4064 | -0.34 | 0.0004 | -0.98 | 0.0889 | -2.42 | 0.0710 | 0.25 | 0.3138 | -0.76 | 0.2017 | -1.37 | 0.5137 | 0.84 | 0.0423 | 0.20 |
| Cfap198B | TTHERM_00476710 | 0.8513 | -0.09 | 0.0927 | 0.23 | 0.4064 | -0.34 | 0.4849 | 0.05 | 0.0889 | -2.42 | 0.0177 | -0.45 | 0.1449 | -1.85 | 0.2997 | -1.01 | 0.0467 | -1.51 | 0.1430 | -0.14 |

Significant reduction or increase of the protein level are marked in red or blue, respectively; TGD – Tetrahymena Genome Database; LFQ – label-free quantification, TMT- tandem mass tag 10-plex isobaric mass tagging approaches.

**Table S9.** Central apparatus components identified among proteins biotinylated in cilia of cells expressing BirA*-tagged Rsp**.**

| TGD number | Protein name | CA projection | BirA*-tagged protein | | | | | | | | | | | | |
|---|---|---|---|---|---|---|---|---|---|---|---|---|---|---|---|
| | | | Rsp3A | Rsp3A | Rsp3A | Rsp3B | Rsp3B | Rsp3B | Rsp3C | Rsp4A | Rsp4A | Rsp4C | Lrrc23A* | Lrrc23B* | Lrrc23B |
| TTHERM_00430030 | Pf6 | C1a | | | | | | 1/1 | | | 1/1 | | 25/18 | 35/24 | |
| TTHERM_00924250 | Ccdc180/Cfap76 | C1c/d | | | | | | | | | | | 17/14 | 14/12 | |
| TTHERM_00705200 | Cfap46 | C1d | 25/24 | 18/9 | 21/11 | 1/1 | | | 4/4 | 29/28 | | 55/39 | 59/38 | 140/60 | 17/11 |
| TTHERM_00049190 | Cfap54 | C1d | 4/4 | | | | | | 4/2 | 7/7 | | 29/22 | 139/81 | 216/107 | 19/12 |
| TTHERM_00530270 | Cfap74 | C1d | | | | | | | | | | 1/1 | 18/14 | 23/15 | |
| TTHERM_00189530 | Cfap221 | C1d | | | | | | | 1/1 | | | 3/3 | 18/9 | 25/14 | 1/1 |
| TTHERM_01142770 | Spef2A C1b | C1b | | | | | | | | | 2/2 | | 15/13 | 17/15 | 1/1 |
| TTHERM_00205170 | Tt170 C1b | C1b | 1/1 | | 1/1 | | | | | | | | 2/2 | 6/4 | |
| TTHERM_00290850 | Androglobin | C1b | | | | | | | | | | | 48/29 | 28/20 | 1/1 |
| TTHERM_00497260 | Cfap70 | C2a | | | | | | | | | | | 10/8 | 15/10 | |
| TTHERM_00551040 | Hydin | C2b | 3/3 | | | | | | | 1/1 | | 9/9 | 200/112 | 268/122 | 5/5 |
| TTHERM_000495990 | Cfap47 | C2b | | | | | | | | | | | 263/107 | 91/56 | |

*TurboID

**Table S10.** Comparative analyses of the levels of potential RS-associated proteins in wild-type cells and RS mutants

| Protein name | Acc number in TGD | RSP3A-KO/WT | | | | RSP3B-KO/WT | | | | RSP3C-KO/WT | | | | CFAP206-KO/WT | | CFAP61-KO/WT | | CFAP91-KO/WT | | | |
|---|---|---|---|---|---|---|---|---|---|---|---|---|---|---|---|---|---|---|---|---|---|
| | | LFQ | | TMT | | LFQ | | TMT | | LFQ | | TMT | | LFQ | | LFQ | | LFQ | | TMT | |
| | | q-value | diffe-rence | q-value | diffe-rence | q-value | diffe-rence | q-value | diffe-rence | q-value | diffe-rence | q-value | diffe-rence | q-value | diffe-rence | q-value | diffe-rence | q-value | diffe-rence | q-value | diffe-rence |
| TtTpr | TTHERM_00623040 | 0.4441 | -0.77 | 0.6006 | -0.05 | 0.1834 | -0.66 | 0.9062 | -0.01 | 0.0091 | -3.42 | 0.0116 | -0.56 | 0.0256 | -2.01 | 0.7405 | 0.14 | 0.0683 | -1.95 | 0.1189 | -0.14 |
| kelch-repeat | TTHERM_00760390 | 0.9112 | 0.06 | 0.8595 | 0.02 | 0.8452 | -0.09 | 0.1661 | 0.11 | 0.9543 | -0.04 | 0.2436 | 0.14 | 0.6226 | -0.23 | 0.5444 | 0.23 | 0.0053 | -3.87 | 0.0017 | -0.52 |
| Lrrc23A | TTHERM_000703669 | 0.2089 | -1.02 | 0.0663 | 0.23 | 0.3478 | -0.45 | 0.0339 | 0.21 | 0.2615 | -1.73 | 0.0178 | 0.42 | 0.2243 | -1.41 | 0.7830 | -0.10 | 0.0035 | -4.12 | 0.0014 | -1.16 |
| Lrrc23B | TTHERM_00105260 | 0.7001 | -0.22 | 0.6874 | 0.04 | 0.3550 | -0.40 | 0.1415 | 0.12 | 0.4826 | -0.45 | 0.0131 | 0.46 | 0.1661 | -0.60 | 0.0477 | -0.94 | 0.0034 | -6.02 | 0.0003 | -1.12 |
| Lrrc | TTHERM_01084360 | 0.5992 | 0.27 | 0.0605 | 0.26 | 0.1355 | -0.95 | 0.0019 | -0.40 | 0.4602 | -0.45 | 0.0285 | 0.34 | 0.0682 | -2.21 | 0.0770 | -1.45 | 0.0085 | -1.27 | 0.0132 | -0.32 |
| Lrrc | TTHERM_00046820 | 0.5539 | -0.37 | 0.0696 | 0.25 | 0.2356 | -0.59 | 0.0646 | -0.16 | 0.4420 | -0.64 | 0.3744 | 0.11 | 0.077 | -1.64 | **0.0328** | **-2.53** | **0.0072** | **-2.11** | **0** | **-0.73** |
| RIIa domain | TTHERM_00537370 | 0.7366 | 0.19 | 0.3208 | -0.09 | 0.8479 | 0.12 | 0.1545 | -0.10 | 0.8104 | -0.16 | 0.1419 | 0.25 | 0.6208 | 0.55 | 0.0605 | 3.1 | 0.0838 | 1.24 | 0 | -0.8 |
| MRNN04 | TTHERM_00324550 | 0.5 | 2.84 | 0.0065 | 0.55 | 0.5868 | -0.27 | 0.0009 | 0.47 | 0.5294 | 0.44 | 0 | 0.71 | 0.3124 | -0.47 | 0.1816 | -0.61 | 0.003 | -4.92 | 0 | -0.65 |
| Stpg1 | TTHERM_00549640 | 0.8430 | 0.16 | 0.0267 | -0.36 | 0.0337 | -1.43 | 0.0170 | -0.29 | 0.9056 | 0.08 | 0.0044 | 0.81 | 0.0185 | -3.19 | 0.0546 | -1.59 | 0.0147 | -3.47 | 0.0354 | -0.90 |
| Stpg2 | TTHERM_00420710 | 0.5100 | 0.64 | 0.1000 | -0.20 | 0.1549 | -0.77 | 0.0028 | -0.38 | 0.3054 | 1.02 | 0.1326 | -0.18 | 0.1105 | -0.94 | 0.1492 | -0.64 | 0.0089 | -3.72 | 0.1069 | -0.14 |

Significant reduction or increase of the protein level are marked in red or blue, respectively; TGD – Tetrahymena Genome Database; LFQ – label-free quantification, TMT- tandem mass tag 10-plex isobaric mass tagging approaches.

**Table S11.** Comparative analyses of the levels of proteins with a putative enzymatic activities in RS mutants

| **Enzyme name** | Acc numer in TGD | RSP3A-KO/WT | | | | RSP3B-KO/WT | | | | RSP3C-KO/WT | | | | CFAP206-KO/WT | | CFAP61-KO/WT | | CFAP91-KO/WT | | | |
|---|---|---|---|---|---|---|---|---|---|---|---|---|---|---|---|---|---|---|---|---|---|
| | | LFQ | | TMT | | LFQ | | TMT | | LFQ | | TMT | | LFQ | | LFQ | | LFQ | | TMT | |
| | | q-value | diffe-rence | q-value | diffe-rence | q-value | diffe-rence | q-value | diffe-rence | q-value | diffe-rence | q-value | diffe-rence | q-value | diffe-rence | q-value | diffe-rence | q-value | diffe-rence | q-value | diffe-rence |
| adenylate kinase 1, Ak1 | TTHERM_00317200 | 0.2096 | -1.03 | 0.2215 | -0.12 | **0.0186** | **-1.50** | **0.0010** | **-0.49** | 0.2443 | -0.93 | 0.4282 | -0.09 | 0.3223 | -0.43 | 0.4288 | 0.29 | 0.2574 | -0.43 | **0.0346** | **-0.22** |
| adenylate kinase 7A, Ak7A | TTHERM_00558060 | 0.9677 | -0.02 | 0.0706 | 0.23 | 0.2455 | -0.51 | 0.6449 | 0.03 | 0.9148 | -0.06 | 0.0778 | 0.24 | 0.3148 | -0.43 | 0.2183 | -0.45 | **0.0095** | **-3.57** | **0.0089** | **-0.37** |
| adenylate kinase 7B, Ak7B | TTHERM_000569069 | 0.6181 | -0.27 | 0.0196 | 0.33 | 0.3412 | -0.42 | 0.0116 | 0.28 | 0.5422 | -0.36 | 0.0306 | 0.35 | 0.3047 | -0.42 | 0.1970 | -0.53 | **0.0048** | **-6.74** | **0.0000** | **-0.99** |
| adenylate kinase 8A, Ak8A | TTHERM_00227800 | 0.5095 | -0.41 | 0.4586 | 0.07 | **0.0181** | **-1.80** | **0.0111** | **-0.28** | 0.2103 | -1.10 | 0.6136 | -0.05 | 0.1451 | -0.69 | 0.1838 | 0.54 | 0.6095 | 0.22 | 0.1733 | -0.11 |
| adenylate kinase 8B, Ak8B | TTHERM_00455540 | **0.0003** | **-4.12** | **0.0065** | **-0.60** | 0.1562 | -0.73 | 0.0723 | -0.16 | 0.8016 | 0.14 | 0.2832 | 0.12 | 0.5638 | -0.25 | 0.5959 | 0.18 | 0.2885 | 0.41 | **0.0134** | **0.31** |
| adenylate kinase 9, Ak9 | TTHERM_00148750 | 0.4987 | -0.40 | 0.0318 | 0.30 | 0.2744 | -0.44 | 0.0031 | 0.35 | 0.4566 | -0.48 | 0.0129 | 0.45 | 0.4154 | -0.30 | 0.1851 | -0.49 | **0.0034** | **-5.47** | **0.0047** | **-0.76** |
| caseine kinase 1, Ck1, | TTHERM_00938880 | 0.3214 | -1.20 | 0.0153 | 0.43 | 0.1869 | -0.79 | 0.0009 | 0.48 | 0.3032 | -0.90 | 0.0172 | 0.47 | 0.3752 | -0.42 | **0.0398** | **-1.77** | **0.0040** | **-3.45** | **0.0215** | **-0.32** |
| Guanylate kinase, Gk1 | TTHERM_00781030 | 0.5793 | -0.30 | 0.8151 | 0.02 | 0.3717 | -0.41 | 0.024 | **0.23** | 0.5442 | -0.33 | 0.0658 | 0.24 | 0.4619 | -0.33 | 0.4631 | -0.31 | **0.0047** | **-4.83** | **0** | **-1.36** |
| Phosphodiesterase, Pde | TTHERM_00293350 | 0.4821 | -0.49 | 0.0100 | 0.53 | 0.3325 | -0.43 | 0.001 | 0.55 | 0.5061 | -0.48 | **0** | **0.75** | 0.7072 | -0.18 | **0.0106** | **-4.61** | **0.005** | **-.5.49** | **0** | **-0.65** |
| PKA catalytic subunit | TTHERM_00433420 | 0.7840 | 0.21 | 0.6428 | -0.04 | 0.0831 | -1.36 | **0.0000** | **-0.98** | 0.5403 | -0.59 | **0.0000** | **-0.61** | 0.9273 | -0.05 | 0.2436 | 0.49 | 0.0206 | -1.50 | **0.0037** | **-0.38** |
| PKA catalytic subunit | TTHERM_00658860 | 0.5455 | -0.68 | 0.1637 | -0.16 | 0.1845 | -1.16 | **0.0019** | **-0.63** | 0.3813 | -1.09 | 0.1141 | -0.23 | 0.2830 | 0.55 | 0.3487 | 0.43 | 0.0075 | -3.58 | 0.7250 | 0.03 |
| PKA regulatory subunit | TTHERM_00623090 | 0.8422 | 0.18 | 0.3634 | -0.09 | **0.0103** | **-3.57** | **0.0035** | **-1.04** | 0.8085 | 0.19 | **0.0120** | **-0.47** | 0.0945 | -0.96 | 0.6476 | -0.16 | 0.0073 | -3.93 | **0.0186** | **-0.27** |

Significant reduction or increase of the protein level are marked in red or blue, respectively; TGD – Tetrahymena Genome Database; LFQ – label-free quantification, TMT- tandem mass tag 10-plex

**Table S12.** Comparative analyses of the levels of inner dynein arms (IDAs) components in *RSP3* mutants

| Protein name | Numer in TGD | RSP3A-KO/WT | | | | RSP3B-KO/WT | | | | RSP3C-KO/WT | | | |
|---|---|---|---|---|---|---|---|---|---|---|---|---|---|
| | | LFQ | | TMT | | LFQ | | TMT | | LFQ | | TMT | |
| | | q-value | difference | q-value | difference | q-value | difference | q-value | difference | q-value | difference | q-value | difference |
| DYH6 | TTHERM_00688470 | 0,427619 | -0,63 | 0,97248 | 0,00 | 0,829776 | -0,11 | 0,0333644 | 0,19 | 0,42251 | -0,58 | 0,22074 | 0,14 |
| DYH7 | TTHERM_00912290 | 0,63228 | -0,26 | 0,92333 | 0,01 | 0,889118 | 0,06 | 0,0436144 | 0,17 | 0,552849 | -0,36 | 0,25214 | 0,14 |
| DYH8 | TTHERM_00531870 | nd | | 0,35423 | -0,09 | nd | | 0,794792 | 0,02 | nd | | 0,05891 | 0,27 |
| DYH9 | TTHERM_00947430 | 0,212183 | -2,35 | **0,01518** | **-0,47** | 0,19376 | -1,15 | 0,138485 | -0,11 | 0,202845 | -2,54 | 0,13222 | -0,19 |
| DYH10 | TTHERM_00420340 | nd | | 0,24242 | -0,12 | nd | | **0,0211327** | **-0,26** | nd | | 0,78056 | -0,03 |
| DYH11 | TTHERM_00252430 | 0,278289 | -1,14 | 0,64207 | 0,04 | 0,079717 | -1,13 | 0,695276 | -0,02 | 0,207409 | -1,33 | 0,61464 | -0,05 |
| DYH12 | TTHERM_00919540 | 0,391149 | -0,81 | 0,77556 | -0,03 | **0,016857** | **-3,02** | **0,0024172** | **-0,39** | 0,216267 | -1,14 | 0,89791 | -0,01 |
| DYH14 | TTHERM_00492830 | 0,627178 | -0,26 | 0,86334 | -0,02 | 0,81443 | 0,11 | 0,0682636 | 0,15 | 0,604414 | -0,28 | 0,36991 | 0,10 |
| DYH15 | TTHERM_00433800 | 0,509668 | -0,39 | 0,53573 | 0,06 | 0,98351 | -0,01 | 0,035494 | 0,19 | 0,58223 | -0,33 | 0,13251 | 0,19 |
| DYH16 | TTHERM_00558640 | 0,500578 | -0,41 | 0,49759 | 0,06 | 0,663928 | -0,19 | 0,0255092 | 0,21 | 0,478641 | -0,44 | 0,13177 | 0,20 |
| DYH17 | TTHERM_00850620 | nd | | 0,16265 | -0,15 | nd | | 0,322829 | -0,06 | nd | | 0,56919 | 0,06 |
| DYH18 | TTHERM_00047540 | nd | | nd | | nd | | nd | | nd | | nd | |
| DYH19 | TTHERM_01027670 | 0,59496 | -0,32 | 0,13983 | 0,16 | 0,650933 | -0,20 | 0,0160624 | 0,26 | 0,456762 | -0,57 | 0,1503 | 0,17 |
| DYH20 | TTHERM_00821980 | nd | | 0,27734 | -0,11 | nd | | 0,546651 | 0,04 | nd | | 0,04042 | 0,29 |
| DYH22 | TTHERM_00565600 | 0,739417 | -0,17 | 0,89428 | 0,01 | 0,80461 | -0,12 | 0,0433816 | 0,18 | 0,728791 | -0,18 | 0,63557 | 0,05 |
| DYH23 | TTHERM_00355100 | nd | | 0,568953 | -0,05 | nd | | 0,716902 | 0,02 | nd | | 0,36156 | 0,12 |
| DYH24 | TTHERM_00193520 | 0,454189 | -0,60 | 0,04083 | 0,26 | 0,0565 | -1,13 | 0,261682 | 0,07 | 0,306765 | -0,88 | 0,23358 | 0,15 |
| DYH25 | TTHERM_00774820 | 0,497702 | -0,46 | 0,29773 | 0,11 | **2,36E-05** | **-4,25** | **0,004094** | **-0,77** | 0,227139 | -1,28 | 0,19224 | -0,16 |
| p28A | TTHERM_00841210 | 0,62273 | 0,34 | 0,13922 | 0,17 | 0,140637 | -1,09 | 0,574056 | 0,04 | 0,529654 | -0,45 | 0,28815 | 0,11 |
| p28B | TTHERM_00319990 | 0,918872 | 0,05 | 0,71511 | 0,04 | **0,186438** | **-0,70** | **0,0378964** | **-0,20** | 0,546171 | -0,42 | 0,43061 | 0,11 |
| p28C | TTHERM_01129720 | 0,931915 | 0,05 | 0,8939 | 0,013 | 0,978215 | -0,016 | 0,0700761 | 0,16 | 0,478333 | -0,44 | 0,59091 | 0,07 |

Significant changes are marked in red; TGD – Tetrahymena Genome Database; LFQ – label-free quantification, TMT- tandem mass tag 10-plex isobaric mass tagging approaches; (nd, not detected) in analyzed ciliomes

**Table S13.** List of primers used in this study. The nucleotide sequences recognized by the restriction endonucleases are in bold.

| Primer name | Nucleotide sequence |
|---|---|
| **Native locus expression** (-3HA and/or –HA-BirA*) | |
| RSP3A-cod-MluI-F | AAAT**ACGCGT**AG GAGGAA CACAAT AGCAGT ATAA |
| RSP3A-cod-BamHI-R | AATT**GGATCC**CT CTGAAG GTAAAT TAAACC CAAAAT C |
| RSP3A-3UTR-PstI-F | TTAA**CTGCAG**CT TAAAGA ATAATG GATTTG TAGAGA TATTG |
| RSP3A-3UTR-XhoI-R | TTAA**CTCGAG**AT CATGTT TAAACT GATTTT AATAGT TATCAA A |
| RSP3B-cod-MluI-F | AAAT**ACGCGT**GT CTGAGG GCTCTA TTGATT TA |
| RSP3B-cod-BamHI-R | AATT**GGATCC**GT ATGATT TCATAT CTAAAA CAGGGA AA |
| RSP3B-3UTR-PstI-F | AAAT**CTGCAG**CA TATTTC TTTGCA AAATGT TTTATA AATTAG |
| RSP3B-3UTR-XhoI-R | AATT**CTCGAG**GT TGGATT TACAGC CAATCT AG |
| RSP3C-cod-MluI-F | AAAT**ACGCGT**CA ATAACC TCCAAG TGAAGA ATCA |
| RSP3C-cod-BamHI-R | AATT**GGATCC**AT CTTCTT CGGGTT CTTCCT CACCTT CTTCAC C |
| RSP3C-3UTR-PstI-F | AAAT**CTGCAG**CA TATTCT AATAAT CTTGAA TGCATG AATA |
| RSP3C-3UTR-XhoI-R | AATT**CTCGAG**AC TCTCTC CATCAC AAAAAT ATCTG |
| RSP4A-cod-MluI-F | AAAT**ACGCGT**CG TACTTT AATTGA CCCTGA TGAGA |
| RSP4A-cod-BamHI-R | AATT**GGATCC**AT TCTCTT CCTCTT CTTCTT ATTATT GCTATT GG |
| RSP4A-3UTR-PstI-F | AAAT**CTGCAG**CT ATCTAT CAATCG ATATATGTATGTATGGTT TC |
| RSP4A-3UTR-XhoI-R | AATT**CTCGAG**AA TATAAA AGTGCA AACGAA ATCATTAATTTTG |
| RSP4B-cod-MluI-F | AAAT**ACGCGT**GT GAAAAT GACATA TAACTT GAAGAT GAAC |
| RSP4B-cod-BamHI-R | AATT**GGATCC**AT TTTCTT CTTCAT CTTCTT CCTTCT TTT |
| RSP4B-3UTR-PstI-F | AAAT**CTGCAG**TTTATTGTCAATAATTAACTATCAATATCAAAAAG |
| RSP4B-3UTR-XhoI-R | ATAT**CTCGAG**GA ATTATT GATGAA GAAGAG TATGAGATTATGC |
| RSP4C-cod-MluI-F | ATTT**ACGCGT**GT ATTAAG CTTTGG ATGACC CTG |
| RSP4C-cod-BamHI-R | AATT**GGATCC**TTCTTATTCTTCTTAAGCATTCTCTT CATTTT CTTC |
| RSP4C-3UTR-PstI-F | AAAT**CTGCAG**GCTTTAAATACATATTAAATACATGTAATTATAAC |
| RSP4C-3UTR-XhoI-R | AATT**CTCGAG**AT ATCACT GTAAAC CTTGTA ATATTA CTAG |
| CFAP61-cod-MluI-F | AATT**ACGCGT**GA TGATAC CCAAAG TCGTGA TG |
| CFAP61-cod-BamHI-R | AATT**GGATCC**AT CTACAT TTACCT TCTTAG GAGGTA CATA |
| CFAP61-3UTR-PstI-F | AATT**CTGCAG**CA AACTAA ACATGT TACTCC TATTGT T |
| CFAP61-3UTR-XhoI-R | AATT**CTCGAG**CA TTCAAT ACTTAA CAGGAG AAATCT TC |
| CFAP91-cod-MluI-F | AATT**ACGCGT**GT CCTGCA ACTCCT ACTTG |
| CFAP91-cod-BamHI-R | AATT**GGATCC**AT TCTAAA CATTTG CGTGCT TATTTG |
| CFAP91-3UTR-PstI-F | AATT**CTGCAG**GA GGAGTA AGTAAC CAACAA ACC |
| CFAP91-3UTR-XhoI-R | AATT**CTCGAG**TC TCTAAT TCTAAA TTCCAG CTTTCA G |
| CFAP206-cod-MluI-F | AAAT**ACGCGT**GT TTATTG CTACCA GGTAAA CCT |
| CFAP206-cod-BamHI-R | AATT**GGATCC**AT TAGTGT CTTTGT CTCTTA ATCCAG T |
| CFAP206-3UTR-PstI-F | AAAT**CTGCAG**CCTTTCCAACTTTATATCAGATATAATTTATT ACC |
| CFAP206-3UTR-XhoI-R | AATT**CTCGAG**AAGCTTATTACAGTTTTAATAATTCCATCTTAGCC |
| LRRC23A-cod-MluI-F | AAAT**ACGCGT**TA TGAGTA AAAAAG AAAAGA AGCCAG |
| LRRC23A-cod-BamHI-R | AATT**GGATCC**AT TTTCTT CTTGCT ATTCTT ATTCTT GTTT |
| LRRC23A-3UTR-PstI-F | AAAT**CTGCAG**TA TTTACT GTAAAA TTTAGT CAGTAC ATCA |
| LRRC23A-3UTR-XhoI-R | AATT**CTCGAG**CT CGTTTT ACTTCT TGAATT TCTATA AATGCA T |
| LRRC23B-cod-MluI-F | AAAT**ACGCGT**TA TGTCAG AGTAGG AAATCG AAG |
| LRRC23B-cod-BamHI-R | AATT**GGATCC**TT CATCTT ACTGTT GCTAAT TTTAGG |
| LRRC23B-3UTR-PstI-F | AAAT**CTGCAG**AT ACATAA ATATGC AATGCA TTAAGT TATC |
| LRRC23B-3UTR-XhoI-R | AATT**CTCGAG**CT CTGGAT AAATTT ACTTCA ATTTGC TCAA |
| Gene knock-out | |
| RSP3A-KO-5-F-ApaI | AATT**GGGCCC**AA AGTATC AATGAA CACTTA ATTAGA ATAGC |

| | |
|---|---|
| RSP3A-KO-5-R-SmaI | AATT**CCCGGG**TG CTTATT CATATT CTTCAA TGAGCT C |
| RSP3A-KO-3-F-PstI | AATT**CTGCAG**AG CTCATT GTGAGT TTGCCG ATATTA CAG |
| RSP3A-KO-3-R-SacII | AATT**CCGCGG**CA ATATCT CTACAA ATCCAT TATTCT TTAAG |
| RSP3B-KO-5-F-ApaI | AATT**GGGCCC**CT AGTATT TAGTTA AACAAG GCATGC |
| RSP3B-KO-5-R-SmaI | AATT**CCCGGG**AC GAGGAT TAGGTC TGTCAA TAA |
| RSP3B-KO-3-F-PstI | AATT**CTGCAG**GC AGCAAG AATTCT AAGTAA AGCA |
| RSP3B-KO-3-R-SacII | AATT**CCGCGG**AG TTTTTG ATAAAG CTTCGT ATAATA TGG |
| RSP3C-KO-5-F-ApaI | AATT**GGGCCC**AC GCGTTA TGCAGT AAAGAA TTTTGTGTTTAG G |
| RSP3C-KO-5-R-SmaI | AATT**CCCGGG**AC TTATGT GTAAGC ATCAAT ACCAG |
| RSP3C-KO-3-F-PstI | AATT**CTGCAG**CA TATTAA CAGATT CCTTGA AGGAAC T |
| RSP3C-KO-3-R-SacII | AATT**CCGCGG**TT AGGAGG AATACC TTCAGT ACTG |
| Co-deletion primers | |
| RSP2-coDel-F | CAGTTC TCATCA AGTTGT AATGCT AAAATG CGGCCG CCTGCA GCTGTA TTTAAG GAGCC |
| RSP2-coDel-R | GGACTC TTTATT GTTATC ATCTTA TGACCG CGGCCG CCTGTT ATTCTG TTAACT AAGGCA TAAGG |
| Primers used to verify deletion of the fragment of RSP3 gene | |
| RSP3A-KO_check_R_2 | TGTTTA AAACCC AAGGCA AGAAAT T |
| RSP3A-KO_check_R_1 | TCCAGA TTTAAC TCAACA AACCTC CAA |
| RSP3A-KO_check_F_1 | GGATGA AAATGA GATAGA GCCAGA T |
| RSP3B-KO_check_R_2 | TGTTTC TTGATA TAAGTT TCTTGT GAGA |
| RSP3B-KO_check_R_1 | TAAGAC AAATTC TCCAAC AAGACC A |
| RSP3B-KO_check_F_1 | AATAAT TAACTG ACAAGC CACCTG A |
| RSP3C-KO_check_F_2 | AAAAGA TCGTCT TCTTGA CCTCTA T |
| RSP3C-KO_check_R_1 | CCTTGG AGAGCA TTATAC TTTGTG T |
| RSP3C-KO_check_F_1 | GGCTCC TTATGA TATTAA GCCATC T |
| RSP2-coDel-spr-F | CATAGA AGCAAA GACTGA TTCCAT |
| RSP2-coDel-spr-R | GACTTC TTAGAG AATATT ATGCAA ATTCG |

#### Supplementary Videos

Video 1. Cilia beating in wild-type *Tetrahymena* cell

Video 2. Cilia beating in RSP3A-KO *Tetrahymena* mutant

Video 3. Cilia beating in RSP3B-KO *Tetrahymena* mutant

Video 4. Cilia beating in RSP3C-KO *Tetrahymena* mutant
