## Supplementary figures and images for "Heterogeneity of radial spokes structural components and associated enzymes in *Tetrahymena* cilia"

### Video S1 -WT.gif

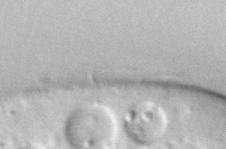

### Video S2-RSP3A.gif

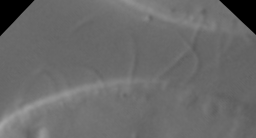

### Video S3a-RSP3B-KO.gif

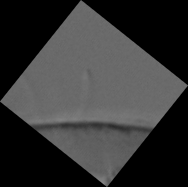

### Video S3b-RSP3B-KO.gif

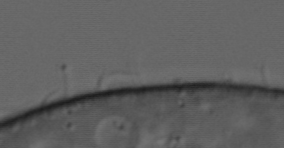

### Video S4-RSP3C-KO.gif

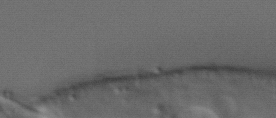
